# Population genomics of incipient allochronic divergence in the Pine Processionary Moth

**DOI:** 10.1101/2025.06.09.657100

**Authors:** Tanguy Muller, Mathieu Gautier, Éric Lombaert, Raphaël Leblois, Laure Sauné, Manuela Branco, Carole Kerdelhué, Charles Perrier

## Abstract

Allochronic divergence is a key evolutionary mechanism that can frequently lead to incipient speciation. Although theoretical models suggest that such divergence is notably facilitated by small population size and genetic polymorphisms influencing reproductive timing, though constrained by genetic load, empirical validation remains limited. We investigated these predictions by re-analyzing a case of allochronic differentiation between two sympatric populations of pine processionary moth (*Thaumetopoea pityocampa*) in Portugal, using whole genome resequencing (IndSeq and PoolSeq) of those two populations and eight allopatric ones. We inferred the demographic history of those populations, assessed their genetic load, and searched for genomic regions associated with life cycle differences. Our analyses revealed a recent split between the sympatric allochronic populations, accompanied by a strong reduction in gene flow, bottlenecks, inbreeding, and accumulation of deleterious variants. Genome scans identified several loci associated with life cycle variation, including genes putatively involved in circadian rhythm regulation, predominantly located on the Z chromosome. We discuss how these empirical genomic findings support theoretical expectations that assortative mating driven by differences in reproductive timing, underpinned by polymorphisms in circadian genes, along with genetic drift and purge of genetic load at high-impact sites, can promote the onset and persistence of allochronic divergence.

## Introduction

Differences in reproductive timing among individuals within a population can lead to the formation of reproductively isolated clusters that gradually accumulate genetic differences over time, a process called allochronic divergence, which may ultimately promote speciation (Coyne and Orr, 2004; Hendry and Day, 2005; Rundle and Nosil, 2005; Taylor and Friesen, 2017). Theoretical models developed by Devaux and Lande (2008) and Weis *et al*. (2014) suggest that, in sympatry, the formation of allochronic clusters of individuals may be promoted notably by genetic drift, the existence of genetic polymorphisms influencing reproductive time, positive assortative mating, and a short individual reproductive window within a longer reproductive season. Isolation by distance between allochronic clusters may also contribute to further reinforce allochronic divergence (Kunkel, 2023). In allopatry, in contrast, spatially divergent selection may be an important driver of allochronic divergence (Jarrett *et al*., 2025). The long-term viability of newly formed allochronic populations could be facilitated by adaptations to the new life cycle (Jarrett *et al*., 2025). In turn, it could be limited by inbreeding depression (Devaux and Lande, 2008), which can be exacerbated by the small population sizes often expected in such contexts. Although empirical investigations revealed the important role of genetic mutations at circadian genes underlying allochrony (Tessnow *et al*., 2022), thorough empirical analyses are needed to evaluate the respective roles of the several evolutionary processes promoting the onset of allochronic divergence.

Divergence in the timing of reproductive activity can stem from variation in genes regulating biological rhythms (Tomioka and Matsumoto, 2015; Hutfilz, 2022). In insects, circadian genes such as *period*, *pdfr*, *CYC*, *cry*, or *timeless* play key roles in controlling diapause duration and termination, thereby ensuring synchronization of reproductive activity with seasonal cycles (Abruzzi *et al*., 2017; Kozak *et al*., 2019; Brady *et al*., 2021). In parallel, genes involved in secondary sexual traits, pheromone production, olfaction, and assortative mating may further contribute to reinforcing reproductive isolation (Matessi *et al*., 2002; Unbehend *et al*., 2013; Kopp *et al*., 2018). Several genome-wide scans comparing allochronic populations have highlighted the involvement of polymorphisms in circadian genes in driving phenological divergence. For instance, Pruisscher *et al*. (2021), using whole-genome resequencing, identified non-synonymous SNPs differentially fixed in the *period* gene between two populations of *Pieris napi* differing in their diapause behavior. However, although such genetic polymorphisms at circadian genes may constitute the raw material for phenological differences, assortative mating and allochronic divergence, it may be difficult to decipher the relative roles of genetic drift, selection resulting from assortative mating, and spatially varying selection, as evolutionary forces driving genetic differentiation at these loci.

Inferring demographic histories is required to determine whether allochronic divergence between two populations emerged in sympatry *via* local strong genetic drift together with non-selective assortative mating based on genetically controlled phenology (Devaux and Lande, 2008; Kunkel, 2023) or was initially produced in allopatry via spatially divergent selection on genetic polymorphisms controlling phenology (Safran *et al*., 2016). Recent advances in statistical population genomics, such as those using simulation-based inference (e.g., Approximate Bayesian Computation or ABC, Raynal *et al*., 2019) allow the inference of demographic events, including temporal fluctuations in population size and gene flow. In addition, methods have recently been developed to measure the fraction of the genome within regions of autozygosity, as well as to track the temporal dynamics of accumulation of these regions within individual genomes (Druet and Gautier, 2017). These complementary approaches may help to reveal the putative role of demography and assortative mating in the onset of allochronic divergence.

While the particular context of small population size may promote allochronic divergence, it is also predicted to decrease genetic diversity and increase genetic load. Indeed, in general, reduced effective population size associated with bottlenecks and inbreeding can lead to the accumulation of genetic load, as purifying selection becomes less efficient in removing deleterious variants (Charlesworth, 2009; Woolfit, 2009; Lanfear *et al*., 2014). However, prolonged periods of small effective population size can also facilitate the purging of highly deleterious mutations, potentially mitigating some of the negative fitness consequences (Robinson *et al*., 2018, 2023). Hence, one could hypothesize that successfully established allochronic populations may have gone through strong bottlenecks and significant purging of the genetic load. Empirically examining whether genetic load is purged during allochronic divergence is crucial to assess the long-term viability and evolutionary trajectories of allochronically divergent populations.

The pine processionary moth (PPM, *Thaumetopoea pityocampa*, Lepidoptera, Notodontidae) provides a unique system to study allochronic divergence. Indeed, in the Mata Nacional de Leiria, Portugal, a recently discovered population (Pimentel *et al*., 2006; Santos *et al*., 2007, 2011) exhibits a clear shift in reproductive timing compared to the locally sympatric winter population (Figure 1 A,B). This so-called “Leiria summer population” (LSP) reproduces in spring, with summer larval development, whereas the sympatric “Leiria winter population” (LWP) follows the typical phenology of the species, with reproduction occurring in summer and larval development occurring in winter. Both populations hence experience strong pre-mating reproductive isolation (Santos *et al*., 2011; Burban *et al*., 2016). Experimental rearing suggested a high heritability of the reproductive period (Branco *et al*., 2017). Previous population genetic studies based on mitochondrial, microsatellite, and RAD-seq data (Santos *et al*., 2007, 2011; Burban *et al*., 2016; Leblois *et al*., 2018) showed low mitochondrial differentiation but high nuclear differentiation between LSP and LWP, suggesting strong bottlenecks and a recent split between these two populations. However, these studies were limited in their ability to elucidate the genomic mechanisms underlying the phenological shift, notably due to the lack of a chromosome-level genome assembly and the sparsity of genetic markers used.

**Figure 1:**
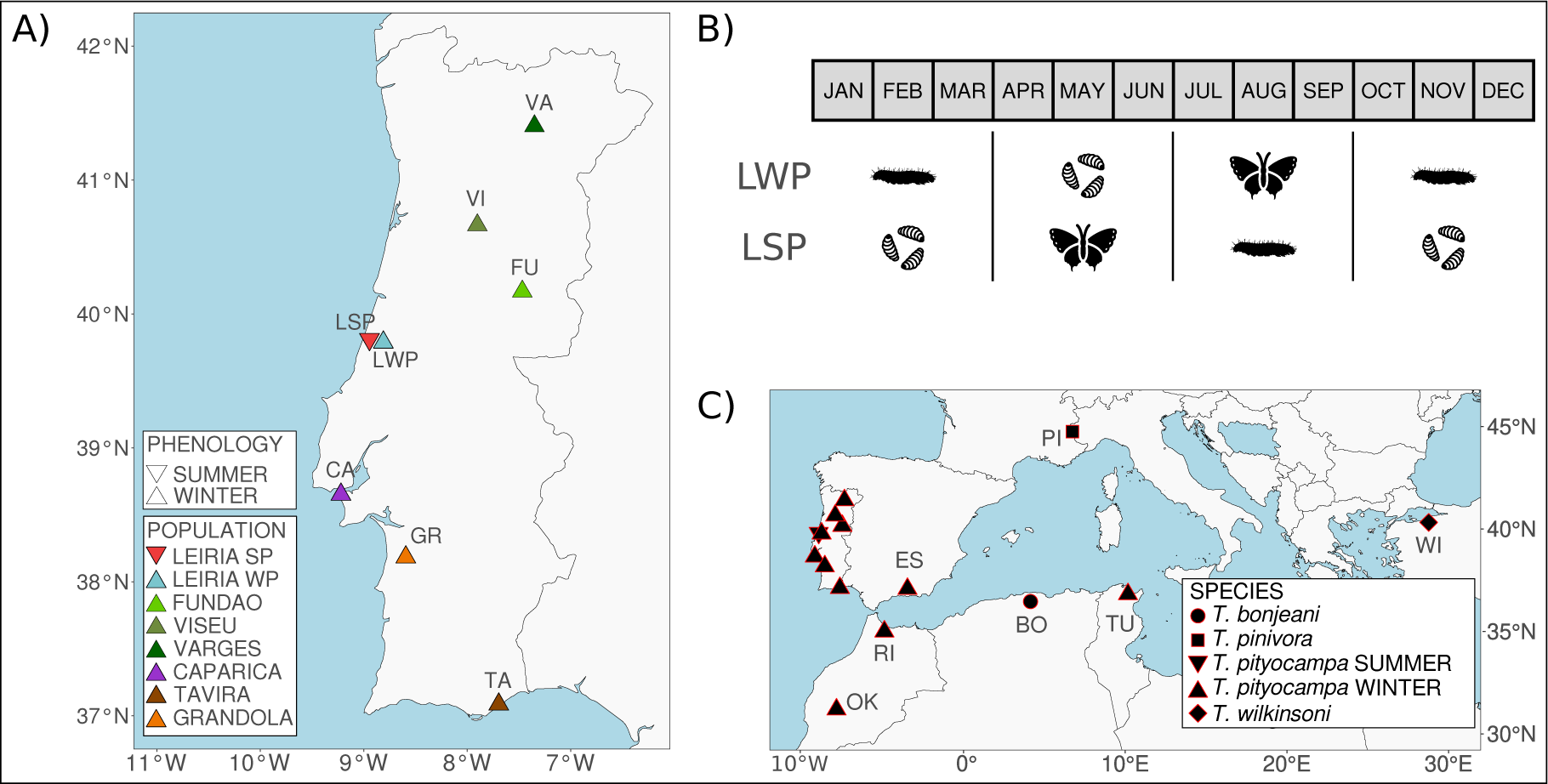
Geographical locations of all sampled populations and life cycle of *T. pityocampa*. **(A)** Map of the sampling locations for PPM populations over Portugal. Colors indicate the different Portuguese populations, and shapes their phenology. The population abbreviations stay the same throughout this study: LSP (Leiria Summer Population; n = 25), LWP (Leiria Winter Population; n = 18), FU (Fundão; n = 10), VI (Viseu; n = 4), VA (Varges; n = 4), CA (Caparica; n = 4), TA (Tavira; n = 4), GR (Grândola; n = 4). **(B)** Annual life cycle of the PPM, illustrating the LWP cycle (winter phenology) and the LSP cycle (summer phenology). Caterpillars represent the larval stage, pupae the pupal stage and moth the emergence period of adults. **(C)** Map of the sampling locations for all populations and species used in this study. Shapes indicate the sampled species. The population abbreviations stay the same throughout this study: ES (Sierra Nevada, Spain; n = 2), RI (Rif, Morocco; n = 2), OK (Oukaimeden, Morocco; n = 2), TU (Tunis, Tunisia; n = 2), BO (*T. bonjeani*; n = 2), PI (*T. pinivora*; n = 2), WI (*T. wilkinsoni;* n = 2).

In this study, we leverage recently developed genomic resources for the PPM, including a chromosome-level reference genome (Gautier *et al*., 2025) and whole genome resequencing data for both individuals (IndSeq) and pooled samples (PoolSeq) from different populations, to investigate the genetic basis and consequences of the PPM allochronic divergence. Specifically, our goals were (i) to reconstruct the recent demographic history of LSP and LWP, (ii) to investigate the genetic bases of phenological variation between these two populations, and (iii) to characterize and compare their respective genetic load. Following the previously aforementioned theoretical works (Devaux and Lande, 2008) and empirical studies on the PPM (Burban *et al*., 2016; Leblois *et al*., 2018), we hypothesized that recent bottlenecks and non-selective assortative mating relaying on genomic variations in circadian genes might have driven the recent onset of allochronic divergence of LSP and that genetic load might have been partly purged. By integrating demographic inferences, genome scans, and analyses of deleterious variation, we provide new empirical insights into the evolutionary mechanisms driving allochronic divergence and its genetic consequences.

## Methods

### Sampling

A total of 369 individuals of PPM were collected between 2002 and 2012 in the context of previous studies (Burban *et al*., 2016, 2020), in the summer (LSP) and winter (LWP) sympatric populations from the Leiria region (Figure 1A; Table 1; Table S1), as well as in more distant locations of Fundão (FU), Viseu (VI), Varges (VA), Caparica (CA), Tavira (TA), and Grândola (GR) in Portugal. For FU, CA, TA, and GR, adult individuals were captured using pheromone traps from mid-April to September. For VI and VA, larvae were collected from nests, with a single larva sampled per nest to avoid collecting siblings. For LSP and LWP, 12 and 6 larvae, and 13 and 12 adults, respectively, were collected following the same sampling strategy described above. All individuals were stored in 95% ethanol immediately after sampling and then kept at -18°C until DNA extraction. We also included 8 individuals referred to as “LateLSP”, sampled alongside LWP individuals (i.e. with a winter phenology) but genetically (at microsatellite and SNP markers) clustering with LSP individuals, and hypothetically resulting from plastic variations of phenology in LSP (Burban *et al*., 2020). We added 9 LSPxLWP hybrids (“F1”) produced by Branco *et al*. (2017) in laboratory. Lastly, 14 individuals were included as outgroups (Figure 1C; Table 1, Table S1): 2 individuals of *T. pityocampa* from Cortijuela (ES, Spain), Bab Barred (RI, Rif region, Morocco), Oukaimeden (OK, Morocco) and Tunis (TU, Tunisia), 2 individuals of *T. bonjeani* from Tikjda (BO, Algeria), 2 individuals of *T. pinivora* from Alpes (PI, France) and 2 individuals of *T. wilkinsoni* from Mudanya (WI, Türkiye).

**Table 1:**
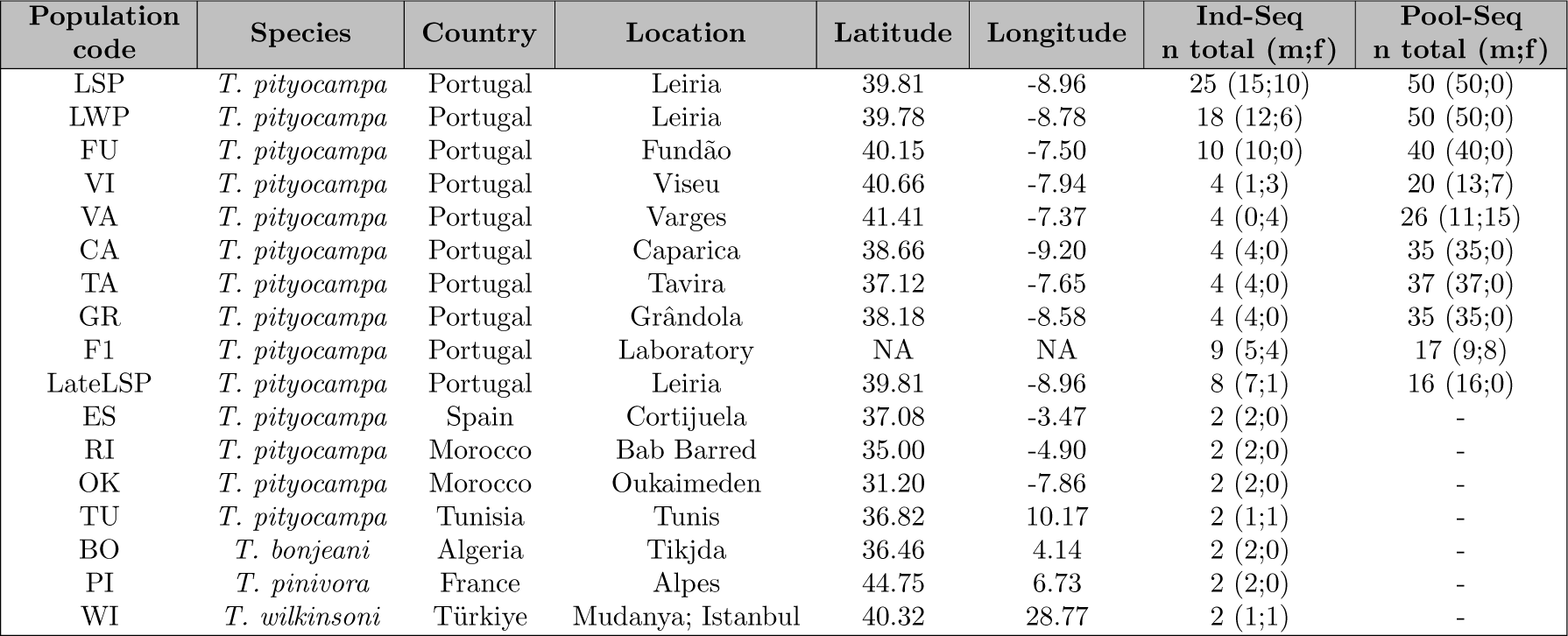
Study populations and resequencing strategy. For each studied population is given a code, species name, country of origin, latitude, longitude, number of resequenced individuals (Ind-Seq) for both sexes (m for males and f for females), and number of individuals in resequenced pools. For pools, the ratio of the Z chromosome to the chromosome 1 median coverage was used to estimate sex ratio (see text for details).

### DNA extraction, library preparation and sequencing

For each Portuguese population, the DNA of 4 to 25 individuals was used for individual resequencing, and pools of 16 to 50 individuals’ DNA were prepared for pool sequencing (Table 1). For each outgroup, the DNA of 2 individuals was used for individual resequencing (Table 1). All DNA samples, available from the aforementioned previous studies (Burban *et al*., 2016, 2020) were quantified on a Qubit fluorometer (Thermo Fisher Scientific), and their quality assessed on agarose gels. Whole genome libraries were constructed using the Illumina Truseq kit following the manufacturer’s recommendations. The quality of each library was inspected on a 2100 Bioanalyzer. The 114 libraries (10 Pool-Seq and 104 Ind-Seq) were paired-end (150 bp) sequenced at the MGX-Montpellier Genomix platform on an Illumina HiSeq2500 (4 libraries) or on a NovaSeq6000 (110 libraries).

### Variant calling

Raw paired-end reads were filtered using *fastp* (v0.21.0; Chen *et al*., 2018) with default parameters to remove contaminant adapter sequences and trim poor quality bases, with a phred-quality score *<* 15. Read pairs with either one read with a proportion of low-quality bases over 40% or containing more than 5 N bases were removed. The median percentage of reads filtered out per library was 1.52% (Figure S1A; Table S2). The median percentage of duplicated reads per library was 6.89% (Figure S1B; Table S2). Filtered reads were then mapped onto the *Tpit2.1* chromosome level reference genome (Gautier *et al*., 2025) using the *bwa-mem2* program (v2.2.1 Vasimuddin *et al*., 2019) with default options. Read alignments with a mapping quality Phred-score *<* 20 or PCR duplicates were removed using *SAMtools* software (v1.14; Danecek *et al*., 2021) with the *view* and the *markdup* program. Finally, the mean depth, percentage of GC and insert size of each individual were checked with the option *stats* of *SAMtools*. The median percentage of reads mapped and properly paired per library was 97.65% for samples from Portugal and 93.95% for outgroups (Figure S2A,B; Table S2). The median read depth was 24.64X for individuals and 52.65X for the pools (Figure S2C; Table S2). We inferred the sex of individuals and sex ratios of pools by comparing the median read depth of the Z chromosome with that of chromosome 1, assuming that males were homogametic ZZ and that females were heterogametic ZW (Figure S3A,B; Table 1, Table S2).

We conducted 4 variant calling using *Freebayes* (v1.3.6; Garrison and Marth, 2012) based on: i) individuals and pools together (ALL VCF), ii) individuals only (IND VCF), iii) pools only (POOL VCF), iv) individuals and pools together for LSP+LWP+FU populations only (DEMO VCF), to be used in different subsequent analyses. For each calling, we used the following parameters, which allow to call variants for either individuals, pools or both (*-C 1, -F 0.01, -G 1, -E -1 --max-complex-gap -1 --haplotype-length -1 -K --limit-coverage 500 -n 3 -m 30 -q 20*). For the specific case of the Z chromosome, the ploidy was set to 1 (*-p* option) for individual females to compute individual genotype likelihoods, while set to 2 for individual males and pools. Remaining multi-nucleotide polymorphisms (MNP) were decomposed using the decompose function of *vt* with the option *-s* (Tan *et al*., 2015). Further filters were applied using *bcftools* (v1.17; Danecek *et al*., 2021), for SNP quality *>*= 30, absence of missing genotypes and missing individuals, reads on both strands and a minimum alternate haplotype count *(INFO/AO >*= 4*, and INFO/AO <*=*(INFO/DP-4))*. Finally, all indels were removed, and only biallelic SNPs were retained. Total site depth filter of *[200-5000]* was applied to ALL VCF, while [200-4000] was applied to IND VCF and POOL VCF, and [100-3000] for the DEMO VCF. This resulted in a total of 8,363,880 SNPs for the ALL VCF, 8,622,533 SNPs for the IND VCF, 9,573,120 SNPs for the POOL VCF, and 7,537,291 SNPs for the DEMO VCF.

Besides, two gVCF files were generated using *bcftools*, with options *-C 50 --max-depth 500 -q 20 -Q 20*: one file (OUT gVCF) including the two highest-depth individuals from each of the LSP, LWP, and FU populations in Portugal, along with all individuals from the outgroup populations, and the second file containing all Portuguese individuals (IND gVCF).

### Characterization of population structure

All analyses were carried out separately on autosomal and Z-linked SNPs. The ALL VCF, IND VCF and POOL VCF datasets were first filtered by discarding low frequency variants (i.e., MAF *<* 5% in at least one population) and pruned based on Linkage Disequilibrium (LD) assessed on all Ind-Seq data using the *--indep* option of *PLINK* (v1.9; Purcell *et al*., 2007) considering a r^2^ threshold of 0.1 and a 20 kb sliding window size (Figure S4). The remaining number of SNPs for autosomes (resp. Z chromosome) was 270,330 (resp. 13,655) for the ALL VCF dataset; 271,712 (resp. 13,705) for the IND VCF dataset; and 268,380 (resp. 13,608) for the POOL VCF dataset. To provide a global view of the structuring of genetic diversity across all the data, we then performed a random allele PCA on the combined Pool-Seq and Ind-Seq data (filtered ALL VCF dataset) using the *randomallele.pca* function implemented in the R package *poolfstats* (v3.0.0; Gautier *et al*., 2022). Unsupervised hierarchical clustering was performed on individual genotypes (from the filtered IND VCF dataset) using *Admixture* v1.3.0 (Alexander *et al*., 2009) considering a number of clusters K ranging from 2 to 10 (the optimal K being assessed using cross-validation run with default settings). Pairwise-population *F*_ST_ were estimated from Pool-Seq data (filtered POOL VCF dataset) using the unbiased estimators (Hivert *et al*., 2018) implemented in the *compute.fstats* function of the R package *poolfstat*. To obtain unbiased estimates of the absolute nucleotide divergence *D*_XY_ for all pairwise population comparisons between LSP, LWP, FU and the outgroups, we relied on unfiltered whole-genome IndSeq data (OUT gVCF) and used *Pixy* v1.2.7, (Korunes and Samuk, 2021). Finally, joint unfolded site frequency spectrum (SFS) graphs were plotted for SP versus WP and for WP versus FU, using allele frequencies among individuals in the POLARIZED VCF.

### Nucleotide diversity, inbreeding and effective population size

Nucleotide diversity (*π*) was estimated for each individual using *Pixy* v1.2.7, (Korunes and Samuk, 2021) on window sizes of 100kb and the IND gVCF. A two-way ANOVA was used to test for *π* differences between the 10 populations and between autosomes and the Z chromosome. Post-hoc tests were conducted using estimated marginal means (i.e., least-squares means or adjusted means) with Bonferroni adjustment.

To characterize individual autozygosity, we relied on the hidden Markov model (HMM) based approach to detect homozygous-by-descent (HBD) segments (Druet and Gautier, 2017) implemented in the R package *RzooRoH* v0.3.2.1 (Bertrand *et al*., 2019). Note that no LD pruning or MAF filtering was carried out (as recommended in e.g., Meyermans *et al*. (2020)) on the analyzed IND VCF dataset. However, SNPs displaying high heterozygosity (Ho *>*= 70%) across all of the PPM individuals, or located within repeated elements identified within the reference genome, were discarded. In addition, since we provided population allele frequencies to *RZooRoH* that were estimated using Pool-Seq data, we applied an additional filter using the *poolfstats* package, requiring a minimum coverage of 10 reads per pool (*min.cov.per.pool = 10*). This resulted in a final dataset of 2,923,200 autosomal SNPs. In the *RZooRoH* models, autozygosity was partitioned into multiple Homozygous By Descent (HBD) classes defined by their rate parameter Rc, which determines their expected length and corresponds to groups of ancestors from different past generations. We here fitted the “layer” model (Druet and Gautier, 2022) specifying 13 HBD classes with rates Rc equal to 2; 4; 8;. . .; 8,192 i.e. corresponding to (modeled) groups of ancestors living 1 to 4096 generations (of recombination) ago. In the absence of dense genetic maps for PPM and to facilitate such timing interpretation, SNP genetic positions were obtained from the physical maps assuming an average genomewide recombination rate of 4 cM per Mb as observed in several other species of *Lepidoptera* (Shipilina *et al*., 2022; i Torres *et al*., 2022; Palahí I Torres *et al*., 2023). The underlying inference model has already been shown to be robust to variable local recombination rate (Druet and Gautier, 2017). As a matter of comparison, we also estimated the fraction of the genome presenting Runs of Homozygosity - RoH (FROH) using *Plink* v1.9 (v1.9; Purcell *et al*., 2007) with the *--homozyg* function, based on the same dataset. Two categories of ROH were tested: ROH *>* 500 kb and ROH *>* 100 kb, to compare ancient and recent inbreeding. We used the following parameters *--homozyg-window-snp 50, --homozyg-snp 50, --homozyg-density 10, --homozyg-gap 100, --homozyg-window-het 2, and --homozyg-window-missing 5*. The individual FROH was calculated by dividing the total length of ROH across all autosomes by the total length of autosomes (579.27 Mb).

Historical effective population sizes (*N_e_*) were inferred with the program *GONe* that implements the approach developed by Santiago *et al*. (2020), assuming a genome-wide average recombination rate of 4 cM/Mb (as justified above). To evaluate the potential impact of recent gene flow on the estimates, we ran analyses with parameter hc (recombination fraction threshold) set to 0.01 or 0.05 (Santiago *et al*., 2020). For each population, we analyzed 100 independent subsets of 50,000 randomly selected SNPs. Near-contemporary *N_e_* at each time point from generation 0 to generation 40 backward in time was then estimated as the harmonic mean of the 100 resulting estimates, following Feng *et al*. (2023). We also estimated the near-contemporary *N_e_* for each population using *currentNe2* (Santiago *et al*., 2025), after removing population-specific monomorphic sites. The analysis was performed with a constant recombination rate of 4 cM/Mb (-r 4) and accounting for population structure (-x).

### Inference of the demographic history among LSP, LWP and FU

We relied on an Approximate Bayesian Computation (ABC) framework to compare different evolutionary scenarios among the sympatric LSP and LWP populations and the more distant winter population FU, and to estimate parameters for the best-supported scenario. Instead of using traditional ABC algorithms, we applied a Random Forest–based ABC approach (ABCRF) (Raynal *et al*., 2019) implemented in the R package *abcrf* (v1.9; Pudlo *et al*., 2016). This method represents a major improvement over classical ABC implementations in several respects. As highlighted by Pudlo et al. (2016), ABCRF (i) often achieves greater discriminative power among competing models, (ii) is more robust to the number and specific choice of summary statistics, (iii) substantially reduces computational cost (by at least an order of magnitude compared to standard rejection or regression-based ABC methods), and (iv) partially resolves the curse of dimensionality by reducing computational complexity in high-dimensional spaces. We compared three scenarios (Figure S5) corresponding to i) a simultaneous divergence of the three populations (concomitant div); ii) a recent divergence of LSP from its sympatric population LWP (recent div); iii) an old divergence of LSP from the two winter populations LWP and FU (old div). Based on results previously obtained by Leblois *et al*. (2018), we introduced the possibility of bottleneck events in every population after the divergence event, and symmetric migration between the different populations up to their common ancestral one.

A total of 158 different summary statistics (see Table S3 for details) were then estimated on the Ind-Seq or the Pool-Seq data (136 and 22 statistics respectively) from the DEMO VCF dataset described above. We then constructed a reference table by computing the same summary statistics on datasets simulated under each of the demographic scenarios using the coalescent simulator implemented in the program *msprime* (v1.2.0; Baumdicker *et al*., 2022). Each simulated genome consisted of 1,000 segments of 10 kb with a fixed per-generation per-base recombination rate *r* = 4.0 *×* 10^−8^ and a fixed mutation rate *µ* = 2.9 *×* 10^−9^, following Leblois *et al*. (2018). For each simulation, model parameters were sampled from the prior distributions detailed in Table S4. To further simulate Pool-Seq datasets, we relied on the *sim.readcounts* function of the R package *poolfstat* (v3.0.0; Gautier *et al*., 2024a) based on the genetic dataset (transformed into sample allele count data) obtained with *msprime* with parameters *lambda.cov* (mean of read coverages assumed to be Poisson distributed) set to 50 for LSP, LWP, and FU pool; *min.rc* (overall minimal read count threshold) set to 2; and *exp.eps* (experimental error rate introducing unequal contribution of individuals to the pool sequences) set to 1%; *seq.eps* (sequencing error rate) of 0.1%; and *genome.size* (to allow simulating spurious SNPs due to sequencing errors that may pass filtering criteria) set to 10 Mb (i.e., 1,000 segments of 10 kb).

To perform model selection, we simulated 50,000 datasets for each of the three demographic scenarios. We then used the *abcrf* function from the *abcrf* R package to build a random forest (RF) of 1,000 trees based on the resulting reference table to ensure stable estimation of the global error rate, as recommended by Collin *et al*. (2021). The best scenario was predicted 10 times using the *predict* function, along with the global and local error rates, across 10 replicate RF analyses based on training sets of 10,000 randomly chosen simulated datasets for each scenario. Finally, the parameter values for the best identified scenario were estimated using regression RF, as implemented in the function *regAbcrf* based on a new reference table constructed with 100,000 simulated datasets.

### SNP annotation and polarization

We annotated and polarized SNPs in order to later inspect populations’ genetic load and search for non-synonymous mutations potentially associated to allochrony.

To predict the potential effect of each SNP, we used *SnpEff* v5.2a (Cingolani *et al*., 2012) and the gff annotation file of the reference genome applied to the complete VCF dataset (ALL VCF). SNPs located within coding regions were categorized as synonymous (SYN”; low predicted impact), missense (MIS”; moderate impact), or loss-of-function (LOF”; high impact) variants. Variants outside coding regions were classified as either intergenic (IG”; located outside exons and splice sites) or “OTHER” (e.g., located within introns or untranslated regions). SnpEff predicted, on the autosomes and the Z respectively, 81,099 and 2,352 SYN, 53,136 and 1,772 MIS, and 764 and 22 LOF.

To determine the ancestral allelic state of each SNP site, we used bam files from a single individual for three outgroups: PPM from Oukaimeden, *T. wilkinsoni*, and *T. bonjeani* and compared their allelic states to those observed in the Portuguese populations under study. For each site, we first assessed whether each outgroup individual was polymorphic, monomorphic, or lacked sufficient coverage. Polymorphic sites were considered monomorphic if the minor allele frequency was within the lowest 10% of the frequency distribution, to avoid misclassifying spurious polymorphisms. Outgroup alleles were only considered informative if coverage *>* 5. Ancestral alleles were defined as the allele shared by at least two outgroups. As a final check, we retained only those sites for which the inferred ancestral allele matched either the reference or alternate allele in our VCF; sites failing this criterion were discarded.

### Estimation of the genetic load in LSP, LWP and FU

To explore the genetic load of each population, we used SNPs called from Ind-Seq data that were successfully polarized and annotated as IG, SYN, MIS, or LOF. Analyses were conducted separately for autosomes and the Z chromosome, with only male individuals included for Z-linked analyses.

For LSP, LWP and FU, considering all the sites that were polymorphic among the Portuguese PPM populations, we estimated expected heterozygosity (He) for the different functional categories of variants (IG, SYN, MIS, LOF). We then calculated, for each individual, the ratio of heterozygosity at non-synonymous sites (including MIS and LOF variants) to heterozygosity at synonymous sites (SYN), expressed as HoN/HoS.

Using the subset of polarized, polymorphic sites shared among LSP, LWP, and FU, we then analyzed the site frequency spectrum (SFS) and the proportions of segregating and fixed ancestral or derived variants across the functional categories. Finally, we used these same data to calculate the *R*_XY_ statistic (Do *et al*., 2015; Grossen *et al*., 2020; Dussex *et al*., 2021) which quantifies the relative excess or deficit of derived alleles in population X compared to population Y. The statistic was computed for the SYN, MIS, and LOF categories, using IG variants as a proxy for neutral expectations. Confidence intervals for *R*_XY_ were obtained via a block jackknife approach following Xue *et al*. (2015), by dividing the genome into 100 blocks with equal numbers of consecutive SNPs. To further investigate the *R*_XY_ patterns, we generated joint SFS plots comparing LSP, LWP, and FU across the different functional categories.

### Search for genomic bases associated with allochrony

To pinpoint genomic regions potentially involved in the phenological shift in LSP, genome scans were performed using the C2 based contrast analysis proposed by Olazcuaga *et al*. (2020) as implemented in the latest version of the *BayPass* (v3.0 Gautier, 2015), which allows joint analyses of Ind-Seq and Pool-Seq data (Camus *et al*., 2024). More precisely, we ran the *BayPass* core model with default parameters on the ALL VCF dataset, with Ind-Seq data provided in the form of genotype likelihoods (GL) and Pool-Seq data in the form of read counts (see the *BayPass* manual for details), analyzing autosomes and the Z chromosome separately. The C2 statistic was defined to contrast samples representative of summer populations (i.e., LSP and LateLSP) and the winter populations (LWP, FU, TA, GR, VA, CA, VI). To identify and delineate significant windows associated with allochrony, i.e. with an excess of high C2 values, we relied on the local-score approach described in Fariello *et al*. (2017) and (re)implemented in the R function *compute.local.scores* available in the *BayPass* (v3.0 Gautier, 2015; Camus *et al*., 2024), using default options but *ξ* = 2.

To go further into the description of the significant windows for association with allochrony identified using *BayPass*, we estimated *F*_ST_ between LSP, LWP, FU, and LateLSP pools over 10 SNPs windows with the *compute.fstats* function from *poolfstat 3.0.0* (using a MAF cutoff of 0.05). We also relied on Ind-Seq data to estimate *D*_XY_ between LSP, LWP, FU, and outgroups, as well as average *π* for LSP and LWP, using 2 Kb window sizes (corresponding on average on the Z chromosome to 10-SNP windows) with *Pixy*, and Tajima’s D with *VCFtools 0.1.16* (Danecek *et al*., 2011). Joint unfolded SFS graphs were plotted for SP versus WP and WP versus FU based on polarized ind-seq data. *LDBlockShow* v1.40 (Dong *et al*., 2021) was used to assess LD patterns. Genotypeplots were used to depict genotypic structure between populations (Whiting, 2022). The read depth of LSP and LWP Ind-Seq libraries were calculated from BAM files using *Mosdepth* (Pedersen and Quinlan, 2018) over 2 kb windows to identify potential candidate CNVs. *Smoove* (https://github.com/brentp/smoove) was used to search for short INDELs and inversions based on Ind-Seq bam files. Lastly, we used *Bedtools* 2.31.1 (Quinlan and Hall, 2010) to list candidate genes located under each window showing high C2 values and we looked for the potential presence of non-synonymous mutations identified using SnpEff.

## Results

### Population structure and differentiation

*F*_ST_, PCA, and Admixture analyses on autosomes revealed strong genetic structure among populations and especially between the summer and winter populations. The pairwise *F*_ST_ values were high between LSP and winter populations, e.g. 0.259 with LWP, 0.197 with FU, and 0.278 with CA (Figure 2A; Table S5), but was only (1.80 *×* 10^−3^) with LateLSP. In contrast, pairwise *F*_ST_ between winter populations was lower, ranging from (3.00 *×* 10^−3^) between FU and VI to 0.216 between LWP and TA. In turn, *D*_XY_ between LSP and LWP was relatively low and comparable to *D*_XY_ between LWP and FU (3.50 *×* 10^−3^ and 3.10 *×* 10^−3^, respectively, Figure 2B; Table S6). *D*_XY_ values between both LSP and LWP versus outgroups were high (1.2 *×* 10^−2^ on average in both cases). The PCA first axis separated the summer populations (LSP and LateLSP), F1 and the other populations (Figure S6). The Admixture analysis yielded a minimal cross-validation error for K = 2, separating summer and winter populations into two distinct genetic clusters (Figure 2C). Increasing the K parameter value further delineated the winter populations (Figure 2D). Finally, while the joint SFS for LSP and LWP showed strong differentiation, with LSP displaying slightly more fixed variants (Figure S7A), the joint SFS for LWP and FU exhibited low differentiation, with LWP showing more fixed variants (Figure S7B).

**Figure 2:**
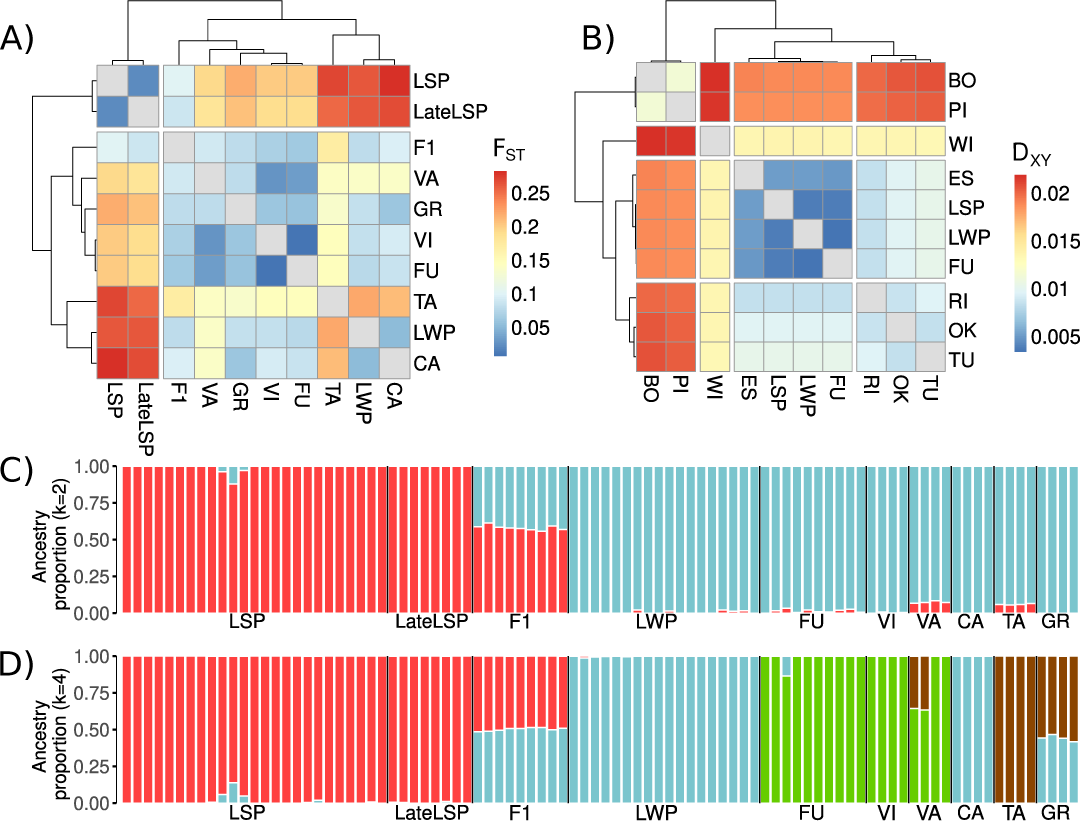
Genetic structure among populations from Portugal. **(A)** Pairwise *F*_ST_ estimates between Portuguese populations. **(B)** pairwise *D*_XY_ estimates between three Portuguese populations (LSP, LWP, and FU) and outgroups. Unsupervised ancestry assignments based on the program Admixture **(C)** at k = 2 and **(D)** at k = 4. All analyses were performed on autosomes. Population codes are defined in Table 1.

### Demographic history inferred using ABCRF

The highest RF classification mean votes (ranging from 391 to 456) and estimated posterior probabilities (ranging from 0.55 to 0.63) provided the strongest support for the recent div compared to the other two scenarios (Table S7, Figure S8). However, although the old div scenario was always the least supported, the concomitant div scenario occasionally received the highest posterior probability in some RF analysis replicates. This is consistent with the relatively recent divergence between LWP and FU, given that the concomitant div scenario represents a limiting case of old div where no differentiation is assumed between the population ancestral to both LWP/LSP and FU.

Figure 3A and Table 2 give the estimated values for the parameters of the recent div scenario (see Figure S9, S10 and S11 and Table S8 for the concomitant div scenario), and revealed a recent divergence between LSP and LWP, occurring approximately a hundred generations ago (74, 90% CI: 13–871). A bottleneck likely occurred around 20 generations ago (CI: 10–129) in all three populations, with estimated effective sizes of few individuals only for LSP (3, CI: 1-174), 13 individuals in LWP (CI: 1-692) and 91 in FU (CI: 2-13,490; Figure 3A; Table 2). Contemporary gene flow among the three populations was significantly reduced following this bottleneck, with the most pronounced reduction observed between LSP and LWP (6.17 *×* 10^−4^; 90% CI: 7.94 *×* 10^−5^ – 3.98 *×* 10^−3^). Finally, LSP showed the lowest current *N_e_* (645 individuals; 90% CI: 115–51,286), while FU showed the highest current *N_e_*(13,182 individuals; 90% CI: 575–114,815). Parameters specifying past events, such as ancient migration or ancient *N_e_*, and founder events were not accurately estimated with large Normalized Mean Absolute Error and flat posterior distribution (Figure S12 and S13; Table 2).

**Figure 3:**
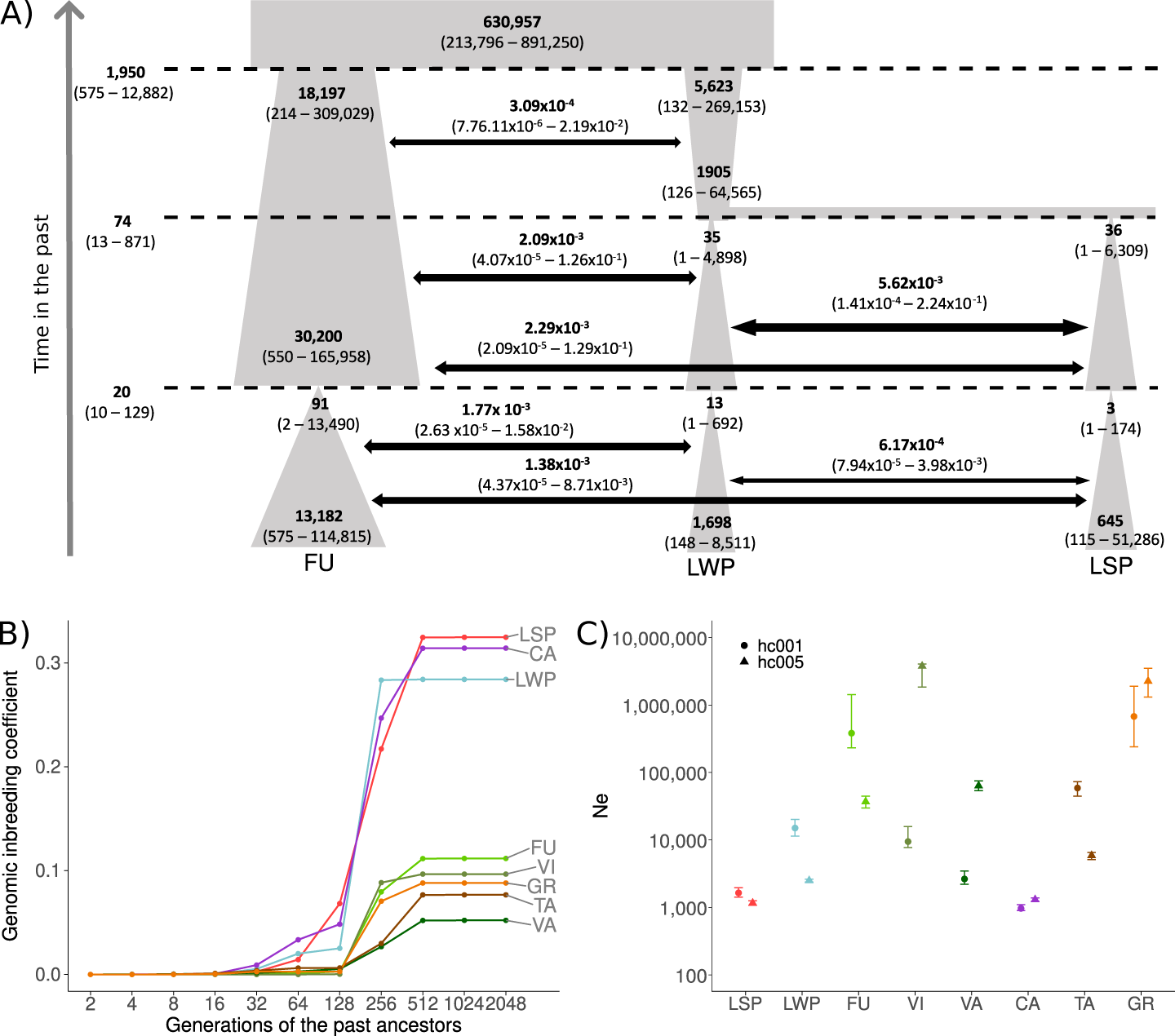
Inference of demographic history, inbreeding and effective population sizes. **(A)** Graphical representation of the best demographic scenario, recent div, of divergence between LSP, LWP and FU populations. Split time and bottleneck time (horizontal dashed lines), populations size and migration rates (horizontal arrows) were estimated from 100,000 simulations. Bold values correspond to the median estimates of the parameters, and numbers in parentheses indicate 95% confidence intervals. **(B)** Genomic inbreeding inferred by RZooRoH on autosomes. Each line represents the mean value of the individual genomic inbreeding coefficient for a population, cumulated through time. The X-axis represents the generations of ancestors contributing to each HBD class. Only HBD classes ranging from 2 to 2048 generations are shown. **(C)** Near-contemporary effective population size (0 to 50 generations) estimated with GONE on autosomes. For each population, the mean and range of *N_e_* estimates obtained using a recombination fraction (hc) of 0.01 and 0.05 are shown.

**Table 2:** Results for estimation of parameters of interest for the scenario recent div. Mean and median estimates for each parameter were calculated using an ABCRF run conducted with a reference table containing 158 summary statistics from 100,000 simulations. Quantiles at 5% (lower bound) and 95% (upper bound) were used to obtain the 90% confidence interval (CI). Mean and median global and local NMAE (normalized mean absolute error, which is the average absolute difference between the point estimate and the true simulated value divided by the true simulated value) were also estimated.

| Parameter | Posterior point estimates of |  | 90% CI |  | Global (prior) NMAE computed from |  | Local (posterior) NMAE computed |  |
| --- | --- | --- | --- | --- | --- | --- | --- | --- |
| Parameter | Mean | Median | Lower bound | Upper bound | Mean | Median | Mean | Median |
| $m_{ancPP}$ | $3.24 \times 10^{-4}$ | $3.09 \times 10^{-4}$ | $7.76 \times 10^{-6}$ | $2.19 \times 10^{-2}$ | 2.12 | 2.07 | 0.45 | 0.45 |
| $m_{ancLSPFU}$ | $1.74 \times 10^{-3}$ | $2.29 \times 10^{-3}$ | $2.09 \times 10^{-5}$ | $1.29 \times 10^{-1}$ | 2.34 | 2.29 | 0.68 | 0.67 |
| $m_{recLSPFU}$ | $9.55 \times 10^{-4}$ | $1.38 \times 10^{-3}$ | $4.37 \times 10^{-5}$ | $8.71 \times 10^{-3}$ | 0.85 | 0.76 | 0.17 | 0.16 |
| $m_{ancLSP LWP}$ | $5.75 \times 10^{-3}$ | $5.62 \times 10^{-3}$ | $1.41 \times 10^{-4}$ | $2.24 \times 10^{-1}$ | 2.49 | 2.46 | 1.10 | 1.11 |
| $m_{recLSP LWP}$ | $6.03 \times 10^{-4}$ | $6.17 \times 10^{-4}$ | $7.94 \times 10^{-5}$ | $4.07 \times 10^{-3}$ | 0.99 | 0.91 | 0.13 | 0.13 |
| $m_{ancLWP FU}$ | $2.19 \times 10^{-3}$ | $2.09 \times 10^{-3}$ | $4.07 \times 10^{-5}$ | $1.26 \times 10^{-1}$ | 1.57 | 1.53 | 0.72 | 0.71 |
| $m_{recLWP FU}$ | $1.15 \times 10^{-3}$ | $1.78 \times 10^{-3}$ | $2.63 \times 10^{-5}$ | $1.58 \times 10^{-2}$ | 0.84 | 0.76 | 0.20 | 0.18 |
| $N_e_{anc.PP}$ | $5.13 \times 10^5$ | $6.31 \times 10^5$ | $2.14 \times 10^5$ | $8.91 \times 10^5$ | 0.01 | 0.01 | 0.00 | 0.00 |
| $N_e_{bot.LSP}$ | $5.49 \times 10^0$ | $3.24 \times 10^0$ | $1.15 \times 10^0$ | $1.74 \times 10^2$ | 4.56 | 3.90 | 4.64 | 3.10 |
| $N_e_{bot.FU}$ | $1.10 \times 10^2$ | $9.12 \times 10^1$ | $2.14 \times 10^0$ | $1.38 \times 10^4$ | 2.79 | 2.34 | 3.25 | 3.04 |
| $N_e_{bot.LWP}$ | $1.78 \times 10^1$ | $1.26 \times 10^1$ | $1.51 \times 10^0$ | $6.92 \times 10^2$ | 4.17 | 3.58 | 1.73 | 1.47 |
| $N_e_{LSP}$ | $1.07 \times 10^3$ | $6.46 \times 10^2$ | $1.15 \times 10^2$ | $5.13 \times 10^4$ | 0.19 | 0.18 | 0.23 | 0.21 |
| $N_e_{FU}$ | $1.12 \times 10^4$ | $1.32 \times 10^4$ | $5.75 \times 10^2$ | $1.15 \times 10^5$ | 0.14 | 0.14 | 0.14 | 0.14 |
| $N_e_{LWP}$ | $1.38 \times 10^3$ | $1.70 \times 10^3$ | $1.51 \times 10^2$ | $8.51 \times 10^3$ | 0.14 | 0.14 | 0.16 | 0.16 |
| $N_e_{Old.FU}$ | $1.95 \times 10^4$ | $3.02 \times 10^4$ | $5.50 \times 10^2$ | $1.66 \times 10^5$ | 0.19 | 0.18 | 0.16 | 0.16 |
| $N_e_{Old.LWP}$ | $2.34 \times 10^3$ | $1.86 \times 10^3$ | $1.26 \times 10^2$ | $6.46 \times 10^4$ | 0.21 | 0.20 | 0.23 | 0.22 |
| $N_e_{ancfound.FU}$ | $1.23 \times 10^4$ | $1.82 \times 10^4$ | $2.14 \times 10^2$ | $3.09 \times 10^5$ | 0.24 | 0.24 | 0.23 | 0.23 |
| $N_e_{ancfound.LWP}$ | $6.03 \times 10^3$ | $5.62 \times 10^3$ | $1.32 \times 10^2$ | $2.69 \times 10^5$ | 0.25 | 0.25 | 0.28 | 0.28 |
| $N_e_{found.LWP}$ | $4.79 \times 10^1$ | $3.55 \times 10^1$ | $1.55 \times 10^0$ | $4.90 \times 10^3$ | 48.49 | 46.37 | 4.29 | 3.91 |
| $N_e_{found.LSP}$ | $4.90 \times 10^1$ | $3.63 \times 10^1$ | $1.35 \times 10^0$ | $6.46 \times 10^3$ | 5.46 | 4.92 | 7.88 | 7.50 |
| $T_{split.PP}$ | $2.24 \times 10^3$ | $1.95 \times 10^3$ | $5.75 \times 10^2$ | $1.29 \times 10^4$ | 0.08 | 0.07 | 0.09 | 0.09 |
| $T_{split.LWP}$ | $8.71 \times 10^1$ | $7.41 \times 10^1$ | $1.29 \times 10^1$ | $8.71 \times 10^2$ | 0.24 | 0.23 | 0.25 | 0.25 |
| $T_{bot}$ | $2.51 \times 10^1$ | $2.04 \times 10^1$ | $1.05 \times 10^1$ | $1.29 \times 10^2$ | 0.22 | 0.20 | 0.19 | 0.17 |

### Population diversity, inbreeding, and effective population size

Nucleotide diversity (*π*) estimated over all autosomes varied greatly among populations (2-ways ANOVA, p-value *<* 0.001), in line with very different demographic histories. The genetic diversity was the lowest in the LSP and LateLSP populations (*π* = 2.1 *×* 10^−3^); intermediate in the LWP and CA populations (*π* = 2.5 *×* 10^−3^); and ranged from *π* = 3.0 *×* 10^−3^ (for TA) to *π* = 3.4 *×* 10^−3^ (for VA) in the other Portuguese populations (Figure S14).

*RZooRoH* analyses showed much higher levels of genomic inbreeding coefficients in LSP, LWP and CA (from 0.284 for LWP to 0.325 for LSP) compared to other populations (from 0.052 for VA to 0.112 for FU). Importantly, *RZooRoH* inferred that most autozygosity concentrated in HBD segments was associated with ancestors living 64 to 256 generations ago (corresponding to classes 128 to 512; Figure 3B), while little genomic inbreeding originated from more recent (*<* 64 generations) or more ancient (*>* 512 generations) ancestors. Moreover, there was little inter-individual variance in the level of inbreeding and the age of large HBD segments within each population (Figure S15). In line with these analyses, LSP, LWP and CA exhibited high FROH values ranging from 0.280 to 0.336, approximately 3 times higher than those of other Portuguese populations, which ranged from 0.059 (for VA) to 0.106 (for FU, Figure S16). The same pattern was observed for the fraction of the genome covered by ROH *>* 500 Kb (Figure S17A). Similarly, the number of both short and long ROH displayed comparable patterns (Figure S17B,C,D).

Finally, the near contemporary *N_e_* estimated with *GONe* indicated a relatively small recent *N_e_* (*<* 1000) for the LSP summer population and its two LWP and CA neighbors *N_e_ <* 5000 and *N_e_ <* 1000, respectively, compared to more distant populations (Figure 3C, Figure S18A,B). Similar results were found using *currentNe2*, with *N_e_* of 999 in LSP (90% CI:540-1849), 652 in LWP (90% CI=330-1289), and 113 in CA (90% CI:33-391). However, *currentNe2* was unable to converge for FU, VI, VA, TA, and GR, which was likely due to the large *N_e_* of these populations.

### Genetic load

Expected heterozygosity was lower in LSP compared to LWP and in LWP compared to FU, for each category of sites (Figure 4A). For each population, expected heterozygosity was lower at LOF sites compared to MIS sites, and at MIS sites compared to SYN and IG sites (Figure 4A). HoN/HoS ratio was higher in LSP and LWP compared to FU (Post-hoc Dunn’s tests with Bonferroni correction; p-values = 1.7 *×* 10^−3^ and 2.0 *×* 10^−5^, respectively) and was not different between LSP and LWP (p-value = 3.5 *×* 10^−1^; Figure 4B). There were lower proportions of homozygous derived genotypes at LOF sites, compared to MIS sites, and at MIS sites compared to SYN and IG sites, in each population (Figure 4C), suggesting an effect of purifying selection at MIS sites and more strikingly at LOF sites. However, proportions of homozygous derived genotypes were higher in LSP than in LWP and in LWP than in FU, for every category of variant (Figure 4C), suggesting accumulation of genetic load in LSP compared to LWP and in LWP compared to FU, likely resulting from reduced purifying selection and increased drift. The SFS (Figure S19) indicated similar trends. *R*_XY_ values *>* 1 were observed when comparing LSP to LWP, LSP to FU, and LWP to FU, for the LOF sites (Figure 4D), indicating a relative accumulation of genetic load at LOF sites in LSP compared to LWP and FU, and in LWP compared to FU. *R*_XY_ values for the SYN and MIS classes were close to 1, suggesting little differences in purge-accumulation of genetic load signals at these sites between populations. Yet, *R*_XY_ values were not significantly different to 1 when using z-score two-tailed test. Joint SFS between populations for each category of sites illustrated the accumulation of derived alleles at LOF variants in LWP and more importantly in LSP, compared to FU (Figure S20).

**Figure 4:**
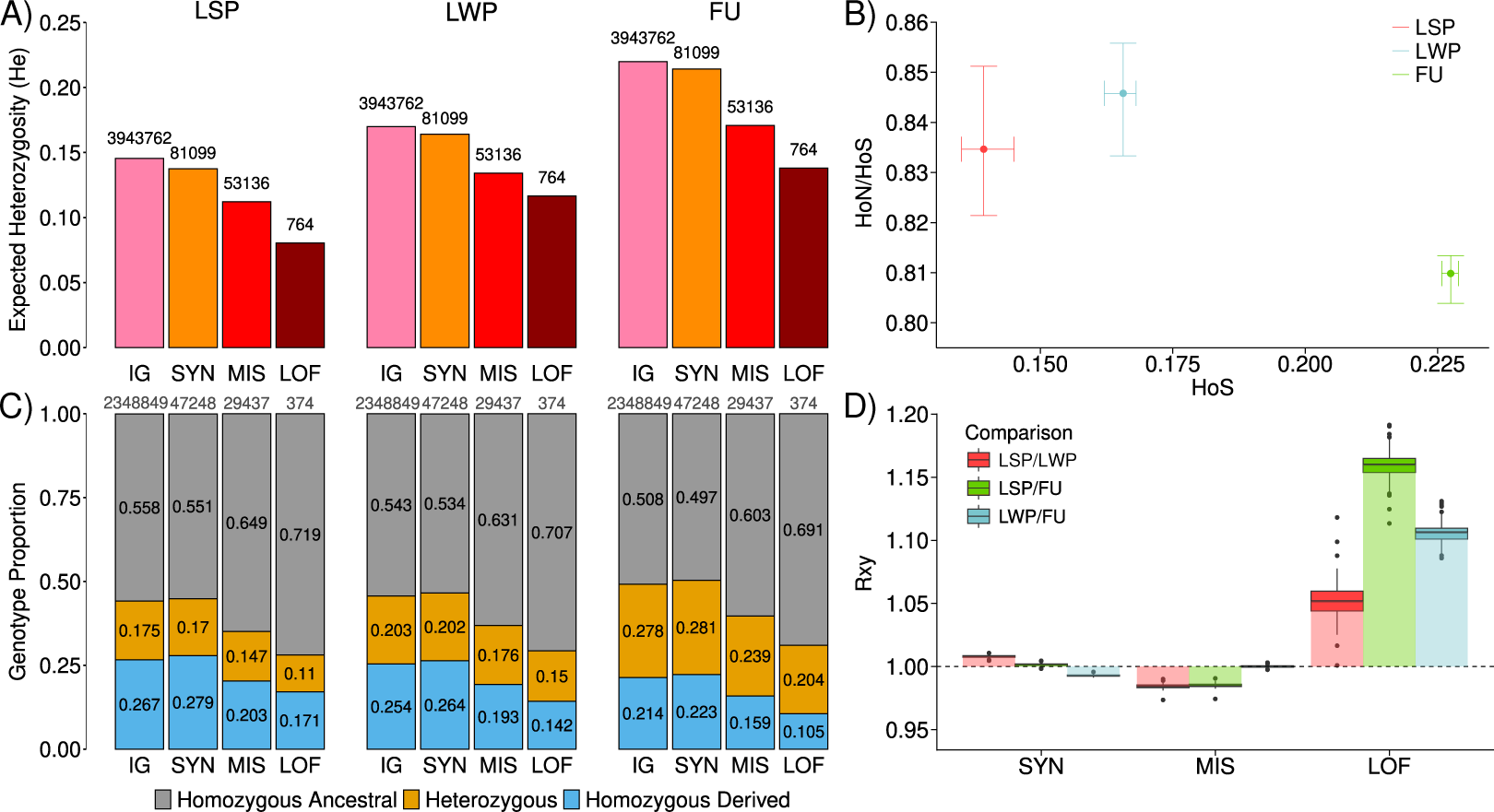
Estimates of genetic load in LSP, LWP and FU on autosomes. **(A)** Expected heterozygosities at IG, SYN, MIS, and LOF sites, with the count of variants indicated above each category. **(B)** Relationship between the ratio of individual heterozygosity at MIS (HoN) to SYN (HoS) sites and individual heterozygosity at SYN (HoS) sites. Each point represents the population median, with confidence intervals based on individual-level variation. **(C)** Proportions of genotypes that were either homozygous ancestral, homozygous derived or heterozygous, at IG, SYN, MIS, and LOF sites, with the count of variants indicated above each category. **(D)** *R*_XY_ statistics at SYN, MIS, and LOF sites.

### Genome scans between summer and winter populations

The C2 contrast analyses comparing summer and winter population samples allowed identifying 135 significant windows (102 genes) on autosomes and 20 windows (25 genes) on the Z chromosome (Figure 5A, 5B; Table S9). Among these significant windows, some —including those showing the highest C2 values and their surrounding regions (+/- 10 kb)— contain several genes known to influence biological rhythms: *NPAS2* (neuronal PAS domain-containing protein 2-like), *GABA_B_* (gamma-aminobutyric acid receptor subunit beta-like), *Coq8* (atypical kinase COQ8B, mitochondiral), *NEURAL4 / NHR* (neuralized-like protein 4), *NEP-2* (neprilysin-2) and *SP2353* (basement membrane-specific heparan sulfate proteoglycan core protein-like). Although the *period* gene was not in a C2 local score window, it lied - 100 kb upstream of the second highest C2 local score window. *F*_ST_ between either LSP and LWP and between LateLSP and LWP were on average high across the entire Z chromosome, while *F*_ST_ between LSP and LateLSP were consistently close to zero (Figure 5C). The region around *NEURAL4 / NHR* and *NEP-2* showed particularly high *F*_ST_ values between either LSP or LateLSP and LWP (Figure 5D). This genetic differentiation pattern was similar across the several comparisons between LSP and winter populations (Figure S21).

**Figure 5:**
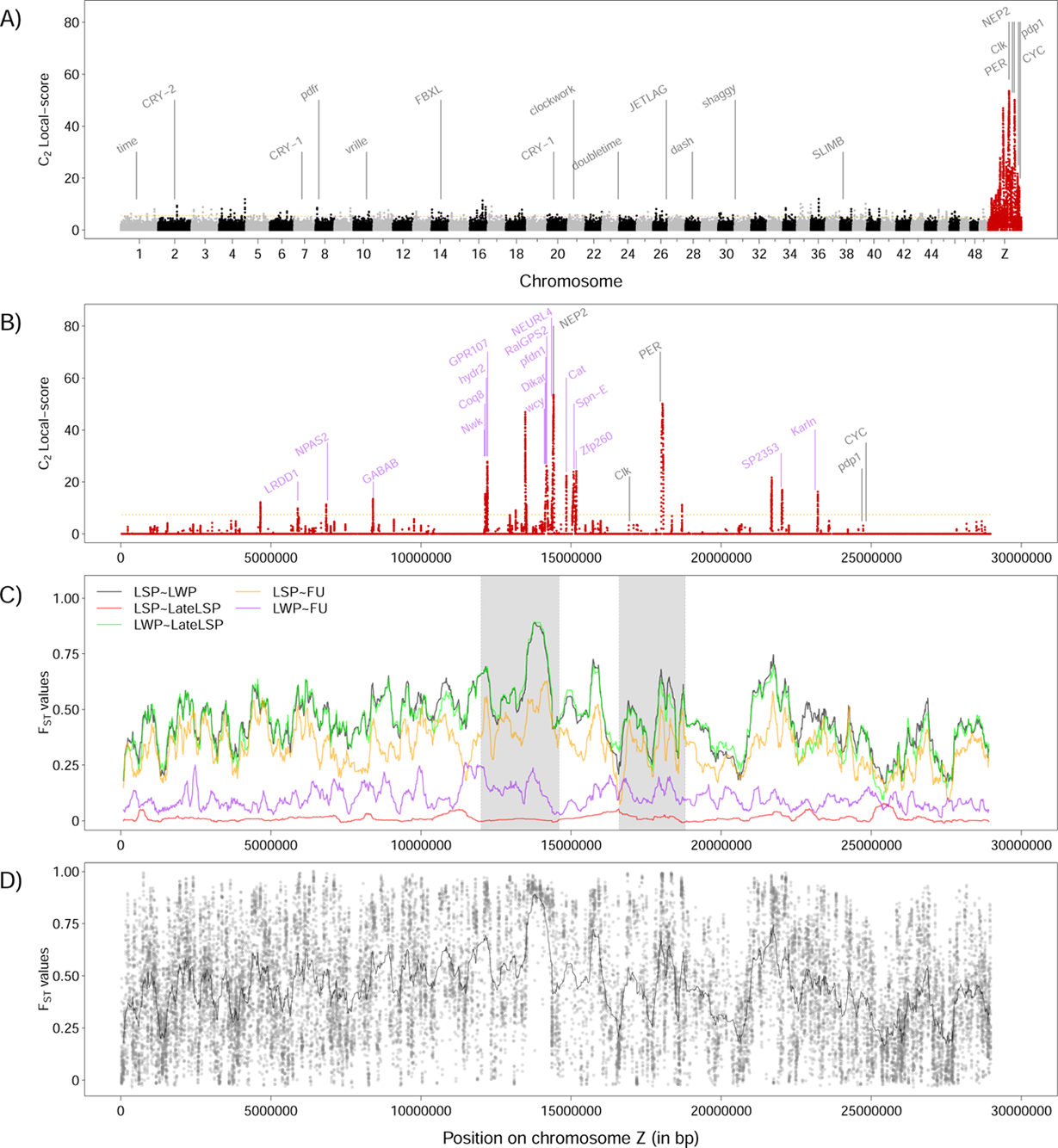
Genome scan looking for loci underlying phenological variations. Manhattan plot of the Local-score based on C2 p-values for the entire genome **(A)** and for the Z chromosome only **(B)**. The horizontal dashed lines indicate the chromosome-specific 1% significance threshold for the local score. Genes potentially linked to circadian rhythm and diapause are written in grey. Z-linked genes related to other functions and identified in high C2 local scores are written in purple. **(C)** *F*_ST_ values along the Z chromosome for five population pairs. The light grey rectangles correspond to the regions zoomed in on Figure 6. **(D)** *F*_ST_ values along the Z chromosome between LSP and LWP. Each point represents the estimated *F*_ST_ value between LSP and LWP pools for 10-SNPs windows.

Among the Z chromosome candidate windows, we focused on two regions of interest, showing high C2 and *F*_ST_ and including *Coq8*, *NEURL4*, *NEP-2*, *Clk* (circadian locomoter output cycles protein kaput) and *period* genes.

First, in the surroundings of the *Coq8* gene, from 12.10 to 12.25 Mb (Figure 6A; Table S10), we observed high C2 local score windows and *F*_ST_ value between LSP and LWP pools (Figure 6C, 6E). *π* in this region was low in both LSP and LWP (Figure 6G). Tajima’s D was negative in LSP while it was close to 0 for LWP (Figure 6I). One potentially deleterious amino acid change from Val to Ala, differentially fixed between LWP and LSP, was found in the *Coq8* gene (Table S11).

**Figure 6:**
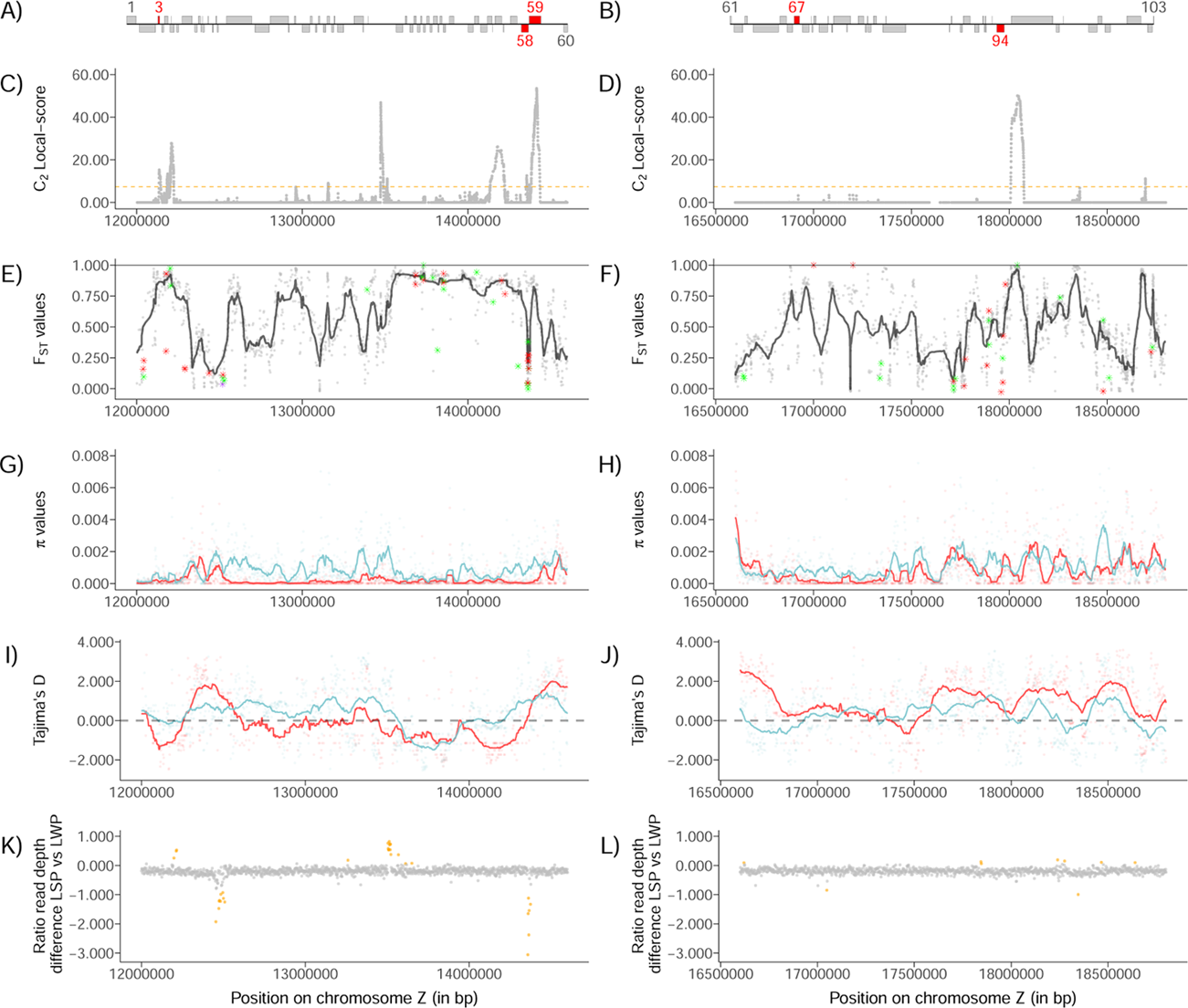
Genetic characteristics of two genomic regions, from 12.00 Mb to 14.60 Mb and from 16.60 Mb to 18.80 Mb, showing high genetic contrasts between LSP and LWP. **(A & B)**: Annotated genes (see also Table S10) present within both genomic regions. Red rectangles indicate genes associated with functions related to circadian rhythm and diapause in insects (3: *Coq8*; 58: *NEURL4*; 59: *NEP-2*; 67: *Clk*; 94: *period*) and grey rectangles represent other genes. **C & D**: Manhattan plots of the Local-score based on C2 p-values, zoomed in on the Z chromosome. The black dashed line represents the chromosome-specific 1% significance threshold for the local score. **E & F**: Pairwise *F*_ST_ between LSP and LWP. Each point represents an estimated value calculated over a 10-SNP sliding window. Green, red and purple stars respectively indicate tolerant, deleterious and loss of function mutations, with their SNP *F*_ST_ values. **G & H**: Pi value of LSP (in red) and LWP (in blue), with each point representing the average Pi over a 2,000 bp window. **I & J**: Tajima’s D in LSP (in red) and LWP (in blue), where each point represents the average Tajima’s D over a 2,000 bp window. **K & L**: Read depth difference between LSP and LWP (see methods), with each point corresponding to average depth over a 2,000 bp window. Negative values indicate an excess of read depth in LWP relative to LSP, and positive values indicate the opposite. Orange dots indicate SNPs exceeding the 99th percentile threshold of the distribution.

Then, the region between 13.60 and 14.30 Mb, upstream of the *NEURL4* and *NEP-2* genes (figure 6A; Table S10), showed several windows with a high C2 local score (Figure 6C) and an almost constantly high *F*_ST_ value between LSP and LWP pools (Figure 6E). The joint SFS between LSP and LWP individuals in this region showed a large proportion of SNPs with opposite allele frequencies (Figure S7C), while LWP and FU exhibited more similar allele frequencies (Figure S7D). This was also clearly depicted by genotype plots (Figure S22, S23). LD was particularly high in this highly differentiated region (Figure S24). *π* in this region was lower in both LSP (mean: *π* = 2.0*×*10^−4^) and LWP (mean: *π* = 8.0 *×* 10^−4^) relative to the Z chromosome’s *π* average (mean LSP: *π* = 7.0 *×* 10^−3^; mean LWP: *π* = 1.3 *×* 10^−3^), and *π* was here lower in LSP than in LWP (Figure 6G). In addition, Tajima’s D was negative over large regions both in LSP and in LWP, from 13.33 Mb to 14.33 Mb and 13.57 Mb to 14.10 Mb, and more negative in LSP than in LWP from 13.90 Mb to 14.33 bp (Figure 6I). The *D*_XY_ scan between LSP and LWP or LSP and FU in this region revealed no significantly high values (Figure S25) and the *D*_XY_ tree’s topologies remained consistent between the entire Z chromosome and this highly differentiated region (Figure S26). Although a few small structural variants were detected by the *Smoove* analysis, we did not identify large INDELs or inversions in this region (not shown). *Mosdepth* nevertheless identified from 14,358,000 to 14,376,000 bp, within *NEURL4* and upstream *NEP-2* (Figure 6K), a lower read depth in LSP (18.13 X on average) than in LWP (42.70 X on average), which may reflect a short duplication or CNV in LWP and/or deletion in LSP. In this specific region, we found an excess of heterozygous individuals in LWP (Ho = 0.83; 140 SNPs with Ho *>* 0.5), but not in LSP (Ho = 0.25; 0 SNP with Ho *>* 0.5), which may reflect the presence of a polymorphic duplication in LWP which was absent (i.e. deleted) in LSP (Figure S27). Evidence of this putative duplication was also found in FU and other winter populations, but not in LateLSP. The excess of heterozygosity in LWP in this region between 14,358,000 to 14,376,000 bp was responsible for the observed narrow drop in *F*_ST_ values at this region (Figure S27). No deleterious mutation with a strong differentiation between LSP and LWP was found in *NEP-2* or *NEURL4*. However, there were several potentially deleterious amino acid changes in the *NEURL4* region showing an excess of read depth and of heterozygosity in LWP, that were at intermediate frequencies in LWP (Table S11) and fixed or almost fixed in LSP and that should hence be considered carefully.

In the surroundings of the *Clk* gene, from 16.93 to 16.96 Mb (Figure 6B; Table S10), we observed high *F*_ST_ windows (Figure 6D, 6F). *π* in this region was low in both LSP and LWP (Figure 6H). Tajima’s D was slightly negative in LWP and positive or close to 0 in LSP, from 16.50 Mb to 17.40 Mb (Figure 6J). No deleterious mutation with strong differentiation between LSP and LWP was found in *Clk*.

Immediately downstream of the *period* gene (Figure 6B) from 18.00 Mb to 18.05 Mb, we observed a window with a high C2 local score and a cluster of windows with high *F*_ST_ values (Figure 6D,F; Table S10). There was no strong decrease in *π* (Figure 6H) nor of Tajima’s D (Figure 6J). One potentially deleterious amino acid change from Val to Ala showing a frequency of 0 in LWP and 0.8 in LSP was found in the *period* gene (Table S11).

### Genetic differentiation, diversity and genetic load on the Z chromosome

Pairwise *F*_ST_ values were higher for the Z chromosome than for autosomes for most of the population pairs (Figure 7A, Figure S28; Table S12). However, this higher differentiation at the Z chromosome was much more pronounced for population pairs including either LSP or LateLSP. In turn, the *D*_XY_ values for the Z chromosome between LSP or LateLSP were comparable to those of the winter Portuguese populations, although *D*_XY_ on autosomes was globally slightly higher than on the Z chromosome (Figure S29,S30; Table S13). *π* values varied significantly between autosomes and the Z chromosome across populations (ANOVA, p-value ¡ 0.001, Figure S31), with a particularly reduced *π* on the Z chromosome in LSP and LateLSP (median values of 6.0*×*10^−4^), compared to other winter populations (median values ranging from *π* = 1.3 *×* 10^−3^ to *π* = 1.9 *×* 10^−3^). Interestingly, while LSP and LWP showed relatively low autosomal *π* compared to other populations, the average ratio of *π* on the Z chromosome over *π* on autosomes was significantly lower for LSP (0.244; Figure 7B) than for LWP (0.523) or the other Portuguese populations (ranging from 0.507 to 0.590), suggesting a striking reduction of genetic diversity on the Z chromosome in LSP.

**Figure 7:**
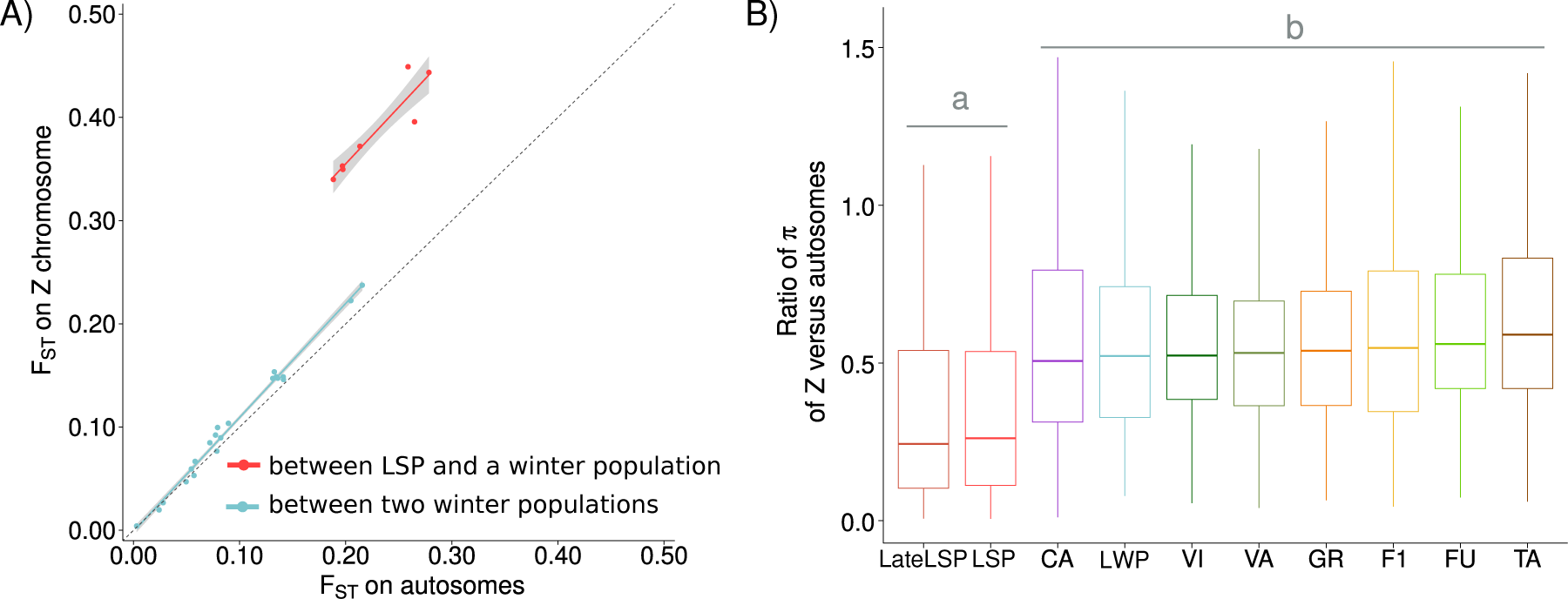
Over-differentiation and reduced nucleotide diversity on the Z chromosome in LSP. **(A)** Pairwise *F*_ST_ values on the Z chromosome compared to pairwise *F*_ST_ values on autosomes. The red dots represent *F*_ST_ values involving comparisons between LSP and winter populations, and the blue dots indicate *F*_ST_ values for comparisons between winter populations. **(B)** Ratio of the median nucleotide diversity (*π*) calculated for the Z chromosome over the median *π* calculated for autosomes, for male individuals. Statistically significant differences were indicated letter *a* and *b*.

The PCA on the Z chromosome showed a clear population genetic structure (Figure S32), with the F1 females grouping either with LSP or LWP individuals (as expected given that they inherited only one Z copy from either a LSP father or a LWP father). Accordingly, the admixture analysis on the Z chromosome revealed the two main distinct genetic groups corresponding to the summer and winter populations (Figure S33A,B), with F1 males mixed between both groups and females grouping with either LSP or LWP individuals. The minimum cross-validation error was observed at K = 2, delineating summer and winter populations. Finally, the joint SFS for LSP and LWP showed strong differentiation on the Z chromosome, with many differentially fixed variants, especially between LSP and LWP (Figure S7E). The joint SFS for LWP and FU on the Z chromosome exhibited less differentiation (Figure S7F).

Similarly as for autosomes, He at the Z chromosome was lower at LOF sites compared to MIS sites, and at MIS sites compared to SYN and IG sites, across all three populations (Figure S34A). Additionally, expected heterozygosity was consistently lower in LSP compared to LWP, and in LWP compared to FU, across all categories of variants. HoN/HoS ratio was higher in LSP and LWP compared to FU (p-values = 2.2 *×* 10^−2^ and 3.5 *×* 10^−2^, respectively) and was not different between LSP and LWP (p-value = 1; Figure S34B). In contrast to autosomes, the proportion of homozygous derived genotypes at polarized LOF sites was lower in LSP compared to LWP and FU (Figure S34C), although there were few LOF sites on the Z (n = 12). Very low *R*_XY_ values were observed when comparing LSP to LWP and LSP to FU at LOF sites (Figure S34D), suggesting stronger purging of LOF derived alleles in LSP compared to LWP and FU. The site frequency spectrum (SFS, Figure S35) supported these trends. Joint SFS analyses further illustrated an excess of derived alleles at SYN and MIS sites in LSP compared to LWP and LSP, but the opposite patterns for the few LOF sites (Figure S36).

## Discussion

In this study, we investigated the genetic causes and consequences of the allochronic divergence between pine processionary moth populations in Portugal. We empirically explored theoretical predictions (Devaux and Lande, 2008; Weis *et al*., 2014) proposing that small *N_e_*, non-selective assortative mating based on the reproduction date, and the existence of polymorphisms at genes controlling for the phenology of reproduction would promote the onset of allochronic divergence. We found a strong differentiation between LSP and LWP sympatric allochronic populations, together with evidence for recent gene flow reduction (but not to the point of total isolation), as previously shown for this system (Santos *et al*., 2011; Burban *et al*., 2016, 2020; Leblois *et al*., 2018). In addition, we found statistical support for recent strong inbreeding and reduction of *N_e_* not only in LSP but also in LWP, suggesting that a common environmental force might have driven a reduction in size of both populations. Rather than being purged, the genetic load in LSP appears to have increased, likely as a consequence of strong drift and reduced efficacy of selection. Our most salient result is the identification of a marked reduction in genetic diversity and enhanced differentiation in several Z-linked regions associated with allochrony, including genes known to be implicated in the regulation of biological rhythms in insects. Moreover, the entire Z chromosome in LSP showed an over-differentiation and a strong reduction in nucleotide diversity, compared to other populations, likely indicating both a strong bottleneck and the onset of a barrier to gene flow on the LSP Z chromosome. We discuss how these empirical genomic findings support theoretical expectations that assortative mating driven by reproductive timing differences underpinned by polymorphisms in circadian genes, along with enhanced genetic drift and purge of genetic load, can promote the onset and persistence of allochronic divergence.

### Demographic history of the allochronic summer population

The most likely scenario of divergence of LSP inferred by ABCRF suggested a split between LSP and LWP approximately 74 generations before sampling, concomitant with strong bottlenecks in both populations. In line with this result, RzooRoh analyzes detected inbreeding generated 64-256 generations ago in LSP, LWP and CA. Similar patterns were reported, for example, by Gautier *et al*. (2024b), who observed a high proportion of short HBD segments in a cattle population that experienced a strong recent founder effect followed by rapid expansion. Importantly, these results of a recent split between LSP and LWP and recent inbreeding in both populations support a scenario of recent sympatric allochronic divergence caused by increased genetic drift, in line with theoretical expectations from Devaux and Lande (2008). Conversely, the results do not support alternative divergence scenarios, such as an ancestral divergence of LSP (potentially in parapatry) and/or a major role of spatially varying selection acting on phenology.

Among the potential environmental drivers of such recent bottlenecks and inbreeding, the management of the pine forest in Portugal may have played an important role. Indeed, large-scale intensive afforestation in Mata National de Leiria started in 1892 and continued throughout the 20th century (Pinho, 2012) with a particular intensification in the 1930s, primarily using maritime pine *Pinus pinaster*. These efforts expanded further in the 1960s, incorporating clear-cut practices and the development of 120 km of forest roads. This widespread plantation of maritime pine may have facilitated the establishment of new populations in which bottlenecks may have occurred via founder effects. In addition, important forest fires have been documented in Portugal, especially in pine tree forests (Mendes, 2007; Aguiar *et al*., 2021), which potentially triggered bottlenecks in pine processionary moth populations.

The fact that our estimated split time between LSP and LWP was slightly more recent than the one inferred by Leblois *et al*. (2018) (560 generations) could be due to the fact that we included as a third population in our ABCRF analysis a more distant and more diverse winter population (FU) compared to Leblois *et al*. (2018) (CA). Besides, the accuracy of demographic parameter estimation may have been improved by the use of a much larger SNP dataset and a more precise SNP calling thanks to the use of a new chromosome-level reference genome (Fuentes-Pardo and Ruzzante, 2017; Martchenko and Shafer, 2023). In addition, using *msprime* enabled the simulation of numerous coalescent scenarios (Kelleher *et al*., 2019; Liu *et al*., 2020) and using ABCRF allowed further improvement of the accuracy of parameter inference by incorporating a diverse set of numerous nonindependent summary statistics from PoolSeq and IndSeq for parameter estimation (Pudlo *et al*., 2016; Raynal *et al*., 2019).

Despite evidence of recent strong bottlenecks in LSP and LWP, intermediate (i.e. relatively high) values of contemporary *N_e_* were estimated (645 using ABCRF and 1,000 using *GONe*). Such *N_e_* in LSP is much lower than *N_e_* estimates sometimes exceeding 10,000 in several other Lepidoptera species (Bortoluzzi *et al*., 2023; García-Berro *et al*., 2023), but is also considerably higher than in endangered or compromised populations of other Lepidoptera species where *N_e_* often falls below 100 or even 50 individuals (Andersen *et al*., 2014; Marí-Mena *et al*., 2019). Hence, the contemporary *N_e_* values estimated for LSP appear high relative to the fact that this population experienced a relatively recent severe bottleneck and is reproductively isolated due to its allochrony. Such relatively high contemporary *N_e_* in LSP may be first explained by the persistence of a certain amount of gene flow from winter populations into LSP, at least at autosomal regions of the genome. In addition, such a large *N_e_* could have been caused by a rapid growth of the LSP population, which is in line with the observed geographical expansion of LSP along the coast of Portugal (Godefroid *et al*., 2016). Such growth could have been facilitated by a relaxation of both intraspecific competition and predation or parasitism by natural enemies (Santos *et al*., 2013). LSP population growth could have also been favored by local climatic conditions compatible (although not optimal) with a summer life cycle (Santos *et al*., 2011; Godefroid *et al*., 2016).

Our three-time lower estimation of *m* between LSP and LWP than between LWP and FU is consistent with the results from Leblois *et al*. (2018) who also reported reduced gene flow between LSP and winter populations. Although the restricted gene flow between LSP and LWP is consistent with the LSP population allochronic life cycle leading to reproductive isolation, it is higher than expected under a strict allochrony. This may notably be explained by phenological plasticity in the summer population. Some individuals - referred to as LateLSP - notably emerge with a winter phenology while they have a summer (LSP) genetic background. Moreover, hybrids between LSP and LWP genetic backgrounds have already been detected in nature (Burban *et al*., 2016) and may then reproduce with either LSP or LWP individuals (see next discussion section). However, such gene flow between LSP and LWP challenges the possibility of maintaining these two genetically differentiated allochronic populations. One explanation for the maintenance of LSP despite gene flow may be the existence of a heterogeneous gene flow along the genome, with reduced gene flow at regions controlling phenology and assortative mating.

### Z-linked genes associated with allochronic divergence

The identification of marked allelic differences between summer and winter populations at the Z chromosome, especially at genes associated with neurotransmission processes related to circadian rhythm regulation, is overall in line with empirical work in insects reporting Z-linked circadian genes associated with phenological variations (Kozak *et al*., 2019; Pruisscher *et al*., 2021; Zhang *et al*., 2022; Tessnow *et al*., 2022, 2025). In particular, the *NEP-2* gene, found in our highest-scoring BayPass window showing a large high *F*_ST_ window immediately after a putative structural variant, belongs to the family of synaptic membrane endopeptidases *neprilysin*, which are known to influence the activity of pigment dispersing factor - *PDF* (Isaac *et al*., 2007). Indeed, Isaac et al. (2007) showed in adults *Drosophila melanogaster* that neprilysin inactivates *PDF* through cleavage at the Serine7-Leucine8 peptide bond, resulting in greatly reduced signaling at the *PDF* -receptor encoded by *pdfr* (Lear *et al*., 2005). *PDF* is known to be a key neurotransmitter regulating rhythmic circadian locomotor behaviors by stabilizing the *period* /*timeless* heterodimer (Seluzicki *et al*., 2014) and by stimulating *Clk/CYC* -dependent transcription in a time-dependent manner (Sabado *et al*., 2017). In addition, *PDF* synchronizes organismal activity with the annual cycle of seasons via *pdfr*, with potential epistatic control of diapause termination occurring through the release of ecdysone, as discovered in the silkmoth *Bombyx mori* (Iga *et al*., 2014; Kozak *et al*., 2019). Interestingly, *PDF* abundance in small ventral lateral neurons (s-LNvs) of *Drosophila melanogaster* is lower on cold and short days compared to warm and long days, indicating an effect of both photoperiod and temperature on *PDF*, with the effect primarily attributed to temperature (Hidalgo *et al*., 2023; Hamanaka *et al*., 2024). Hence, our result may suggest a genetic polymorphism between LSP and LWP regulating the expression of *NEP-2* as a putative modifier of diapause timing, a hypothesis that requires further investigation. The potential role of *NEURL4* may be more indirect given that *NEURL4* has been shown to be involved in cellular metabolism regulation (Liu and Boulianne, 2017). Since it is located immediately upstream *NEP-2*, both genes could be considered as a functional module, with *NEURL4* delivering cellular energy delivered for *NEP-2* neprilysin production. Hence, a genetic polymorphism modifying the expression of *NEURL4*, or more likely of both *NEURL4* and *NEP-2*, in LSP could alter diapause timing. Alternatively, a genetic polymorphism within the *NEURL4* gene in LSP, modifying cellular energy regulation, could alter *NEP-2* neprilysin production and hence diapause timing. Other genes - *NPAS2*, *GABAR*, *Coq8* and *SP2353* - potentially involved in circadian or biological rhythm were also detected in significant, although lower, C2 local score windows. *NPAS2* is a paralog of the *Clk* gene, can functionally replace *Clk* to regulate circadian rhythms in mammals (DeBruyne *et al*., 2007; Peng *et al*., 2021). Furthermore, *NPAS2* pairs with *BMAL1* to form a heterodimer, and together they target the transcription of *period* and *CRY* genes, both crucial for regulating the circadian clock in mammals (Vitaterna *et al*., 1994).*GABA* receptors have been shown to affect diapause hormone levels in the silkworm *Bombyx mori* (Cui *et al*., 2021), and disrupting the *period* gene in silkworms influences feedback regulation of the *GABA* pathway by modifying the circadian clock pathway’s signal output. The *Coq8* gene was found to affect neuronal phenotypes in *Drosophila*, especially the development and maintenance of photoreceptors (Hura *et al*., 2022). The *SP2353* gene was found to cause abnormal photoreceptor axon pathfinding and/or differentiation phenotypes in *Drosophila* (Marrone *et al*., 2011). Lastly, the *period* gene, immediately upstream of a high C2 local score window in our study, is well known to influence seasonal timing in Lepidoptera (Sandrelli *et al*., 2007), notably affecting egg hatching (Ikeda *et al*., 2019, 2021; Nartey *et al*., 2021) and being associated with the induction of photoperiodic diapause as well as its termination (Kozak *et al*., 2019; Pruisscher *et al*., 2018, 2021; Nelson *et al*., 2022).

From an evolutionary perspective, the location on the Z chromosome of genetic polymorphisms putatively controlling biological rhythms may be particularly important. Indeed, given the reduced *N_e_*of sex chromosomes compared to autosomes (driven by heterogametic ZW females), a faster evolution of these chromosomes is predicted (*Fast-Z evolution* - Vicoso and Charlesworth (2006)). Faster-Z likely explains increased genetic differentiation and reduced diversity at the Z chromosome compared to autosomes, in all the populations of our study, as well as in other Lepidoptera (Meisel and Connallon, 2013; Mongue *et al*., 2022). However, the over-differentiation of the Z chromosome in the summer population was much higher relative to genome-wide differentiation than in the winter populations, indicating more differentiation in LSP than solely expected *via* the Faster-Z effect. The fact that *D*_XY_ values were low between summer and winter populations along the Z chromosome suggests that the over-differentiation Z of LSP arose recently from a common local gene pool. Importantly, the Z of LSP showed a nearly four-fold reduction of nucleotide diversity compared to autosomes, which was twice the nearly two-fold reduction observed in other populations, suggesting a dramatic bottleneck on the Z of LSP, causing its over-differentiation. Overall, the most parsimonious explanation of such reduced diversity and increased differentiation of the Z in LSP may be a strong founder effect caused by non-selective assortative mating driven by high effect Z-linked mutations, for example linked to *NEP-2*, underlying the phenological shift of LSP, together with heterogeneous gene flow along the genome (low gene flow at autosomes and almost null gene flow at the Z chromosome). In contrast, a role of divergent selection *a fortiori* as a trigger of allochrony seems unlikely given the small *N_e_* in LSP and in particular at the time of LSP split, especially so in sympatry. Nevertheless, after the onset of the allochronic divergence, during population growth after bottleneck, positive selection might have acted in LSP on several genes implicated in the several adaptations theoretically needed to fit its new phenology (Jarrett *et al*., 2025). This expectation might be in line with our finding of numerous genes associated to phenology, distributed across the Z chromosome and with large variations in c2 local scores, resembling a polygenic signature of response to positive selection in LSP. Lastly, a biased sex ratio towards females in LSP during its founding event could also have contributed to the reduction of genetic diversity on the Z of this population. This hypothesis is however difficult to test using genetic data given the strong genomic footprints of selection on the Z chromosome in LSP which likely drove an extreme reduction of genetic diversity at the entire Z.

Yet, the molecular nature, age, and contribution of each of the putative genetic variants associated with the allochronic shift in LSP require further investigation. In addition, analyzing INDELs, CNVs, and inversions for both LSP and LWP based on long read sequencing data (Wadsworth *et al*., 2015; Bastide *et al*., 2022) would help to characterize putative structural variants between and better infer point mutations. RNAseq or RNAi studies of the aforementioned genes throughout different life stages and seasons would also validate their potential roles. Simulations in SLIM (Haller and Messer, 2023) could also help to better understand the dynamics of the evolution of these Z-linked genetic variants associated with allochrony. Lastly, laboratory crosses between LSP and LWP and genotyping of recombinant families, although very challenging, would probably allow us to pinpoint more precisely loci underlying the phenological variations between both populations (Branco *et al*., 2017).

### Extent of inbreeding and genetic load in the allochronic summer population

Genome-wide inbreeding levels in LSP, LWP and CA were high compared to other populations sampled in Portugal, likely resulting from the aforementioned bottlenecks of higher intensity in the planted forests between Lisbon and the Mata Nacional de Leiria. These high inbreeding levels were comparable to those observed in critically endangered vertebrate species (Robinson *et al*., 2016, 2018; Grossen *et al*., 2020; Dussex *et al*., 2023; Kardos, 2023) or invasive insect species (Taylor *et al*., 2024a). The relatively short ROH segments found (mostly between 100 kb and 1 Mb, with very few exceeding 1 Mb) correspond to a size range commonly reported in Lepidoptera species with low *N_e_* or limited dispersal (Bortoluzzi *et al*., 2023). The difference in inbreeding levels between, on one hand, LSP, LWP, and CA and, on the other hand, the other populations, is comparable to what was found between populations of the Lepidoptera *Satyrium semiluna* (MacDonald *et al*., 2025), with different levels of spatial isolation. Overall, these results confirm a peculiar history of PPM coastal populations (represented here by LSP, LWP, and CA), which experienced stronger bottlenecks and stronger inbreeding compared to more inland Portuguese populations.

In each of the three populations for which we studied genetic load - LSP, LWP, FU - we found signals consistent with past selection acting on both the realized and masked load on autosomes, as reflected by lower proportions of derived homozygote and heterozygous genotypes at LOF sites compared to MIS sites, and to MIS sites compared to SYN sites. The fact that the masked load - measured through the proportion of heterozygous non-synonymous sites - decreased from FU to LWP to LSP is likely attributable to genetic drift, which reduces overall genetic diversity under strong bottlenecks. An opposite pattern was found for the realized load - estimated via the proportion of derived homozygote non-synonymous sites - which decreased from LSP to LWP to FU. This likely reflects that drift not only reduces diversity but also promotes fixation of deleterious alleles, especially in small or inbred populations. On the other hand, Rxy allowed us to assess the relative excess of deleterious alleles compared to a neutral expectation. The high Rxy values for LOF sites in LSP (compared to LWP and FU) and in LWP (compared to FU) suggest that drift alone is insufficient to explain these patterns. This overall signal of relative accumulation of deleterious mutations in LSP and LWP compared to FU, and in LSP compared to LWP, likely reflects that selection was less efficient in purging the genetic load in LSP and LWP compared to FU and in LSP compared to LWP. This pattern may reflect that effective population sizes in LSP and LWP became too small to sustain efficient purifying selection, leading to a reduced ability to eliminate deleterious variants. The greater accumulation of genetic load in LSP might be explained by even smaller *N_e_* and lower m compared to LWP. These results align with theoretical predictions that slightly deleterious variants (MIS) and masked genetic load tend to accumulate under relaxed purifying selection in small or bottlenecked populations (Ohta, 1992; Keller, 2002; Charlesworth and Eyre-Walker, 2007; Charlesworth, 2009). However, our results contrast with some empirical studies reporting a deficit of strongly deleterious mutations in small or inbred populations, particularly in endangered species or populations that have experienced severe bottlenecks (Grossen *et al*., 2020; Kleinman-Ruiz *et al*., 2022; Bourgeois *et al*., 2024; Taylor *et al*., 2024b; Lombaert *et al*., 2025), but align with other empirical studies showing no purging or accumulation of strongly deleterious mutations (Femerling *et al*., 2023; Zeitler *et al*., 2023). One may wonder whether the high proportion of homozygous derived genotypes at strongly deleterious sites could have reduced the fitness of LSP, potentially threatening the long-term persistence of this allochronic population. Answering this question may require rearing LSP and LWP in a common garden and inspecting proxies of individual fitness. In turn, the signal of genetic purge detected at strongly deleterious mutations on the Z chromosome in LSP only may be due to the aforementioned particularly strong bottleneck that might have occurred at this chromosome in LSP. This result is also in line with the high efficiency of purifying selection on strongly deleterious recessive mutations on sex chromosomes since they are hemizygous in ZW genotypes, as shown in (Rousselle *et al*., 2016).

## Conclusion

Several of our results are in line with the theoretical expectation that small *N_e_*, assortative mating between individuals sharing similar reproductive time, and polymorphisms at genes controlling for phenology may promote the onset of allochronic divergence (Devaux and Lande, 2008; Kunkel, 2023). In particular, we found evidence for recent strong bottlenecks in LSP and in LWP and in another nearby population, suggesting that a regional environmental driver was causative of the genetic drift and inbreeding that may then have allowed allochronic divergence in LSP. Strong genetic differentiation between LSP and LWP in the vicinity of circadian genes on the Z chromosome was probably underlying phenological differences and associated with strong assortative mating. The localization of these genetic polymorphisms on the Z chromosome has likely been an important factor facilitating the divergence between LSP and LWP, further limiting gene swamping from the winter populations in the summer population. Further work is required to characterize in depth the molecular nature, age, and role of mutations underlying allochrony in LSP.

## Data availability statement

All raw data (fastq files for Pool-Seq and Ind-Seq data) will be available in the SRA repository BioProject: PRJNA1269716 (publicly available upon paper publication). Scripts will be available at https://github.com/TanguyMuller/Genomic of Allochrony.

## Acknowledgments and funding

The authors wish to thank the ANR for a collaborative grant (LOADEXP; ANR-22-CE02-0009) to C.P. which covered half of sequencing costs and half PhD scholarship to T.M.; funding from INRAE to C.P. covering half of the sequencing costs and half of a PhD scholarship; the MGX-GENOMICS platform for sequencing; the Genotoul Bio-informatic platform. We are grateful to A. Ducasse, J. Moulin, R. Vitalis, V. Danneville, J.M. Gigleux, P. Doinjashvili, C. Robin, for helping with administrative tasks; P. Nouhaud, M. De Navascues, A. Fraimout, and M. Uhl for discussions regarding bioinformatic and statistical analyses; K. Ipekdal (Hacettepe University, Ankara, Türkiye), N. Rahim (Higher National School of Biotechnology Taoufik Khaznadar, Constantine, Algeria) and F. Goussard (URZF, Orléans, France) for sharing samples of T. wilkinsoni, T. bonjeani and T. pinivora, respectively, M.L. Ben Jamâa (Institut National Agronomique de Tunis, Tunisia) for samples from Tunisia, and J. Hodar (University of Granada, Spain) for samples from Spain.

## Notes

### Competing Interest Statement

The authors have declared no competing interest.

### Summary of Updates

minor revisions during the peer review process

