## Supplementary material for "Population genomics of incipient allochronic divergence in the Pine Processionary Moth": supp files

### 1 Supplementary Tables

| Sample ID | Study ID | Population | Taxon name | Stage | Locality | Sampling date | Tissue for extraction | Collected by |
|---|---|---|---|---|---|---|---|---|
| PP_Portugal_SP_pool_seq2015 | LSP_P | LSP | Thaumetopoea pityocampa | adult | Leiria | 2012 | head and thorax | Susana Rocha |
| PP_Portugal_SP_CKER00298_seq2024 | LSP_1 | LSP | Thaumetopoea pityocampa | adult | Leiria | 14/06/2012 | head and thorax | Susana Rocha |
| PP_Portugal_SP_CKER00299_seq2024 | LSP_2 | LSP | Thaumetopoea pityocampa | adult | Leiria | 14/06/2012 | head and thorax | Susana Rocha |
| PP_Portugal_SP_CKER00300_seq2024 | LSP_3 | LSP | Thaumetopoea pityocampa | adult | Leiria | 14/06/2012 | head and thorax | Susana Rocha |
| PP_Portugal_SP_CKER00301_seq2024 | LSP_4 | LSP | Thaumetopoea pityocampa | adult | Leiria | 14/06/2012 | head and thorax | Susana Rocha |
| PP_Portugal_SP_CKER00302_seq2024 | LSP_5 | LSP | Thaumetopoea pityocampa | adult | Leiria | 14/06/2012 | head and thorax | Susana Rocha |
| PP_Portugal_SP_CKER00303_seq2024 | LSP_6 | LSP | Thaumetopoea pityocampa | adult | Leiria | 14/06/2012 | head and thorax | Susana Rocha |
| PP_Portugal_SP_CKER00304_seq2024 | LSP_7 | LSP | Thaumetopoea pityocampa | adult | Leiria | 31/05/2012 | head and thorax | Susana Rocha |
| PP_Portugal_SP_CKER00371_seq2023 | LSP_8 | LSP | Thaumetopoea pityocampa | adult | Leiria | 14/06/2012 | head and thorax | Susana Rocha |
| PP_Portugal_SP_CKER00373_seq2023 | LSP_9 | LSP | Thaumetopoea pityocampa | adult | Leiria | 14/06/2012 | head and thorax | Susana Rocha |
| PP_Portugal_SP_CKER00379_seq2023 | LSP_10 | LSP | Thaumetopoea pityocampa | adult | Leiria | 14/06/2012 | head and thorax | Susana Rocha |
| PP_Portugal_SP_CKER00380_seq2022 | LSP_11 | LSP | Thaumetopoea pityocampa | adult | Leiria | 14/06/2012 | head and thorax | Susana Rocha |
| PP_Portugal_SP_CKER00399_seq2022 | LSP_12 | LSP | Thaumetopoea pityocampa | adult | Leiria | 14/06/2012 | head and thorax | Susana Rocha |
| PP_Portugal_SP_CKER00463_seq2022 | LSP_13 | LSP | Thaumetopoea pityocampa | adult | Leiria | 17/05/2012 | head and thorax | Susana Rocha |
| PP_Portugal_SP_CKER_F3_SP_seq2015 | LSP_14 | LSP | Thaumetopoea pityocampa | adult | Crossing | NA | head and thorax | Susana Rocha |
| PP_Portugal_SP_CKER_F4_SP_seq2015 | LSP_15 | LSP | Thaumetopoea pityocampa | adult | Crossing | NA | head and thorax | Susana Rocha |
| PP_Portugal_SP_CKER_M1_SP_seq2015 | LSP_16 | LSP | Thaumetopoea pityocampa | adult | Crossing | NA | head and thorax | Susana Rocha |
| PP_Portugal_SP_CKER_M2_SP_seq2015 | LSP_17 | LSP | Thaumetopoea pityocampa | adult | Crossing | NA | head and thorax | Susana Rocha |
| PP_Portugal_SPcr_CKER00589_seq2024 | LSP_18 | LSP | Thaumetopoea pityocampa | adult | Crossing | NA | head and thorax | Susana Rocha |
| PP_Portugal_SPcr_CKER00594_seq2024 | LSP_19 | LSP | Thaumetopoea pityocampa | adult | Crossing | NA | head and thorax | Susana Rocha |
| PP_Portugal_SPcr_CKER00603_seq2024 | LSP_20 | LSP | Thaumetopoea pityocampa | adult | Crossing | NA | head and thorax | Susana Rocha |
| PP_Portugal_SPcr_CKER00606_seq2024 | LSP_21 | LSP | Thaumetopoea pityocampa | adult | Crossing | NA | head and thorax | Susana Rocha |
| PP_Portugal_SPcr_CKER00613_seq2024 | LSP_22 | LSP | Thaumetopoea pityocampa | adult | Crossing | NA | head and thorax | Susana Rocha |
| PP_Portugal_SPcr_CKER00625_seq2024 | LSP_23 | LSP | Thaumetopoea pityocampa | adult | Crossing | NA | head and thorax | Susana Rocha |
| PP_Portugal_SPcr_CKER00630_seq2024 | LSP_24 | LSP | Thaumetopoea pityocampa | adult | Crossing | NA | head and thorax | Susana Rocha |
| PP_Portugal_SPcr_CKER00633_seq2024 | LSP_25 | LSP | Thaumetopoea pityocampa | adult | Crossing | NA | head and thorax | Susana Rocha |
| PP_Portugal_WP_pool_seq2015 | LWP_P | LWP | Thaumetopoea pityocampa | adult | Leiria | 2012 | head and thorax | Susana Rocha |
| PP_Portugal_WP_CKER00482_seq2023 | LWP_1 | LWP | Thaumetopoea pityocampa | adult | Leiria | 22/08/2012 | head and thorax | Susana Rocha |
| PP_Portugal_WP_CKER00486_seq2022 | LWP_2 | LWP | Thaumetopoea pityocampa | adult | Leiria | 08/08/2012 | head and thorax | Susana Rocha |
| PP_Portugal_WP_CKER00491_seq2022 | LWP_3 | LWP | Thaumetopoea pityocampa | adult | Leiria | 08/08/2012 | head and thorax | Susana Rocha |
| PP_Portugal_WP_CKER00503_seq2022 | LWP_4 | LWP | Thaumetopoea pityocampa | adult | Leiria | 08/08/2012 | head and thorax | Susana Rocha |
| PP_Portugal_WP_CKER00505_seq2022 | LWP_5 | LWP | Thaumetopoea pityocampa | adult | Leiria | 08/08/2012 | head and thorax | Susana Rocha |
| PP_Portugal_WP_CKER00508_seq2022 | LWP_6 | LWP | Thaumetopoea pityocampa | adult | Leiria | 08/08/2012 | head and thorax | Susana Rocha |
| PP_Portugal_WP_CKER00517_seq2024 | LWP_7 | LWP | Thaumetopoea pityocampa | adult | Leiria | 06/09/2012 | head and thorax | Susana Rocha |
| PP_Portugal_WP_CKER00518_seq2024 | LWP_8 | LWP | Thaumetopoea pityocampa | adult | Leiria | 08/08/2012 | head and thorax | Susana Rocha |
| PP_Portugal_WP_CKER00519_seq2024 | LWP_9 | LWP | Thaumetopoea pityocampa | adult | Leiria | 08/08/2012 | head and thorax | Susana Rocha |
| PP_Portugal_WP_CKER00520_seq2024 | LWP_10 | LWP | Thaumetopoea pityocampa | adult | Leiria | 08/08/2012 | head and thorax | Susana Rocha |
| PP_Portugal_WP_CKER00538_seq2022 | LWP_11 | LWP | Thaumetopoea pityocampa | adult | Leiria | 06/09/2012 | head and thorax | Susana Rocha |
| PP_Portugal_WP_CKER00540_seq2022 | LWP_12 | LWP | Thaumetopoea pityocampa | adult | Leiria | 30/07/2012 | head and thorax | Susana Rocha |
| PP_Portugal_WPcr_CKER00559_seq2024 | LWP_13 | LWP | Thaumetopoea pityocampa | adult | Crossing | 28/08/2009 | head and thorax | Susana Rocha |
| PP_Portugal_WPcr_CKER00561_seq2024 | LWP_14 | LWP | Thaumetopoea pityocampa | adult | Crossing | 06/08/2009 | head and thorax | Susana Rocha |
| PP_Portugal_WPcr_CKER00563_seq2024 | LWP_15 | LWP | Thaumetopoea pityocampa | adult | Crossing | 07/08/2009 | head and thorax | Susana Rocha |
| PP_Portugal_WPcr_CKER00596_seq2024 | LWP_16 | LWP | Thaumetopoea pityocampa | adult | Crossing | 13/07/2011 | head and thorax | Susana Rocha |
| PP_Portugal_WPcr_CKER00597_seq2024 | LWP_17 | LWP | Thaumetopoea pityocampa | adult | Crossing | 04/08/2011 | head and thorax | Susana Rocha |
| PP_Portugal_WPcr_CKER00598_seq2024 | LWP_18 | LWP | Thaumetopoea pityocampa | adult | Crossing | 07/08/2011 | head and thorax | Susana Rocha |
| PP_Portugal_Fundao_pool_seq2022 | FU_P | FU | Thaumetopoea pityocampa | adult | Fundão | 2012 | thorax | Susana Rocha |
| PP_Portugal_Fundao_CKER01105_seq2022 | FU_1 | FU | Thaumetopoea pityocampa | adult | Fundão | 13/08/2012 | thorax | Susana Rocha |
| PP_Portugal_Fundao_CKER01111_seq2023 | FU_2 | FU | Thaumetopoea pityocampa | adult | Fundão | 10/09/2012 | thorax | Susana Rocha |
| PP_Portugal_Fundao_CKER01113_seq2022 | FU_3 | FU | Thaumetopoea pityocampa | adult | Fundão | 13/08/2012 | thorax | Susana Rocha |
| PP_Portugal_Fundao_CKER01114_seq2023 | FU_4 | FU | Thaumetopoea pityocampa | adult | Fundão | 13/08/2012 | thorax | Susana Rocha |
| PP_Portugal_Fundao_CKER01171_seq2023 | FU_5 | FU | Thaumetopoea pityocampa | adult | Fundão | 30/07/2012 | thorax | Susana Rocha |
| PP_Portugal_Fundao_CKER01173_seq2023 | FU_6 | FU | Thaumetopoea pityocampa | adult | Fundão | 30/07/2012 | thorax | Susana Rocha |
| PP_Portugal_Fundao_CKER01174_seq2023 | FU_7 | FU | Thaumetopoea pityocampa | adult | Fundão | 30/07/2012 | thorax | Susana Rocha |
| PP_Portugal_Fundao_CKER01175_seq2023 | FU_8 | FU | Thaumetopoea pityocampa | adult | Fundão | 13/08/2012 | thorax | Susana Rocha |
| PP_Portugal_Fundao_CKER01178_seq2023 | FU_9 | FU | Thaumetopoea pityocampa | adult | Fundão | 13/08/2012 | thorax | Susana Rocha |
| PP_Portugal_Fundao_CKER01187_seq2023 | FU_10 | FU | Thaumetopoea pityocampa | adult | Fundão | 16/07/2012 | thorax | Susana Rocha |
| PP_Portugal_Viseu_pool_seq2022 | VI_P | VI | Thaumetopoea pityocampa | larvae | Viseu | 2002 | entire specimen | Helela Santos |
| PP_Portugal_Viseu_1230_viseu_seq2022 | VI_1 | VI | Thaumetopoea pityocampa | larvae | Viseu | XX/02/2002 | entire specimen | Helela Santos |
| PP_Portugal_Viseu_1235_viseu_seq2022 | VI_2 | VI | Thaumetopoea pityocampa | larvae | Viseu | XX/02/2002 | entire specimen | Helela Santos |
| PP_Portugal_Viseu_CKERnana1_seq2023 | VI_3 | VI | Thaumetopoea pityocampa | larvae | Viseu | XX/02/2002 | entire specimen | Helela Santos |
| PP_Portugal_Viseu_CKERnana2_seq2023 | VI_4 | VI | Thaumetopoea pityocampa | larvae | Viseu | XX/02/2002 | entire specimen | Helela Santos |
| PP_Portugal_Varges_pool_seq2022 | VA_P | VA | Thaumetopoea pityocampa | larvae | Varges | 2003/2004 | entire specimen | Susana Rocha |
| PP_Portugal_Varges_CKER01301_seq2023 | VA_1 | VA | Thaumetopoea pityocampa | larvae | Varges | 2003/2004 | entire specimen | Susana Rocha |
| PP_Portugal_Varges_CKER01302_seq2023 | VA_2 | VA | Thaumetopoea pityocampa | larvae | Varges | 2003/2004 | entire specimen | Susana Rocha |
| PP_Portugal_Varges_CKER01308_seq2022 | VA_3 | VA | Thaumetopoea pityocampa | larvae | Varges | 2003/2004 | entire specimen | Susana Rocha |
| PP_Portugal_Varges_CKER01309_seq2022 | VA_4 | VA | Thaumetopoea pityocampa | larvae | Varges | 2003/2004 | entire specimen | Susana Rocha |
| PP_Portugal_Caparica_pool_seq2022 | CA_P | CA | Thaumetopoea pityocampa | adult | Caparica | 2012 | thorax | Susana Rocha |
| PP_Portugal_Caparica_CKER01261_seq2022 | CA_1 | CA | Thaumetopoea pityocampa | adult | Caparica | 30/07/2012 | thorax | Susana Rocha |
| PP_Portugal_Caparica_CKER01262_seq2023 | CA_2 | CA | Thaumetopoea pityocampa | adult | Caparica | 27/08/2012 | thorax | Susana Rocha |
| PP_Portugal_Caparica_CKER01263_seq2023 | CA_3 | CA | Thaumetopoea pityocampa | adult | Caparica | 24/09/2012 | thorax | Susana Rocha |
| PP_Portugal_Caparica_CKER01269_seq2022 | CA_4 | CA | Thaumetopoea pityocampa | adult | Caparica | 27/08/2012 | thorax | Susana Rocha |
| PP_Portugal_Tavira_pool_seq2022 | TA_P | TA | Thaumetopoea pityocampa | adult | Tavira | 2012 | thorax | Susana Rocha |
| PP_Portugal_Tavira_CKER01120_seq2022 | TA_1 | TA | Thaumetopoea pityocampa | adult | Tavira | 27/06/2012 | thorax | Susana Rocha |
| PP_Portugal_Tavira_CKER01199_seq2023 | TA_2 | TA | Thaumetopoea pityocampa | adult | Tavira | 27/08/2012 | thorax | Susana Rocha |
| PP_Portugal_Tavira_CKER01200_seq2022 | TA_3 | TA | Thaumetopoea pityocampa | adult | Tavira | 27/08/2012 | thorax | Susana Rocha |
| PP_Portugal_Tavira_CKER01201_seq2023 | TA_4 | TA | Thaumetopoea pityocampa | adult | Tavira | 27/06/2012 | thorax | Susana Rocha |
| PP_Portugal_Grandola_pool_seq2022 | GR_P | GR | Thaumetopoea pityocampa | adult | Grândola | 2012 | thorax | Susana Rocha |
| PP_Portugal_Grandola_CKER01231_seq2022 | GR_1 | GR | Thaumetopoea pityocampa | adult | Grândola | 30/07/2012 | thorax | Susana Rocha |

*Continued on next page...*

| Sample ID | Study ID | Population | Taxon name | Stage | Locality | Sampling date | Tissue for extraction | Collected by |
|---|---|---|---|---|---|---|---|---|
| PP_Portugal_Grandola_CKER01232_seq2022 | GR_2 | GR | Thaumetopoea pityocampa | adult | Grândola | 30/07/2012 | thorax | Susana Rocha |
| PP_Portugal_Grandola_CKER01233_seq2023 | GR_3 | GR | Thaumetopoea pityocampa | adult | Grândola | 30/07/2012 | thorax | Susana Rocha |
| PP_Portugal_Grandola_CKER01235_seq2023 | GR_4 | GR | Thaumetopoea pityocampa | adult | Grândola | 30/07/2012 | thorax | Susana Rocha |
| PP_Portugal_F1_pool_seq2024 | F1_P | F1 | Thaumetopoea pityocampa | adult | Crossing | 2009/2011/2012 | head and thorax | Susana Rocha |
| PP_Portugal_F1_CKER00569_seq2024 | F1_1 | F1 | Thaumetopoea pityocampa | adult | Crossing | 23/06/2009 | head and thorax | Susana Rocha |
| PP_Portugal_F1_CKER00570_seq2024 | F1_2 | F1 | Thaumetopoea pityocampa | adult | Crossing | 24/06/2009 | head and thorax | Susana Rocha |
| PP_Portugal_F1_CKER00572_seq2024 | F1_3 | F1 | Thaumetopoea pityocampa | adult | Crossing | 22/06/2009 | head and thorax | Susana Rocha |
| PP_Portugal_F1_CKER00573_seq2024 | F1_4 | F1 | Thaumetopoea pityocampa | adult | Crossing | 29/06/2009 | head and thorax | Susana Rocha |
| PP_Portugal_F1_CKER00600_seq2024 | F1_5 | F1 | Thaumetopoea pityocampa | adult | Crossing | 27/06/2011 | head and thorax | Susana Rocha |
| PP_Portugal_F1_CKER00601_seq2024 | F1_6 | F1 | Thaumetopoea pityocampa | adult | Crossing | 29/06/2011 | head and thorax | Susana Rocha |
| PP_Portugal_F1_CKER00602_seq2024 | F1_7 | F1 | Thaumetopoea pityocampa | adult | Crossing | 04/07/2011 | head and thorax | Susana Rocha |
| PP_Portugal_F1_CKER00616_seq2024 | F1_8 | F1 | Thaumetopoea pityocampa | adult | Crossing | 27/04/2012 | head and thorax | Susana Rocha |
| PP_Portugal_F1_CKER00617_seq2024 | F1_9 | F1 | Thaumetopoea pityocampa | adult | Crossing | 28/04/2012 | head and thorax | Susana Rocha |
| PP_Portugal_LateSP_pool_seq2024 | LateLSP_P | LateLSP | Thaumetopoea pityocampa | adult | Leiria | 2012 | head and thorax | Susana Rocha |
| PP_Portugal_LateSP_CKER00473_seq2024 | LateLSP_1 | LateLSP | Thaumetopoea pityocampa | adult | Leiria | 06/09/2012 | head and thorax | Susana Rocha |
| PP_Portugal_LateSP_CKER00475_seq2024 | LateLSP_2 | LateLSP | Thaumetopoea pityocampa | adult | Leiria | 06/09/2012 | head and thorax | Susana Rocha |
| PP_Portugal_LateSP_CKER00477_seq2024 | LateLSP_3 | LateLSP | Thaumetopoea pityocampa | adult | Leiria | 20/09/2012 | head and thorax | Susana Rocha |
| PP_Portugal_LateSP_CKER00478_seq2024 | LateLSP_4 | LateLSP | Thaumetopoea pityocampa | adult | Leiria | 06/09/2012 | head and thorax | Susana Rocha |
| PP_Portugal_LateSP_CKER00479_seq2024 | LateLSP_5 | LateLSP | Thaumetopoea pityocampa | adult | Leiria | 27/07/2012 | head and thorax | Susana Rocha |
| PP_Portugal_LateSP_HS107_seq2024 | LateLSP_6 | LateLSP | Thaumetopoea pityocampa | adult | Crossing | NA | head and thorax | Susana Rocha |
| PP_Portugal_LateSP_HS180_seq2024 | LateLSP_7 | LateLSP | Thaumetopoea pityocampa | adult | Crossing | NA | head and thorax | Susana Rocha |
| PP_Portugal_LateSP_P016_seq2024 | LateLSP_8 | LateLSP | Thaumetopoea pityocampa | adult | Crossing | NA | head and thorax | Susana Rocha |
| PP_Espagne-bas_Cortijuela-1418m_CKER00939 | ES_1 | ES | Thaumetopoea pityocampa | adult | Cortijuela | 13/07/2012 | thorax | José A. Hodar |
| PP_Espagne-bas_Cortijuela-1418m_CKER00959 | ES_2 | ES | Thaumetopoea pityocampa | adult | Cortijuela | 30/07/2012 | thorax | José A. Hodar |
| PP_Maroc_RIF_CKER00688 | RI_1 | RI | Thaumetopoea pityocampa | larvae | RIF-Bab Barred | NA | thorax | Driss Ghaioule |
| PP_Maroc_RIF_CKER00689 | RI_2 | RI | Thaumetopoea pityocampa | larvae | RIF-Bab Barred | NA | thorax | Driss Ghaioule |
| PP_Maroc_Oukaimeden_CKER00656 | OK_1 | OK | Thaumetopoea pityocampa | larvae | Oukaimeden | NA | thorax | Jérome Rousselet |
| PP_Maroc_Oukaimeden_CKER00683 | OK_2 | OK | Thaumetopoea pityocampa | larvae | Oukaimeden | NA | thorax | Jérome Rousselet |
| PP_Tunisie_Tunis_CKER00690 | TU_1 | TU | Thaumetopoea pityocampa | larvae | Tunis | NA | thorax | Jérome Rousselet |
| PP_Tunisie_Tunis_CKER00691 | TU_2 | TU | Thaumetopoea pityocampa | larvae | Tunis | NA | thorax | Jérome Rousselet |
| Bonjeani_Bonjeani_Tikjda_CKER00704 | BO_1 | BO | Thaumetopoea bonjeani | larvae | Tikjda | NA | thorax | NA |
| Bonjeani_Bonjeani_Tikjda_CKER00705 | BO_2 | BO | Thaumetopoea bonjeani | larvae | Tikjda | NA | thorax | NA |
| Pinivora_Pinivora_Alpes_CKER00668 | PI_1 | PI | Thaumetopoea pinivora | larvae | Alpes | NA | thorax | Jérome Rousselet |
| Pinivora_Pinivora_Alpes_CKER00677 | PI_2 | PI | Thaumetopoea pinivora | larvae | Alpes | NA | thorax | Jérome Rousselet |
| Wilkinsoni_Wilkinsoni_Turquie_CKER00146 | WI_1 | WI | Thaumetopoea wilkinsoni | NA | Mudanya | XX/03/2011 | thorax | Karaman Ipekdal |
| Wilkinsoni_Wilkinsoni_Turquie_CKER00213 | WI_2 | WI | Thaumetopoea wilkinsoni | NA | Istanbul | XX/03/2011 | thorax | Karaman Ipekdal |

Table S1: **Description of the data for the 90 individuals, 10 pools, and 14 outgroups used in this study.** Sample identifier, study identifier, putative species, developmental stage, geographical origin, sampling date, tissue used for DNA extraction, and collector information for each sample of this study. Individuals not obtained from field sampling but from laboratory rearing are labeled as "Crossing" in the locality column. For pool sampling, multiple nests were collected over a period, and one individual per nest was selected to avoid sampling siblings. Pools LSP, LWP, FU, CA, TA, GR and LateLSP were composed of adults captured using pheromone traps, while pools VI and VA consisted of field-collected caterpillars, and pool F1 included adults resulting from laboratory crossings.

| Study ID | data | Reads filtered (in %) | Duplicate reads (in %) | Read properly paired (in %) | Mean insert size (in bp) | Mean depth (in X) | GC (in %) | Depth mean ratio | Depth median ratio | Sex |
|---|---|---|---|---|---|---|---|---|---|---|
| LSP_P | pool | 3,228 | 3,697 | 97 | 241,9 | 47,015 | 37,753 | 1,009 | 1 | - |
| LSP_1 | individus | 2,262 | 6,935 | 96,8 | 333,7 | 16,659 | 37,552 | 1,002 | 1 | M |
| LSP_2 | individus | 2,466 | 5,86 | 96,9 | 317,8 | 14,56 | 37,621 | 1,054 | 1,071 | M |
| LSP_3 | individus | 2,461 | 6,389 | 96,6 | 322,6 | 17,337 | 37,588 | 1,039 | 1 | M |
| LSP_4 | individus | 2,46 | 6,181 | 96,4 | 337,1 | 15,729 | 37,564 | 1,005 | 1 | M |
| LSP_5 | individus | 3,372 | 6,726 | 96,7 | 317,7 | 19,204 | 37,504 | 1,035 | 1 | M |
| LSP_6 | individus | 2,542 | 6,721 | 96,6 | 340 | 18,602 | 37,586 | 1,004 | 1 | M |
| LSP_7 | individus | 3,114 | 5,877 | 97 | 330 | 17,91 | 37,617 | 1,005 | 1 | M |
| LSP_8 | individus | 1,37 | 13,624 | 97,8 | 372,6 | 73,125 | 37,518 | 1,017 | 1,013 | M |
| LSP_9 | individus | 1,046 | 10,667 | 97,8 | 366,7 | 25,737 | 37,812 | 1,012 | 1,04 | M |
| LSP_10 | individus | 0,955 | 9,739 | 97,9 | 363 | 24,838 | 37,595 | 1,014 | 1 | M |
| LSP_11 | individus | 1,519 | 7,316 | 97,7 | 362,3 | 30,509 | 37,714 | 1,007 | 0,968 | M |
| LSP_12 | individus | 1,426 | 6,386 | 97,7 | 358,4 | 32,011 | 37,797 | 1,037 | 1,031 | M |
| LSP_13 | individus | 1,648 | 7,869 | 97,6 | 371,9 | 32,787 | 37,747 | 1,009 | 1 | M |
| LSP_14 | individus | 0,234 | 2,056 | 97,3 | 439,3 | 28,766 | 37,683 | 0,514 | 0,483 | F |
| LSP_15 | individus | 0,231 | 2,153 | 97,3 | 422 | 28,892 | 37,782 | 0,513 | 0,483 | F |
| LSP_16 | individus | 0,226 | 1,967 | 97,2 | 450,9 | 28,174 | 37,378 | 1,007 | 1 | M |
| LSP_17 | individus | 0,237 | 2,029 | 97,4 | 436,7 | 32,849 | 37,613 | 1,002 | 0,97 | M |
| LSP_18 | individus | 3,134 | 5,787 | 97 | 287,5 | 17,497 | 37,462 | 0,506 | 0,471 | F |
| LSP_19 | individus | 2,87 | 6,437 | 96,7 | 300,6 | 17,916 | 37,305 | 0,513 | 0,5 | F |
| LSP_20 | individus | 4,167 | 6,696 | 97,3 | 283 | 22,683 | 37,627 | 0,525 | 0,478 | F |
| LSP_21 | individus | 4,636 | 6,524 | 96,6 | 249,7 | 17,366 | 37,45 | 0,52 | 0,5 | F |
| LSP_22 | individus | 3,013 | 6,801 | 97,4 | 309,7 | 23,004 | 37,485 | 0,52 | 0,5 | F |
| LSP_23 | individus | 17,819 | 5,131 | 97,8 | 259 | 16,004 | 38,396 | 0,518 | 0,467 | F |
| LSP_24 | individus | 7,327 | 6,738 | 97,5 | 279 | 15,735 | 37,758 | 0,515 | 0,467 | F |
| LSP_25 | individus | 3,886 | 6,515 | 96,7 | 268 | 16,727 | 37,598 | 0,512 | 0,5 | F |
| LWP_P | pool | 2,631 | 3,481 | 97,5 | 226,9 | 43,548 | 37,818 | 1,015 | 1 | - |
| LWP_1 | individus | 1,532 | 12,622 | 97,7 | 361,1 | 56,364 | 37,8 | 1,013 | 1 | M |
| LWP_2 | individus | 1,59 | 6,585 | 97,5 | 357,5 | 31,359 | 37,695 | 1,056 | 1,032 | M |
| LWP_3 | individus | 1,618 | 7,325 | 97,3 | 373,7 | 39,983 | 37,575 | 1,032 | 1,025 | M |
| LWP_4 | individus | 1,518 | 6,891 | 97,7 | 365,1 | 46,329 | 37,672 | 1,022 | 1 | M |
| LWP_5 | individus | 1,251 | 6,438 | 97,6 | 373,3 | 32,348 | 37,693 | 1,019 | 1 | M |
| LWP_6 | individus | 1,429 | 6,225 | 97,5 | 356,8 | 35,2 | 37,703 | 1,023 | 1 | M |
| LWP_7 | individus | 3,725 | 6,186 | 95,7 | 257,5 | 20,596 | 37,468 | 1,007 | 1 | M |
| LWP_8 | individus | 3,197 | 6,256 | 96,9 | 301,6 | 18,17 | 37,044 | 1,015 | 1 | M |
| LWP_9 | individus | 3,162 | 6,156 | 96,7 | 328,3 | 19,164 | 37,678 | 1,018 | 1 | M |
| LWP_10 | individus | 3,958 | 5,952 | 96,3 | 296,7 | 17,416 | 37,799 | 1,048 | 1,062 | M |
| LWP_11 | individus | 1,767 | 6,583 | 97,5 | 356,2 | 29,411 | 37,737 | 1,024 | 1 | M |
| LWP_12 | individus | 1,493 | 7,104 | 97,6 | 371,8 | 36,224 | 37,716 | 1,016 | 1 | M |
| LWP_13 | individus | 3,497 | 5,804 | 96,9 | 314,7 | 18,574 | 37,414 | 0,526 | 0,474 | F |
| LWP_14 | individus | 5,409 | 5,63 | 97,5 | 272,2 | 19,462 | 37,73 | 0,525 | 0,5 | F |
| LWP_15 | individus | 5,007 | 6,494 | 97,2 | 249,6 | 20,346 | 37,558 | 0,543 | 0,5 | F |
| LWP_16 | individus | 6,902 | 6,637 | 98 | 249 | 25,561 | 37,844 | 0,54 | 0,48 | F |
| LWP_17 | individus | 4,854 | 7,527 | 97,1 | 265,8 | 21,752 | 38,298 | 0,533 | 0,476 | F |
| LWP_18 | individus | 2,597 | 6,093 | 97,4 | 297,9 | 21,942 | 37,806 | 0,547 | 0,476 | F |
| FU_P | pool | 1,324 | 10,941 | 97,7 | 326,3 | 52,479 | 37,656 | 1,027 | 1 | - |
| FU_1 | individus | 1,818 | 8,652 | 97,6 | 335 | 34,463 | 37,794 | 1,028 | 1,029 | M |
| FU_2 | individus | 0,806 | 11,43 | 97,3 | 399 | 22,219 | 37,205 | 1,006 | 1 | M |
| FU_3 | individus | 1,27 | 5,837 | 97,7 | 322,5 | 21,859 | 37,859 | 1,01 | 0,955 | M |
| FU_4 | individus | 0,97 | 7,562 | 97,9 | 368,4 | 36,105 | 37,481 | 1,026 | 1 | M |
| FU_5 | individus | 1,055 | 5,326 | 98 | 361 | 22,802 | 37,342 | 1,031 | 1 | M |
| FU_6 | individus | 1,049 | 11,147 | 97,4 | 377,5 | 24,267 | 37,388 | 1,032 | 1 | M |
| FU_7 | individus | 0,924 | 13,397 | 97,7 | 374,2 | 24,159 | 37,456 | 1,029 | 1 | M |
| FU_8 | individus | 0,965 | 5,914 | 97,6 | 392,2 | 21,875 | 37,427 | 1,016 | 1 | M |
| FU_9 | individus | 1,379 | 11,142 | 97,6 | 370,1 | 23,684 | 37,556 | 1,031 | 1,043 | M |
| FU_10 | individus | 0,977 | 6,145 | 97,8 | 395,4 | 29,154 | 37,414 | 1,027 | 1 | M |
| VI_P | pool | 1,448 | 12,537 | 97,9 | 345,5 | 58,08 | 37,715 | 0,835 | 0,817 | - |
| VI_1 | individus | 1,295 | 9,143 | 97,8 | 348 | 24,237 | 37,901 | 0,541 | 0,5 | F |
| VI_2 | individus | 1,218 | 10,521 | 97,5 | 345,4 | 25,095 | 37,744 | 1,033 | 1 | M |
| VI_3 | individus | 1,049 | 15,668 | 97,3 | 402,1 | 30,378 | 37,588 | 0,533 | 0,484 | F |
| VI_4 | individus | 1,341 | 13,896 | 97,7 | 383,6 | 22,851 | 37,526 | 0,535 | 0,522 | F |
| VA_P | pool | 1,476 | 11,884 | 97,5 | 349,3 | 47,913 | 37,673 | 0,737 | 0,714 | - |
| VA_1 | individus | 1,427 | 12,451 | 97,6 | 405,5 | 26,76 | 37,4 | 0,53 | 0,481 | F |
| VA_2 | individus | 0,991 | 13,507 | 97,4 | 396,6 | 24,647 | 37,317 | 0,526 | 0,48 | F |
| VA_3 | individus | 1,83 | 10,203 | 97,8 | 360,6 | 21,211 | 37,626 | 0,531 | 0,5 | F |
| VA_4 | individus | 1,576 | 8,368 | 97,8 | 360,9 | 20,387 | 37,754 | 0,532 | 0,5 | F |
| CA_P | pool | 1,295 | 9,621 | 97,8 | 329,6 | 55,204 | 38,524 | 1,022 | 1 | - |
| CA_1 | individus | 1,318 | 7,279 | 97,7 | 337,4 | 27,513 | 37,791 | 1,026 | 1 | M |
| CA_2 | individus | 0,769 | 12,458 | 97,3 | 389,6 | 29,155 | 37,115 | 1,004 | 0,967 | M |
| CA_3 | individus | 0,955 | 12,195 | 97,4 | 396,2 | 27,743 | 37,434 | 1,029 | 1 | M |
| CA_4 | individus | 1,23 | 6,295 | 97,8 | 337,4 | 26,628 | 37,887 | 1,022 | 1 | M |
| TA_P | pool | 1,468 | 7,94 | 97,7 | 324,6 | 52,825 | 37,739 | 1,021 | 1,019 | - |
| TA_1 | individus | 1,229 | 6,733 | 97,4 | 333,7 | 25,796 | 37,804 | 1,014 | 1 | M |
| TA_2 | individus | 0,902 | 12,5 | 97,4 | 390,3 | 25,421 | 37,306 | 1,02 | 1 | M |
| TA_3 | individus | 1,394 | 5,724 | 97,3 | 322,1 | 22,179 | 37,614 | 1,008 | 1 | M |

*Continued on next page...*

| Study ID | data | Reads filtered (in %) | Duplicate reads (in %) | Read properly paired (in %) | Mean insert size (in bp) | Mean depth (in X) | GC (in %) | Depth mean ratio | Depth median ratio | Sex |
|---|---|---|---|---|---|---|---|---|---|---|
| TA_4 | individus | 0,844 | 11,461 | 97,6 | 389,7 | 24,531 | 37,268 | 1,024 | 1 | M |
| GR_P | pool | 1,604 | 7,535 | 97,9 | 312,3 | 50,327 | 37,682 | 1,022 | 1 | - |
| GR_1 | individus | 1,368 | 7,285 | 97,2 | 326,9 | 29,829 | 37,633 | 1,021 | 1 | M |
| GR_2 | individus | 1,471 | 5,492 | 97,1 | 325,3 | 22,555 | 38,401 | 1,006 | 1 | M |
| GR_3 | individus | 0,842 | 13,091 | 97,4 | 387,3 | 30,051 | 37,434 | 1,028 | 1,033 | M |
| GR_4 | individus | 0,885 | 12,104 | 97,7 | 385,3 | 28,576 | 37,463 | 1,02 | 1 | M |
| F1_P | pool | 4,775 | 8,397 | 97,5 | 281,8 | 62,665 | 37,436 | 0,769 | 0,758 | - |
| F1_1 | individus | 2,756 | 5,955 | 97,1 | 313,3 | 18,482 | 37,27 | 1,017 | 1 | M |
| F1_2 | individus | 6,069 | 7,696 | 97,2 | 235,1 | 23,321 | 37,029 | 0,525 | 0,5 | F |
| F1_3 | individus | 2,506 | 7,765 | 97,2 | 323,6 | 21,515 | 37,459 | 1,013 | 1 | M |
| F1_4 | individus | 3,865 | 6,259 | 97 | 263,9 | 17,972 | 37,317 | 0,536 | 0,5 | F |
| F1_5 | individus | 3,453 | 5,86 | 96,9 | 290,7 | 24,826 | 37,477 | 1,014 | 1 | M |
| F1_6 | individus | 5,453 | 6,888 | 97,2 | 245,7 | 20,555 | 37,838 | 1,012 | 1 | M |
| F1_7 | individus | 3,632 | 6,35 | 97,8 | 281,1 | 16,076 | 37,69 | 0,527 | 0,5 | F |
| F1_8 | individus | 3,223 | 6,587 | 97,3 | 318 | 25,166 | 37,481 | 1,009 | 1 | M |
| F1_9 | individus | 2,602 | 6,539 | 96,9 | 299 | 18,049 | 37,495 | 0,528 | 0,5 | F |
| LateLSP_P | pool | 3,1 | 7,644 | 97,6 | 281,2 | 56,381 | 38,572 | 0,989 | 0,983 | - |
| LateLSP_1 | individus | 3,31 | 6,175 | 97,2 | 297,2 | 15,981 | 39,322 | 1,009 | 1 | M |
| LateLSP_2 | individus | 2,47 | 7,042 | 96,9 | 313,4 | 20,243 | 37,636 | 0,998 | 1 | M |
| LateLSP_3 | individus | 2,585 | 6,117 | 96,7 | 333,5 | 19,203 | 37,461 | 1,01 | 1 | M |
| LateLSP_4 | individus | 2,847 | 5,563 | 96,9 | 334 | 19,432 | 37,564 | 1,009 | 1 | M |
| LateLSP_5 | individus | 3,961 | 6,371 | 96,5 | 300,2 | 29,062 | 37,597 | 1 | 1 | M |
| LateLSP_6 | individus | 3,642 | 6,482 | 96,4 | 311,1 | 22,402 | 37,269 | 1,007 | 1 | M |
| LateLSP_7 | individus | 4,716 | 6,457 | 97,3 | 294,2 | 18,041 | 38,247 | 0,547 | 0,5 | F |
| LateLSP_8 | individus | 2,088 | 6,448 | 96,4 | 352,7 | 16,383 | 37,126 | 1,008 | 1 | M |
| ES_1 | outgroups | 2,574 | 7,066 | 97,4 | 373,9 | 28,242 | 38,33 | 1,021 | 1 | M |
| ES_2 | outgroups | 1,003 | 12,639 | 97,9 | 362,6 | 48,159 | 37,575 | 1,014 | 1 | M |
| RI_1 | outgroups | 1,355 | 11,376 | 96 | 455,5 | 25,414 | 37,151 | 1,04 | 1 | M |
| RI_2 | outgroups | 1,34 | 13,225 | 96 | 474,3 | 25,199 | 37,321 | 0,566 | 0,48 | F |
| OK_1 | outgroups | 1,284 | 1,443 | 95,6 | 462,7 | 24,079 | 37,423 | 0,997 | 1 | M |
| OK_2 | outgroups | 1,197 | 13,954 | 95,7 | 437,6 | 29,337 | 37,273 | 1,013 | 1,034 | M |
| TU_1 | outgroups | 1,147 | 10,78 | 94,9 | 422,7 | 22,599 | 37,493 | 1,005 | 1 | M |
| TU_2 | outgroups | 1,224 | 13,328 | 95,2 | 423,7 | 20,011 | 37,518 | 0,517 | 0,45 | F |
| BO_1 | outgroups | 0,978 | 12,7 | 82,1 | 361,9 | 22,448 | 37,156 | 1,001 | 1 | M |
| BO_2 | outgroups | 0,94 | 11,391 | 81,1 | 396,7 | 22,387 | 37,046 | 0,975 | 0,958 | M |
| PI_1 | outgroups | 0,999 | 14,156 | 81 | 390,8 | 20,41 | 37,386 | 1,251 | 1,05 | M |
| PI_2 | outgroups | 1,462 | 12,112 | 78,2 | 497,5 | 18,58 | 37,155 | 1,171 | 1 | M |
| WI_1 | outgroups | 1,001 | 16,189 | 94 | 402,8 | 27,368 | 37,53 | 1,035 | 1 | M |
| WI_2 | outgroups | 1,004 | 13,894 | 93,9 | 379,9 | 26,58 | 37,961 | 0,589 | 0,5 | F |

Table S2: **Information about quality of the data for the 90 individuals, 10 pools, and 14 outgroups used in this study**. All libraries were paired-end (150 bp) sequenced on the MGX-Montpellier Genomix platform using a NovaSeq6000, with the exception of 4 individual libraries sequenced on an Illumina HiSeq2500 (LSP_14, LSP_15, LSP_16 and LSP_17). The percentage of filtered reads was obtained after processing the raw paired-end reads with fastp 0.21.0 (Chen *et al.*, 2018) using the default settings. The duplicate reads were estimated after filtering of raw paired-end reads. Filtered reads were then mapped onto the *Tpit2.1* chromosome level reference genome (Gautier et al. in prep) using the *bwa-mem2 2.2.1* program (Vasimuddin *et al.*, 2019) with default options. Read alignments with a mapping quality Phred-score <20 or PCR duplicates were removed using *SAMtools 1.14* software (Danecek *et al.*, 2021) with the view (option *-q 20*) and the markdup program. The percentage of reads mapped and properly paired, mean insert size, mean depth and percentage of GC was obtained from BAM files using option *stats* of *Samtools 1.14*. Genotypic sex of individuals, pools and outgroup was inferred by comparing the median and mean coverage of the Z chromosome (28.98 Mb) with that of the largest autosome (chromosome 1, 26.55 Mb). For sexing individuals, the males were identified by similar coverage for both chromosomes 1 and Z (i.e. ratio near of 1), while the heterogametic females were identified by half coverage on chromosome Z compared to chromosome 1 (i.e. ratio near of 0.5). To identify the number of males and females in each pool, the ratio of the median coverage of Z to autosome is expected to be $\rho = (2nm + nf)/(2n)$, where $nm$ is the number of males, $nf$ is the number of females and $n = nm + nf$. Using this ratio, the estimated haploid size of the pool was calculated as $\zeta = (1-\rho) * 2n * 1 + (n - (1-\rho) * 2n) * 2$, where $(1-\rho) * 2n * 1$ represents the number of copies contributed by females and $(n - (1-\rho) * 2n) * 2$ represents the number of copies contributed by males, giving the number of males and females for the pool.

| Summary statistic | Data | Software use |
|---|---|---|
| $F_{\mathrm{ST}}$ | pool | *computeFST* from *poolfstat* |
| $F_{\mathrm{ST}}$ LSP∼LWP | pool | *compute.fstat* from *poolfstat* |
| $F_{\mathrm{ST}}$ LSP∼FU | pool | *compute.fstat* from *poolfstat* |
| $F_{\mathrm{ST}}$ LWP∼FU | pool | *compute.fstat* from *poolfstat* |
| HET LSP | pool | *compute.fstat* from *poolfstat* |
| HET LSP | individual | *–hardy* from *VCFtools* |
| HET LWP | pool | *compute.fstat* from *poolfstat* |
| HET LWP | individual | *–hardy* from *VCFtools* |
| HET FU | pool | *compute.fstat* from *poolfstat* |
| HET FU | individual | *–hardy* from *VCFtools* |
| *f2* LSP∼LWP | pool | *compute.fstat* from *poolfstat* |
| *f2* LSP∼FU | pool | *compute.fstat* from *poolfstat* |
| *f2* LWP∼FU | pool | *compute.fstat* from *poolfstat* |
| *f3* LSP | pool | *compute.fstat* from *poolfstat* |
| *f3* LWP | pool | *compute.fstat* from *poolfstat* |
| *f3* FU | pool | *compute.fstat* from *poolfstat* |
| Pfix in LSP | pool | *poolfstat* |
| Pfix in LWP | pool | *poolfstat* |
| Pfix in FU | pool | *poolfstat* |
| Pfix ref in LSP | pool | *poolfstat* |
| Pfix ref in LWP | pool | *poolfstat* |
| Pfix ref in FU | pool | *poolfstat* |
| Pfix alt in LSP | pool | *poolfstat* |
| Pfix alt in LWP | pool | *poolfstat* |
| Pfix alt in FU | pool | *poolfstat* |
| D Tajima LSP | individual | *–TajimaD 10000* from *VCFtools* |
| D Tajima LWP | individual | *–TajimaD 10000* from *VCFtools* |
| D Tajima FU | individual | *–TajimaD 10000* from *VCFtools* |
| Rs LSP LWP | individual | In-house python script |
| Rs LSP FU | individual | In-house python script |
| Rs LWP FU | individual | In-house python script |
| Rf LSP LWP | individual | In-house python script |
| Rf LSP FU | individual | In-house python script |
| Rf LWP FU | individual | In-house python script |
| Wx | individual | In-house python script |
| Wxn | individual | In-house python script |
| WF | individual | In-house python script |
| WFn | individual | In-house python script |
| LSP SFS 0 to 50 | individual | *–counts* from *VCFtools* |
| LWP SFS 0 to 36 | individual | *–counts* from *VCFtools* |
| FU SFS 0 to 20 | individual | *–counts* from *VCFtools* |

Table S3: **All summary statistics used for ABCRF**. A total of 158 summary statistics were used for ABCRF analyses on individual and pool data using poolfstat 3.0.0 (Gautier *et al.*, 2024) and VCFtools 0.1.16 (Danecek *et al.*, 2011) $F_{ST}$ : $F_{ST}$ estimate for all the three population and for each pair of populations. HET : Heterozygosity for each population calculted on individual and pool data. $f$ : allele-shared f-statistics from Patterson *et al.* (2012) computed for each pair ($f2$) and triplet ($f3$). Pfix : Proportion of monomorphic loci within each population calculated on pool. Additionally, we decompose this proportion into two components: the proportion of monomorphic loci for the reference allele and the proportion for the alternate allele. D Tajima : Tajima's D statistic in bins with size of 10,000 bp for each population. The following 10 summary statistics are produced by an in-house Python script that enhances the statistics described in Navascués *et al.* (2014). Rf : Runs of polymorphic sites found in the alignment fixed differences. Rs : Runs of polymorphic sites found in the alignment shared polymorphisms. Wx : Proportion of fixed site in one population but polymorphic in the other population. Wxn : Proportion of fixed site in one population but polymorphic in the other population taking only polymorphic site. WF : Proportion of fixed site in one population but fixed differentially in the other. WFn : Proportion of fixed site in one population but fixed differentially in the other taking only polymorphic site. SFS: The proportions of each allele class ranging from 0 to 2n, where n is the number of individuals sequenced for a given population.

| | | concomitent_div | | recent_div | | old_div | |
|---|---|---|---|---|---|---|---|
| **Parameter** | **Distribution** | **Minimum** | **Maximum** | **Minimum** | **Maximum** | **Minimum** | **Maximum** |
| Ne_anc_PP | log-uniform | 100000 | 1000000 | 100000 | 1000000 | 100000 | 1000000 |
| Ne_LSP | log-uniform | 100 | 100000 | 100 | 100000 | 100 | 100000 |
| Ne_LWP | log-uniform | 100 | 10000 | 100 | 10000 | 100 | 10000 |
| Ne_FU | log-uniform | 100 | 200000 | 100 | 200000 | 100 | 200000 |
| Ne_Old_LSP | log-uniform | - | - | - | - | 100 | 100000 |
| Ne_Old_LWP | log-uniform | - | - | 100 | 100000 | 100 | 100000 |
| Ne_Old_FU | log-uniform | - | - | 100 | 200000 | - | - |
| Ne_LSP_found | log-uniform | 1 | Ne_Anc_PP | 1 | Ne_Old_LWP | 1 | Ne_Anc_PP |
| Ne_LWP_found | log-uniform | 1 | Ne_Anc_PP | 1 | Ne_Old_LWP | 1 | Ne_Old_LWP |
| Ne_FU_found | log-uniform | 100 | Ne_Anc_PP | 100 | Ne_Anc_PP | 100 | Ne_Old_LWP |
| Ne_LSP_ancfound | log-uniform | - | - | - | - | 1 | Ne_Anc_PP |
| Ne_LWP_ancfound | log-uniform | - | - | 100 | Ne_Anc_PP | 100 | Ne_Anc_PP |
| Ne_FU_ancfound | log-uniform | - | - | 100 | Ne_Anc_PP | 100 | Ne_Anc_PP |
| Ne_bot_LSP | log-uniform | 1 | Ne_LSP | 1 | Ne_LSP | 1 | Ne_Old_LSP |
| Ne_bot_LWP | log-uniform | 1 | Ne_LWP | 1 | Ne_LWP | 1 | Ne_LWP |
| Ne_bot_FU | log-uniform | 1 | Ne_FU | 1 | Ne_Old$_{F}U$ | 1 | Ne_FU |
| Tsplit_PP | log-uniform | 500 | 20000 | 500 | 20000 | 500 | 20000 |
| Tsplit_WP | log-uniform | - | - | 11 | Tsplit_PP-200 if Tspilt_PP<2000<br>Tsplit_PP-500 if Tsplit_PP>2000 | 11 | Tsplit_PP-200 if Tsplitt_PP<2000<br>Tsplit_PP-500 if Tsplit_PP>2000 |
| Tbot | log-uniform | 10 | t1 | 10 | t2 | 10 | t2 |
| $F_{ST}$ for m_anc_LSPLWP* | log-uniform | 0.001 | 0.5 | 0.001 | 0.5 | 0.001 | 0.5 |
| $F_{ST}$ for m_rec_LSPLWP* | log-uniform | 0.001 | 0.5 | 0.001 | 0.5 | 0.001 | 0.5 |
| $F_{ST}$ for m_anc_LSPFU* | log-uniform | 0.001 | 0.5 | 0.001 | 0.5 | 0.001 | 0.5 |
| $F_{ST}$ for m_rec_LSPFU* | log-uniform | 0.001 | 0.5 | 0.001 | 0.5 | 0.001 | 0.5 |
| $F_{ST}$ for m_anc_LWPFU* | log-uniform | 0.001 | 0.5 | 0.001 | 0.5 | 0.001 | 0.5 |
| $F_{ST}$ for m_rec_LWPFU* | log-uniform | 0.001 | 0.5 | 0.001 | 0.5 | 0.001 | 0.5 |
| $F_{ST}$ for m_ancPP* | log-uniform | - | - | 0.001 | 0.5 | 0.001 | 0.5 |

Table S4: **Prior distribution of each parameter for the three scenarios**. The parameters are presented for the 3 scenarios. Ne_Anc_PP : ancestral population size; Ne_LSP : size of the actual LSP population; Ne_LWP : size of the actual LWP population; Ne_FU : size of the actual FU population; Ne_Old_LSP : size of the LSP population before the bottleneck when LSP is in the oldest divergence; Ne_Old_LWP : size of the LWP population before the split; Ne_Old_FU : size of the FU population before the bottleneck when FU is in the oldest divergence; Ne_LSP_found : size of the LSP population at the time of its founder event; Ne_LWP_found : size of the LWP population at the time of its founder event; Ne_FU_found : size of the FU population at the time of its founder event; Ne_LSP_ancfound : size of the founder LSP population at the time of the oldest divergence; Ne_LWP_ancfound : size of the founder LWP population at the time of the oldest divergence; Ne_FU_ancfound : size of the founder FU population at the time of the oldest divergence; Ne_bot_LSP : size of the LSP population at the time of the bottleneck; Ne_bot_LWP : size of the LWP population at the time of the bottleneck; Ne_bot_FU : size of the FU population at the time of the bottleneck; Tsplit_PP : time of the oldest divergence; Tsplit_WP : time of the most recent split; Tbot : time of the bottleneck event; m_anc_LSPLWP : symmetric migration between LSP and LWP before the bottleneck; m_rec_LSPLWP : symmetric migration between LSP and LWP after the bottleneck; m_anc_LSPFU : symmetric migration between LSP and FU before the bottleneck; m_rec_LSPFU : symmetric migration between LSP and FU after the bottleneck; m_anc_LWPFU : symmetric migration between LWP and FU before the bottleneck; m_anc_LWPFU : symmetric migration between LWP and FU after the bottleneck; m_ancPP : symmetric migration before the recent split.
* For migration calculations, to have coherent migration with *Ne*, we used the *Nm* product (effective number of migrants), where $N_m = \frac{(1/F_{st})-1}{4}$, with the $F_{st}$ value drawn from a log-normal distribution ranging from 0.001 to 0.5, and $m = \frac{N_m}{N_{pop}}$.

| Population | LSP | LateLSP | F1 | LWP | CA | TA | GR | VA | VI | FU |
|---|---|---|---|---|---|---|---|---|---|---|
| **LSP** | - | | | | | | | | | |
| **LateLSP** | 0,018 | - | | | | | | | | |
| **F1** | 0,099 | 0,084 | - | | | | | | | |
| **LWP** | 0,259 | 0,259 | 0,076 | - | | | | | | |
| **CA** | 0,278 | 0,265 | 0,093 | 0,050 | - | | | | | |
| **TA** | 0,265 | 0,250 | 0,168 | 0,216 | 0,205 | - | | | | |
| **GR** | 0,213 | 0,202 | 0,076 | 0,079 | 0,057 | 0,131 | - | | | |
| **VA** | 0,188 | 0,177 | 0,088 | 0,133 | 0,135 | 0,136 | 0,077 | - | | |
| **VI** | 0,197 | 0,186 | 0,064 | 0,079 | 0,089 | 0,141 | 0,058 | 0,024 | - | |
| **FU** | 0,197 | 0,186 | 0,060 | 0,072 | 0,082 | 0,141 | 0,055 | 0,028 | 0,003 | - |

Table S5: $F_{ST}$ **values for each pair of Portuguese populations on autosomes.**

| Population | LSP | LWP | FU | ES | RI | OK | TU | WI | BO | PI |
|---|---|---|---|---|---|---|---|---|---|---|
| **LSP** | - | | | | | | | | | |
| **LWP** | 0,0035 | - | | | | | | | | |
| **FU** | 0,0036 | 0,0031 | - | | | | | | | |
| **ES** | 0,0048 | 0,0048 | 0,0047 | - | | | | | | |
| **RI** | 0,0082 | 0,0082 | 0,0082 | 0,0082 | - | | | | | |
| **OK** | 0,0095 | 0,0095 | 0,0095 | 0,0095 | 0,0083 | - | | | | |
| **TU** | 0,0101 | 0,0101 | 0,0100 | 0,0100 | 0,0092 | 0,0081 | - | | | |
| **WI** | 0,0133 | 0,0133 | 0,0133 | 0,0133 | 0,0132 | 0,0131 | 0,0130 | - | | |
| **BO** | 0,0186 | 0,0186 | 0,0186 | 0,0187 | 0,0197 | 0,0201 | 0,0203 | 0,0216 | - | |
| **PI** | 0,0183 | 0,0183 | 0,0183 | 0,0184 | 0,0194 | 0,0198 | 0,0200 | 0,0214 | 0,0111 | - |

Table S6: $D_{XY}$ **values for LSP, LWP, FU and outgroups populations on autosomes.**

| Data subset | Global error rate | local error rate | sd prior error | mean votes concomitent_div | sd votes concomitent_div | mean votes recent_div | sd votes recent_div | mean votes old_div | sd votes old_div | posterior probability | sd posterior probability | selected model |
|---|---|---|---|---|---|---|---|---|---|---|---|---|
| 1 | 0,431 | 0,416 | 0,0014 | 464,5 | 16,55 | 352,6 | 14,99 | 182,9 | 11,42 | 0,584 | 0,01 | concomitant_div |
| 2 | 0,433 | 0,415 | 0,0013 | 377 | 16,59 | 414,7 | 16,61 | 208,3 | 14,75 | 0,585 | 0,02 | recent_div |
| 3 | 0,430 | 0,434 | 0,0011 | 393,8 | 18,54 | 381,7 | 13,16 | 224,5 | 14,67 | 0,566 | 0,03 | concomitant_div |
| 4 | 0,433 | 0,365 | 0,0009 | 403,1 | 12,44 | 410,8 | 12,27 | 186,1 | 14,64 | 0,635 | 0,03 | recent_div |
| 5 | 0,432 | 0,409 | 0,0014 | 422 | 16,83 | 382,1 | 18,53 | 195,9 | 11,18 | 0,591 | 0,01 | concomitant_div |
| 6 | 0,437 | 0,439 | 0,0012 | 355,8 | 10,46 | 437,3 | 13,79 | 206,9 | 13,40 | 0,561 | 0,02 | recent_div |
| 7 | 0,432 | 0,386 | 0,0013 | 380,8 | 11,27 | 391,6 | 9,87 | 227,6 | 11,82 | 0,614 | 0,03 | recent_div |
| 8 | 0,434 | 0,379 | 0,0017 | 368,7 | 12,86 | 428,9 | 15,10 | 202,4 | 11,07 | 0,621 | 0,02 | recent_div |
| 9 | 0,440 | 0,411 | 0,0009 | 353,8 | 18,69 | 455,8 | 15,75 | 190,4 | 10,41 | 0,589 | 0,02 | recent_div |
| 10 | 0,437 | 0,446 | 0,0015 | 378,6 | 15,06 | 390,6 | 11,32 | 230,8 | 12,43 | 0,554 | 0,01 | recent_div |

Table S7: **Results for scenario choice**. The three compared scenarios are detailed in Figure S5. Results are given for the 10 subset of 10,000 simulations for each scenario. Standard deviations over the 10 replicate analyses are given, in addition to the means. The local error rate is $1 - posterior probability$.

| | Posterior point estimates of | | 90% CI | | Global (prior) NMAE computed from | | Local (posterior) NMAE computed | |
|---|---|---|---|---|---|---|---|---|
| Parameter | Mean | Median | Lower bound | Upper bound | Mean | Median | Mean | Median |
| m_anc_LSPFU | -3,39 | -3,32 | -4,82 | -2,10 | 3,48 | 3,40 | 0,21 | 0,21 |
| m_rec_LSPFU | -3,32 | -3,18 | -4,86 | -2,30 | 3,08 | 2,94 | 0,16 | 0,15 |
| m_anc_LSPLWP | -2,56 | -2,66 | -3,95 | -1,03 | 2,93 | 2,88 | 0,51 | 0,52 |
| m_rec_LSPLWP | -3,41 | -3,43 | -4,18 | -2,63 | 1,16 | 1,04 | 0,10 | 0,10 |
| m_anc_LWPFU | -3,33 | -3,19 | -4,96 | -2,04 | 1,16 | 1,06 | 0,18 | 0,17 |
| m_rec_LWPFU | -3,25 | -3,10 | -4,76 | -2,30 | 1,17 | 1,09 | 0,17 | 0,16 |
| Ne_anc_PP | 5,74 | 5,83 | 5,35 | 5,96 | 0,02 | 0,02 | 0,00 | 0,00 |
| Ne_bot_LSP | 1,24 | 1,22 | 0,17 | 2,43 | 36,42 | 30,43 | 2,04 | 1,75 |
| Ne_bot_FU | 2,27 | 2,19 | 0,51 | 4,13 | 2,83 | 2,45 | 0,84 | 0,79 |
| Ne_bot_LWP | 1,47 | 1,43 | 0,23 | 2,88 | 6,68 | 6,07 | 1,92 | 1,80 |
| Ne_LSP | 2,87 | 2,70 | 2,06 | 4,35 | 0,16 | 0,15 | 0,18 | 0,16 |
| Ne_FU | 4,48 | 4,54 | 3,55 | 5,22 | 0,11 | 0,11 | 0,09 | 0,09 |
| Ne_LWP | 3,39 | 3,49 | 2,47 | 3,96 | 0,12 | 0,11 | 0,11 | 0,11 |
| Ne_found_FU | 4,00 | 4,04 | 2,32 | 5,48 | 0,25 | 0,25 | 0,23 | 0,23 |
| Ne_found_LWP | 4,08 | 4,25 | 2,25 | 5,49 | 0,26 | 0,26 | 0,24 | 0,24 |
| Ne_found_LSP | 1,71 | 1,31 | 0,14 | 4,73 | 3,62 | 3,04 | 13,46 | 10,48 |
| Tbot | 1,82 | 1,70 | 1,08 | 2,90 | 0,24 | 0,23 | 0,25 | 0,23 |
| Tsplit_PP | 3,47 | 3,45 | 2,77 | 4,20 | 0,08 | 0,07 | 0,09 | 0,09 |

Table S8: **Results for estimation of parameters of interest for the scenario *concomitent_div*.** Mean and median estimates for each parameter were calculated using an ABCRF run, conducted with a reference table containing 158 summary statistics from 100,000 simulations. Quantile at 5 (lower bound) and 95% (upper bound) was used to have the 90% condifdence interval (CI). We inferred computed global and local accuracy metrics corresponding to global and local NMAE (normalized mean absolute error which is the average absolute difference between the point estimate and the true simulated value divided by the true simulated value) with the mean and the median as points estimates.

| Genes | Function | Chromosome | Start Position (in bp) | End Position (in bp) | max. Lscore ($-\log_{10} \mathbf{P}_{C2}$) |
|---|---|---|---|---|---|
| l(2)37Cc | protein l(2)37Cc | 1 | 7086688 | 7092154 | 6.26 (5.19) |
| shu | inactive peptidyl-prolyl cis-trans isomerase shutdown-like | 1 | 7094871 | 7098433 | 6.26 (5.19) |
| MYO7a | myosin-VIIa-like isoform X1 | 1 | 7101279 | 7119323 | 6.26 (5.19) |
| DNAH7 | dynein axonemal heavy chain 7 | 1 | 20256316 | 20315622 | 6.14 (7.15) |
| tweek | transmembrane protein KIAA1109 | 1 | 24601825 | 24680002 | 8.04 (5.17) |
| Tret1 | facilitated trehalose transporter Tret1-2 homolog isoform X2 | 3 | 15481926 | 15557587 | 7.9 (6.06) |
| peritrophin-A | peritrophin-1-like isoform X2 | 4 | 2262121 | 2265100 | 8.14 (6.1) |
| ASTER-B/LAM4 | protein Aster-B-like isoform X2/<br>membrane-anchored lipid-binding protein LAM4-like isoform X4 | 4 | 8674178 | 8732605 | 7.71 (4.76) |
| MYO9A | unconventional myosin-IXa-like isoform X4 | 4 | 11762364 | 11820758 | 6.43 (6.9) |
| ZFP431 | zinc finger protein 431-like isoform X2 | 5 | 4910610 | 5236798 | 5.62 (5.68) |
| FASN1 | fatty acid synthase-like isoform X1 | 5 | 5017837 | 5040104 | 5.62 (5.68) |
| eIF4H1 | eukaryotic translation initiation factor 4H isoform X3 | 5 | 12739271 | 12755087 | 5.53 (5.91) |
| ABDH11 | protein ABHD11-like | 5 | 12943125 | 13000397 | 6.23 (8.23) |
| CCNB1IP1 | ubiquitin-protein ligase CCNB1IP1-like | 6 | 6676751 | 6677515 | 5.51 (5.13) |
| AmFPI-1 | fungal protease inhibitor-1-like isoform X2 | 8 | 11815953 | 11827125 | 5.9 (5) |
| AstC-R1 | allatostatin receptor type C | 9 | 12972534 | 12973781 | 5.36 (4.44) |
| MON2 | protein MON2 homolog isoform X3 | 10 | 10619648 | 10673875 | 7.07 (5.21) |
| SNMP2 | sensory neuron membrane protein 2 | 11 | 10639483 | 10671897 | 7.62 (4.86) |
| nclb | periodic tryptophan protein 2 homolog | 11 | 10673773 | 10696420 | 7.62 (4.86) |
| emb | exportin-1 | 13 | 2213566 | 2221112 | 7.19 (4.81) |
| Clptm1 | cleft lip and palate transmembrane protein 1 homolog | 13 | 2229188 | 2233065 | 7.19 (4.81) |
| mGluR | metabotropic glutamate receptor 2 | 14 | 724899 | 733156 | 7.87 (7.49) |
| Ork1 | potassium channel subfamily K member 18 | 14 | 5045071 | 5280319 | 6.76 (5.46) |
| WDR48 | WD repeat-containing protein 48 homolog | 15 | 8893309 | 8898302 | 5.76 (7.76) |
| ZSCAN21/ZNF219 | zinc finger and SCAN domain-containing protein 21 | 16 | 4803695 | 4939251 | 7.51 (4.86) |
| BmLHA | lipase member H-A isoform X1 | 16 | 8527212 | 8547261 | 11.35 (5.21) |
| CLIPB9 | CLIP domain-containing serine protease B9-like | 17 | 5776296 | 5800136 | 5.37 (5.03) |
| PC | phosphatidylcholine:ceramide cholinephosphotransferase 2 isoform X1 | 17 | 9193406 | 9358062 | 5.53 (4.14) |
| Npl4 | nuclear protein localization protein 4 homolog isoform X1 | 20 | 149419 | 174659 | 5.51 (4.19) |
| SLC23A2 | solute carrier family 23 member 2 isoform X1 | 20 | 1258692 | 1300590 | 5.81 (5.2) |
| ckn | caskin-2 isoform X1 | 20 | 5431521 | 5431868 | 5.9 (4.37) |
| DYRK2 | probable serine/threonine-protein kinase dyrk2 | 21 | 1136699 | 1328116 | 5.94 (6.96) |
| PCFT | proton-coupled folate transporter-like | 21 | 4844736 | 4864940 | 5.64 (4.07) |
| SLC | sialin isoform X1 | 21 | 6267638 | 6330502 | 5.74 (7.74) |
| bai | transmembrane emp24 domain-containing protein bai isoform X1 | 22 | 3429817 | 3440708 | 5.91 (5.9) |
| Rox8 | nucleolysin TIAR | 22 | 3462491 | 3476949 | 5.91 (5.9) |
| Dsx | protein doublesex isoform X1 | 22 | 5656778 | 5845445 | 5.56 (4.26) |
| Sn | protein singed | 23 | 530661 | 569312 | 6.68 (6.39) |
| MANF | mesencephalic astrocyte-derived neurotrophic factor homolog | 23 | 1750925 | 1763373 | 5.73 (5.98) |
| ATP synthase subunit f | putative ATP synthase subunit f, mitochondrial | 23 | 3777992 | 3779324 | 5.73 (5.98) |
| RNaseZ | zinc phosphodiesterase ELAC protein 2 isoform X1 | 23 | 3781220 | 3827690 | 5.73 (5.98) |
| koi | klaroid protein | 23 | 4926183 | 4980140 | 6.96 (5.85) |
| Dscam2 | Down syndrome cell adhesion molecule-like protein Dscam2 | 24 | 600758 | 666934 | 5.53 (4.5) |
| CCDC47 | PAT complex subunit CCDC47 | 26 | 6077465 | 6086495 | 7.24 (5.07) |
| ilys-3 | invertebrate-type lysozyme 3-like | 26 | 6093712 | 6112332 | 7.24 (5.07) |
| KAT6B | histone acetyltransferase KAT6B isoform X3 | 26 | 9195478 | 9229839 | 5.93 (7.93) |
| Jerky | Jerky protein | 27 | 5853355 | 5853732 | 5.66 (5.24) |
| ACE | angiotensin-converting enzyme | 27 | 8483406 | 8485526 | 5.23 (6.07) |
| dri | protein dead ringer isoform X1 | 28 | 3817042 | 4072213 | 5.31 (4.81) |
| FH | fumarate hydratase, mitochondrial-like isoform X1 | 31 | 8838573 | 8840219 | 5.66 (5.02) |
| AQP | aquaporin AQPAn.G | 31 | 8848141 | 8850924 | 5.66 (5.02) |
| Mth2 | G-protein coupled receptor Mth2 isoform X4 | 35 | 2617293 | 2668356 | 10.01 (5.79) |
| PHC3 | polyhomeotic-like protein 3 isoform X3 | 35 | 3825415 | 4083633 | 8.81 (5.04) |
| hid | protein HID1 | 35 | 7801692 | 7831733 | 6.69 (5.59) |
| hRED1 | double-stranded RNA-specific editase 1-like isoform X3 | 35 | 9433836 | 9449512 | 6.35 (5.6) |
| CGRP1 | calcitonin gene-related peptide type 1 receptor isoform X2 | 35 | 9736346 | 9753981 | 9.82 (5.36) |
| trpm | transient receptor potential cation channel trpm isoform X18 | 36 | 786465 | 807130 | 5.47 (7.47) |
| Stard13 | stAR-related lipid transfer protein 13 isoform X1 | 37 | 4275905 | 4692751 | 5.46 (5.35) |
| klar | klarsicht protein isoform X3 | 37 | 6193212 | 6513041 | 5.63 (5.25) |
| FucTA | glycoprotein 3-alpha-L-fucosyltransferase A-like | 37 | 8712093 | 8720394 | 7.2 (4.56) |
| IDE | insulin-degrading enzyme | 38 | 7898268 | 7933296 | 8.07 (5.69) |
| brat | brain tumor protein isoform X1 | 39 | 5182473 | 5302948 | 5.84 (5.1) |
| LRR24 | leucine-rich repeat-containing protein 24-like | 39 | 6405432 | 6407186 | 5.64 (5.19) |
| Neurobeachin | neurobeachin isoform X5 | 39 | 7746643 | 8489304 | 10.09 (7.3) |
| mRRF2 | ribosome-releasing factor 2, mitochondrial-like | 40 | 7488163 | 7517536 | 8.09 (7.4) |
| AMPdeam | AMP deaminase 2 isoform X3 | 40 | 7519151 | 7538957 | 8.09 (7.4) |
| bnl | fibroblast growth factor 3 isoform X1 | 41 | 3000079 | 3262487 | 7.49 (4.07) |
| TRH-DE | thyrotropin-releasing hormone-degrading ectoenzyme isoform X2 | 42 | 4800183 | 4830409 | 7.34 (5.19) |
| betaTub60D | tubulin beta chain | 43 | 6191342 | 6193643 | 6.29 (7.49) |
| PrBP | cGMP-specific 3',5'-cyclic phosphodiesterase isoform X2 | 43 | 7144790 | 7192884 | 6.26 (5.73) |

*Continued on next page...*

| Genes | Function | Chromosome | Start Position (in bp) | End Position (in bp) | max. Lscore ($-\log_{10} \mathbf{P}_{C2}$) |
|---|---|---|---|---|---|
| Dop2R | alpha-2Db adrenergic receptor | 46 | 1554494 | 1976066 | 6.25(5.33) |
| Mblk-1 | mushroom body large-type Kenyon cell-specific protein 1 isoform X1 | 46 | 3648230 | 3847118 | 5.27 (5.95) |
| polybromo | protein polybromo-1 isoform X12 | 46 | 6245055 | 6284964 | 5.21 (4.25) |
| C/EBPZ | CCAAT/enhancer-binding protein zeta | 49 | 2929107 | 2960526 | 5.14 (5.24) |
| Arp2 | actin-related protein 2 | 49 | 2961902 | 2977048 | 5.14 (5.24) |
| LRDD1 | leucine-rich repeat and death domain-containing protein 1 | Z | 5881689 | 5913766 | 9.79 (4.47) |
| NPAS2 | neuronal PAS domain-containing protein 2-like isoform X2 | Z | 6823186 | 6951715 | 11.29 (4.72) |
| $GABA_B$ | gamma-aminobutyric acid receptor subunit beta-like isoform X1 | Z | 8383068 | 8436899 | 13.43 (6.53) |
| Nwk | protein nervous wreck isoform X3 | Z | 12063177 | 12158967 | 15.31 (4.65) |
| Coq8 | atypical kinase COQ8B, mitochondrial isoform X1 | Z | 12173033 | 12182476 | 27.75 (4.93) |
| hydr2 | phosphatidylserine lipase ABHD16A | Z | 12195371 | 12206944 | 27.75 (4.93) |
| GPR107 | protein GPR107 | Z | 12211842 | 12231778 | 27.75 (4.93) |
| wcy | WW domain-containing adapter protein with coiled-coil homolog isoform X2 | Z | 14088679 | 14125327 | 26.15 (4.62) |
| Dikar | uncharacterized protein Dikar isoform X3 | Z | 14128253 | 14155383 | 26.15 (4.62) |
| Pfdn1 | prefoldin subunit 1 | Z | 14157876 | 14158262 | 26.15 (4.62) |
| RalGPS2 | ras-specific guanine nucleotide-releasing factor RalGPS2 isoform X3 | Z | 14173553 | 14213051 | 26.15 (4.62) |
| NEURL4 | neuralized-like protein 4 | Z | 14329649 | 14371365 | 12.58 (5.94) |
| NEP2 | neprilysin-2 isoform X3 | Z | 14376632 | 14445049 | 53.53 (5.85) |
| Cat | catalase domain-containing protein | Z | 14839535 | 14841911 | 22.33 (4.48) |
| Spn-E | Probable ATP-dependent RNA helicase spindle-E | Z | 15100596 | 15100898 | 23.88 (4.99) |
| Zfp260/ZNF260 | zinc finger protein 260-like isoform X1 | Z | 15156568 | 15177529 | 24.14 (4.03) |
| SP2353 | basement membrane-specific heparan sulfate proteoglycan core protein-like isoform X5 | Z | 21755125 | 22034490 | 16.92 (3.5) |
| Karln | kalirin isoform X2 | Z | 22998621 | 23258778 | 16.18 (4.1) |

Table S9: **Genes identified in significant windows detected by Baypass.** A total of 93 genes were identified, with 75 on autosomes and 18 on the Z chromosome. Genes annotated as 'uncharacterized' or 'unnamed' are not included in this report. The table gives for each gene i) its position in the genome; ii) its start and end position in pb iii); and the maximum Lindley score (Lscore) and the most significant C2 Pvalues (in $-\log_{10}$ scale) of the significant window detected by Baypass for the gene.

| Gene number | Chromosome | Start Position (in bp) | End Position (in bp) | Function |
|---|---|---|---|---|
| 1 | chrZ | 11987971 | 12043052 | PHD finger protein rhinoceros |
| 2 | chrZ | 12063177 | 12158967 | protein nervous wreck isoform X3 |
| 3 | chrZ | 12173033 | 12182476 | atypical kinase COQ8B, mitochondrial isoform X1 |
| 4 | chrZ | 12195371 | 12206944 | phosphatidylserine lipase ABHD16A |
| 5 | chrZ | 12211842 | 12231778 | protein GPR107 |
| 6 | chrZ | 12244534 | 12269171 | ras-related and estrogen-regulated growth inhibitor |
| 7 | chrZ | 12250229 | 12250768 | - |
| 8 | chrZ | 12284245 | 12294275 | nuclear receptor subfamily 2 group E member 1 |
| 9 | chrZ | 12316396 | 12375653 | calcium-binding mitochondrial carrier protein SCaMC-2 isoform X1 |
| 10 | chrZ | 12391057 | 12405774 | ATP-binding cassette sub-family F member 2 |
| 11 | chrZ | 12413253 | 12423645 | ADP,ATP carrier protein-like |
| 12 | chrZ | 12430332 | 12431312 | - |
| 13 | chrZ | 12436662 | 12437236 | unnamed protein product |
| 14 | chrZ | 12437508 | 12448069 | LETM1 domain-containing protein 1 |
| 15 | chrZ | 12470451 | 12470984 | zinc finger BED domain-containing protein 5-like |
| 16 | chrZ | 12495275 | 12501841 | LETM1 domain-containing protein 1 isoform X2 |
| 17 | chrZ | 12518212 | 12521445 | uncharacterized protein LOC113508497 |
| 18 | chrZ | 12524303 | 12530521 | solute carrier family 66 member 3 |
| 19 | chrZ | 12579998 | 12728836 | CUB and sushi domain-containing protein 2 |
| 20 | chrZ | 12748253 | 12833907 | protein phosphatase 1 regulatory subunit 12B isoform X15 |
| 21 | chrZ | 12840043 | 12947155 | disco-interacting protein 2 isoform X3 |
| 22 | chrZ | 12947184 | 12952002 | disco-interacting protein 2 isoform X2 |
| 23 | chrZ | 12982358 | 12990755 | - |
| 24 | chrZ | 13028944 | 13073596 | arf-GAP with Rho-GAP domain, ANK repeat and PH domain-containing protein 2 isoform X1 |
| 25 | chrZ | 13077744 | 13087647 | EF-hand calcium-binding domain-containing protein 1 isoform X1 |
| 26 | chrZ | 13107244 | 13116233 | calcitonin gene-related peptide type 1 receptor-like isoform X3 |
| 27 | chrZ | 13125979 | 13132874 | sodium/potassium/calcium exchanger 3 |
| 28 | chrZ | 13154399 | 13169573 | sodium channel protein Nach isoform X1 |
| 29 | chrZ | 13177561 | 13190742 | peptidoglycan-recognition protein LC-like isoform X1 |
| 30 | chrZ | 13203575 | 13215814 | LITAF domain-containing protein-like |
| 31 | chrZ | 13217721 | 13222705 | beta-parvin |
| 32 | chrZ | 13238586 | 13275797 | - |
| 33 | chrZ | 13336071 | 13392293 | leucine-rich repeat-containing protein 15-like |
| 34 | chrZ | 13390605 | 13392293 | leucine-rich repeat-containing protein 15-like |
| 35 | chrZ | 13418745 | 13419188 | NADH-ubiquinone oxidoreductase subunit 8 |
| 36 | chrZ | 13586620 | 13626650 | PEHE domain-containing protein |
| 37 | chrZ | 13639824 | 13650283 | voltage-gated potassium channel subunit beta-2 isoform X7 |
| 38 | chrZ | 13667785 | 13686880 | sodium-coupled monocarboxylate transporter 2-like |
| 39 | chrZ | 13691643 | 13706591 | sodium/potassium/calcium exchanger 3 isoform X6 |
| 40 | chrZ | 13713472 | 13727880 | sodium/potassium/calcium exchanger 3-like |
| 41 | chrZ | 13731927 | 13738546 | - |
| 42 | chrZ | 13748190 | 13759422 | MATH and LRR domain-containing protein PFE0570w-like |
| 43 | chrZ | 13768649 | 13782629 | ER degradation-enhancing alpha-mannosidase-like protein 3 |
| 44 | chrZ | 13787641 | 13799064 | - |
| 45 | chrZ | 13808047 | 13831311 | - |
| 46 | chrZ | 13835369 | 13840467 | General receptor for phosphoinositides 1-associated scaffold protein |
| 47 | chrZ | 13846524 | 13858058 | collagen alpha-1(I) chain-like |
| 48 | chrZ | 13866511 | 13879875 | ras-GEF domain-containing family member 1B |
| 49 | chrZ | 13904081 | 13928683 | dynein heavy chain 6, axonemal |
| 50 | chrZ | 14013897 | 14043787 | uncharacterized oxidoreductase YtbE isoform X1 |
| 51 | chrZ | 14051198 | 14077312 | dynein axonemal heavy chain 6 |
| 52 | chrZ | 14088679 | 14125327 | WW domain-containing adapter protein with coiled-coil homolog isoform X2 |
| 53 | chrZ | 14128253 | 14155383 | hypothetical protein B5X24_HaOG215492 |
| 54 | chrZ | 14157876 | 14158262 | prefoldin subunit 1 |
| 55 | chrZ | 14173553 | 14213051 | ras-specific guanine nucleotide-releasing factor RalGPS2 isoform X3 |
| 56 | chrZ | 14223548 | 14225742 | - |
| 57 | chrZ | 14264408 | 14303098 | protein cab-1 isoform X2 |
| 58 | chrZ | 14329649 | 14371365 | neuralized-like protein 4 |
| 59 | chrZ | 14376632 | 14445049 | neprilysin-2 isoform X3 |
| 60 | chrZ | 14581829 | 14604332 | amyloid beta A4 precursor protein-binding family B member 1-interacting protein isoform X2 |
| 61 | chrZ | 16613707 | 16618186 | surfeit locus protein 1 |
| 62 | chrZ | 16633941 | 16677507 | ankyrin repeat domain-containing protein 11 isoform X2 |
| 63 | chrZ | 16687474 | 16700769 | prostaglandin E synthase 2 |
| 64 | chrZ | 16729550 | 16856282 | dendritic arbor reduction protein 1-like isoform X2 |
| 65 | chrZ | 16863869 | 16896005 | multidrug resistance protein homolog 49-like |
| 66 | chrZ | 16899831 | 16927769 | multidrug resistance protein homolog 49-like isoform X1 |
| 67 | chrZ | 16936019 | 16961825 | circadian locomoter output cycles protein kaput isoform X5 |
| 68 | chrZ | 16976454 | 17010019 | - |
| 69 | chrZ | 17020977 | 17021477 | - |

*Continued on next page...*

| Gene number | Chromosome | Start Position (in bp) | End Position (in bp) | Function |
|---|---|---|---|---|
| 70 | chrZ | 17024265 | 17029564 | palmitoyltransferase ZDHHC16 |
| 71 | chrZ | 17033848 | 17045209 | T-cell acute lymphocytic leukemia protein 1 homolog |
| 72 | chrZ | 17059983 | 17103359 | peptide transporter family 1 isoform X1 |
| 73 | chrZ | 17105678 | 17105890 | - |
| 74 | chrZ | 17123077 | 17128673 | BPTI/Kunitz domain-containing protein |
| 75 | chrZ | 17132826 | 17217781 | SPARC-related modular calcium-binding protein 1 isoform X4 |
| 76 | chrZ | 17201176 | 17202323 | - |
| 77 | chrZ | 17256295 | 17283880 | ubiquitin carboxyl-terminal hydrolase 35-like isoform X2 |
| 78 | chrZ | 17292335 | 17299073 | - |
| 79 | chrZ | 17303197 | 17304030 | ubiquinol-cytochrome c reductase iron-sulfur subunit-like |
| 80 | chrZ | 17305098 | 17320734 | - |
| 81 | chrZ | 17333282 | 17348101 | polypeptide N-acetylgalactosaminyltransferase 35A-like isoform X2 |
| 82 | chrZ | 17380362 | 17496295 | discoidin domain-containing receptor 2 isoform X2 |
| 83 | chrZ | 17395812 | 17396822 | - |
| 84 | chrZ | 17714780 | 17715841 | lipase 3-like |
| 85 | chrZ | 17724181 | 17725569 | - |
| 86 | chrZ | 17769200 | 17780533 | - |
| 87 | chrZ | 17784747 | 17818900 | V-type proton ATPase 116 kDa subunit a 1-like isoform X1 |
| 88 | chrZ | 17841253 | 17864057 | - |
| 89 | chrZ | 17882159 | 17886635 | - |
| 90 | chrZ | 17890894 | 17892370 | - |
| 91 | chrZ | 17894621 | 17896670 | - |
| 92 | chrZ | 17897358 | 17900133 | - |
| 93 | chrZ | 17929504 | 17929716 | protein grindelwald |
| 94 | chrZ | 17953375 | 17989958 | period circadian protein isoform X29 |
| 95 | chrZ | 18025529 | 18234179 | hypothetical protein SFRURICE_007727 |
| 96 | chrZ | 18251699 | 18267049 | RING finger and SPRY domain-containing protein 1-like |
| 97 | chrZ | 18286057 | 18286383 | - |
| 98 | chrZ | 18414106 | 18456948 | flotillin-1 isoform X1 |
| 99 | chrZ | 18461197 | 18480632 | sulfhydryl oxidase 1 isoform X3 |
| 100 | chrZ | 18495927 | 18523956 | transcription factor 25-like isoform X6 |
| 101 | chrZ | 18606436 | 18676640 | protein scabrous |
| 102 | chrZ | 18710159 | 18734424 | intraflagellar transport protein 122 homolog |
| 103 | chrZ | 18740887 | 18741897 | fork head domain-containing protein crocodile-like |

Table S10: **Genes identified in the zoomed regions from 12,000,000 to 14,600,000 bp and from 16,600,600 to 18,800,000 bp**. The gene number corresponds to the numbering in Figures 6A and 6B. Genes that are not annotated in the gff file are indicated with a "-".

| Chromosome | Position | Gene | Transcript ID | Mutation type MIS | REF/ALT | Well polarized | Ancestral | Derived | AA Ancestral ou Ref | AA Derived ou Alt | Frequency derived or alt LSP | Frequency derived or alt LWP |
|---|---|---|---|---|---|---|---|---|---|---|---|---|
| chrZ | 12177551 | Coq8 | atypical kinase COQ8B, mitochondrial | DELETERIOUS | C/G | Yes | C | G | Glu | Gln | 0,00 | 0,13 |
| chrZ | 12178791 | Coq8 | atypical kinase COQ8B, mitochondrial | DELETERIOUS | A/G | Yes | A | G | Val | Ala | 0,00 | 1,00 |
| chrZ | 12180901 | Coq8 | atypical kinase COQ8B, mitochondrial | DELETERIOUS | G/A | Yes | G | A | Leu | Phe | 0,00 | 0,04 |
| chrZ | 12181298 | Coq8 | atypical kinase COQ8B, mitochondrial | DELETERIOUS | G/A | No | ? | ? | Thr | Ile | 0,00 | 0,00 |
| chrZ | 12182215 | Coq8 | atypical kinase COQ8B, mitochondrial | DELETERIOUS | C/T | Yes | C | T | Asp | Asn | 0,00 | 0,00 |
| chrZ | 14329788 | NEURL4 | neuralized-like protein 4 | TOLERANT | A/T | No | ? | ? | Tyr | Phe | 0,00 | 0,00 |
| chrZ | 14334752 | NEURL4 | neuralized-like protein 4 | TOLERANT | T/G | Yes | T | G | Ile | Met | 0,00 | 0,00 |
| chrZ | 14334819 | NEURL4 | neuralized-like protein 4 | TOLERANT | C/T | Yes | C | T | Pro | Ser | 0,03 | 0,00 |
| chrZ | 14341299 | NEURL4 | neuralized-like protein 4 | DELETERIOUS | C/T | No | ? | ? | Thr | Met | 0,00 | 0,00 |
| chrZ | 14357887 | NEURL4 | neuralized-like protein 4 | DELETERIOUS | G/A | Y | G | A | Asp | Asn | 0,00 | 0,00 |
| chrZ | 14361997 | NEURL4 | neuralized-like protein 4 | DELETERIOUS | G/A | No | ? | ? | Glu | Lys | 0,13 | 0,50 |
| chrZ | 14362006 | NEURL4 | neuralized-like protein 4 | TOLERANT | G/A | No | ? | ? | Ala | Thr | 0,13 | 0,50 |
| chrZ | 14362007 | NEURL4 | neuralized-like protein 4 | DELETERIOUS | C/T | No | ? | ? | Ala | Val | 0,17 | 0,50 |
| chrZ | 14362033 | NEURL4 | neuralized-like protein 4 | TOLERANT | A/T | Yes | A | T | Met | Leu | 0,00 | 0,46 |
| chrZ | 14363683 | NEURL4 | neuralized-like protein 4 | DELETERIOUS | A/G | No | ? | ? | Asp | Gly | 0,20 | 0,58 |
| chrZ | 14363694 | NEURL4 | neuralized-like protein 4 | TOLERANT | A/G | No | ? | ? | Met | Val | 0,20 | 0,54 |
| chrZ | 14363707 | NEURL4 | neuralized-like protein 4 | DELETERIOUS | G/A | No | ? | ? | Gly | Glu | 0,20 | 0,54 |
| chrZ | 14363713 | NEURL4 | neuralized-like protein 4 | DELETERIOUS | A/T | No | ? | ? | Asn | Ile | 0,17 | 0,54 |
| chrZ | 14363717 | NEURL4 | neuralized-like protein 4 | TOLERANT | A/C | No | ? | ? | Glu | Asp | 0,17 | 0,50 |
| chrZ | 14363719 | NEURL4 | neuralized-like protein 4 | DELETERIOUS | C/T | No | ? | ? | Ala | Val | 0,17 | 0,50 |
| chrZ | 14363722 | NEURL4 | neuralized-like protein 4 | DELETERIOUS | T/C | No | ? | ? | Val | Ala | 0,20 | 0,50 |
| chrZ | 14363727 | NEURL4 | neuralized-like protein 4 | TOLERANT | A/G | No | ? | ? | Ile | Val | 0,20 | 0,50 |
| chrZ | 14365818 | NEURL4 | neuralized-like protein 4 | DELETERIOUS | T/A | No | ? | ? | Met | Lys | 0,20 | 0,50 |
| chrZ | 14365850 | NEURL4 | neuralized-like protein 4 | DELETERIOUS | G/A | No | ? | ? | Glu | Lys | 0,20 | 0,50 |
| chrZ | 14367257 | NEURL4 | neuralized-like protein 4 | TOLERANT | T/G | No | ? | ? | Leu | Val | 0,20 | 0,50 |
| chrZ | 14367270 | NEURL4 | neuralized-like protein 4 | DELETERIOUS | T/C | No | ? | ? | Met | Thr | 0,20 | 0,50 |
| chrZ | 14367366 | NEURL4 | neuralized-like protein 4 | DELETERIOUS | G/C | No | ? | ? | Arg | Pro | 0,00 | 0,00 |
| chrZ | 14382659 | NEP-2 | neprilysin-2 isoform | TOLERANT | C/T | No | ? | ? | Arg | His | 0,00 | 0,00 |
| chrZ | 14386593 | NEP-2 | neprilysin-2 isoform | DELETERIOUS | G/A | No | ? | ? | Ala | Val | 0,00 | 0,00 |
| chrZ | 16942563 | Clk | circadian locomoter output cycles protein kaput | TOLERANT | C/T | Yes | C | T | Gly | Ser | 0,00 | 0,13 |
| chrZ | 17953457 | period | period circadian protein | TOLERANT | C/T | Yes | C | T | Arg | His | 0,00 | 0,00 |
| chrZ | 17958550 | period | period circadian protein | DELETERIOUS | G/A | Yes | G | A | Thr | Ile | 0,17 | 0,42 |
| chrZ | 17960217 | period | period circadian protein | TOLERANT | T/C | Yes | T | C | Ile | Val | 0,00 | 0,00 |
| chrZ | 17966180 | period | period circadian protein | TOLERANT | A/T | Yes | A | T | Leu | Met | 0,17 | 0,00 |
| chrZ | 17967070 | period | period circadian protein | DELETERIOUS | C/T | Yes | C | T | Cys | Tyr | 0,83 | 1,00 |
| chrZ | 17967604 | period | period circadian protein | TOLERANT | A/G | Yes | A | G | Ser | Pro | 0,00 | 0,00 |
| chrZ | 17969351 | period | period circadian protein | DELETERIOUS | G/A | Yes | G | A | Ser | Phe | 0,00 | 0,50 |
| chrZ | 17974548 | period | period circadian protein | TOLERANT | C/T | Yes | C | T | Ser | Asn | 0,00 | 0,04 |
| chrZ | 17978608 | period | period circadian protein | DELETERIOUS | A/T | No | ? | ? | Val | Asp | 0,00 | 0,00 |
| chrZ | 17980250 | period | period circadian protein | DELETERIOUS | A/G | Yes | A | G | Val | Ala | 0,80 | 0,00 |
| chrZ | 17982524 | period | period circadian protein | TOLERANT | T/A | No | ? | ? | Glu | Asp | 0,00 | 0,00 |
| chrZ | 17982553 | period | period circadian protein | TOLERANT | T/C | Yes | T | C | Ile | Val | 0,00 | 0,04 |

Table S11: All missense (MIS) mutations identified in the *Coq8*, *NEURL4*, *NEP-2*, *Clk*, and *period* genes. For each site, the table reports the position on the Z chromosome, the mutation type, the reference and alternative alleles (REF/ALT), the polarization status, the ancestral and derived states (or “?” if unresolved), the corresponding amino acids (ancestral/derived or REF/ALT if not polarized), and the frequency of the derived (or alternative) allele estimated on ind-seq data for LSP and LWP.

| Population | LSP | LateLSP | F1 | LWP | CA | TA | GR | VA | VI | FU |
|---|---|---|---|---|---|---|---|---|---|---|
| **LSP** | - | | | | | | | | | |
| **LateLSP** | 0,018 | - | | | | | | | | |
| **F1** | 0,177 | 0,144 | - | | | | | | | |
| **LWP** | 0,449 | 0,420 | 0,122 | - | | | | | | |
| **CA** | 0,444 | 0,408 | 0,126 | 0,047 | - | | | | | |
| **TA** | 0,396 | 0,365 | 0,189 | 0,237 | 0,223 | - | | | | |
| **GR** | 0,372 | 0,339 | 0,101 | 0,077 | 0,053 | 0,147 | - | | | |
| **VA** | 0,340 | 0,309 | 0,114 | 0,154 | 0,149 | 0,147 | 0,092 | - | | |
| **VI** | 0,353 | 0,321 | 0,095 | 0,100 | 0,104 | 0,146 | 0,067 | 0,020 | - | |
| **FU** | 0,350 | 0,318 | 0,091 | 0,085 | 0,090 | 0,149 | 0,059 | 0,027 | 0,004 | - |

Table S12: $F_{ST}$ **values for each pair of Portuguese populations on the Z chromosome.**

| Population | LSP | LWP | FU | ES | RI | OK | TU | WI | BO | PI |
|---|---|---|---|---|---|---|---|---|---|---|
| **LSP** | - | | | | | | | | | |
| **LWP** | 0,0019 | - | | | | | | | | |
| **FU** | 0,0020 | 0,0017 | - | | | | | | | |
| **ES** | 0,0030 | 0,0030 | 0,0030 | - | | | | | | |
| **RI** | 0,0063 | 0,0063 | 0,0063 | 0,0063 | - | | | | | |
| **OK** | 0,0079 | 0,0079 | 0,0079 | 0,0079 | 0,0067 | - | | | | |
| **TU** | 0,0083 | 0,0083 | 0,0083 | 0,0083 | 0,0075 | 0,0059 | - | | | |
| **WI** | 0,0118 | 0,0118 | 0,0118 | 0,0118 | 0,0117 | 0,0117 | 0,0115 | - | | |
| **BO** | 0,0158 | 0,0158 | 0,0158 | 0,0159 | 0,0166 | 0,0171 | 0,0173 | 0,0183 | - | |
| **PI** | 0,0157 | 0,0157 | 0,0156 | 0,0158 | 0,0164 | 0,0169 | 0,0171 | 0,0182 | 0,0082 | - |

Table S13: $D_{XY}$ **values for each pair of Portuguese populations on the Z chromosome.**

### 2 Supplementary Figures

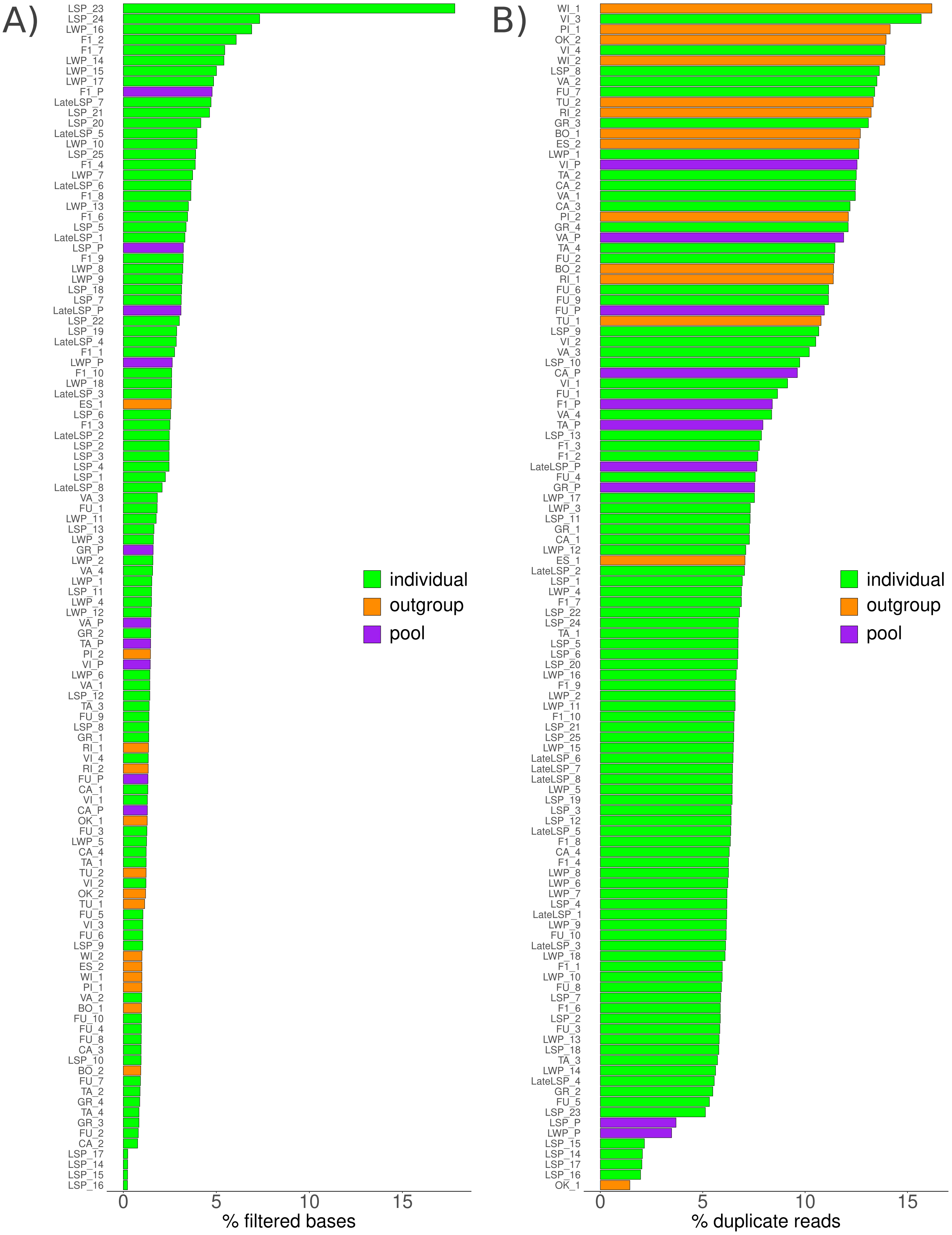


Figure S1: **(A)** The percentage of filtered-out reads for each Portuguese individual, pool and outgroup. **(B)** The percentage of duplicated reads for each individual, pool and outgroup. All Portuguese individuals, pools and outgroups were paired-end (150 bp) sequenced using a NovaSeq6000, with the exception of 4 individuals sequenced on an Illumina HiSeq2500 (LSP_14, LSP_15, LSP_16 and LSP_17).

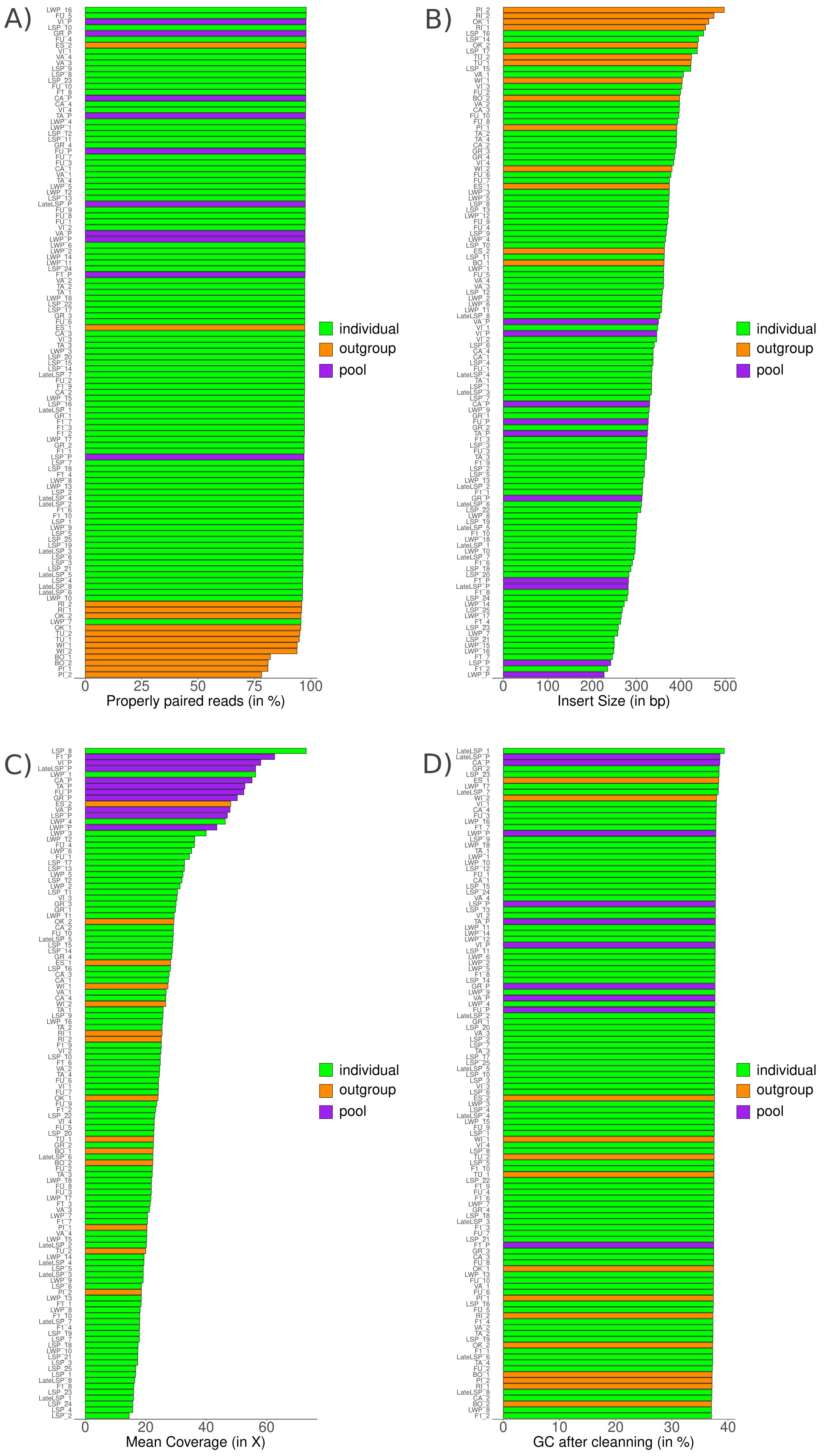


Figure S2: **(A)** The percentage of reads mapped and properly paired, **(B)** the mean insert size, **(C)** the mean depth and **(D)** the percentage of GC content for each Portuguese individual, pool and outgroup.

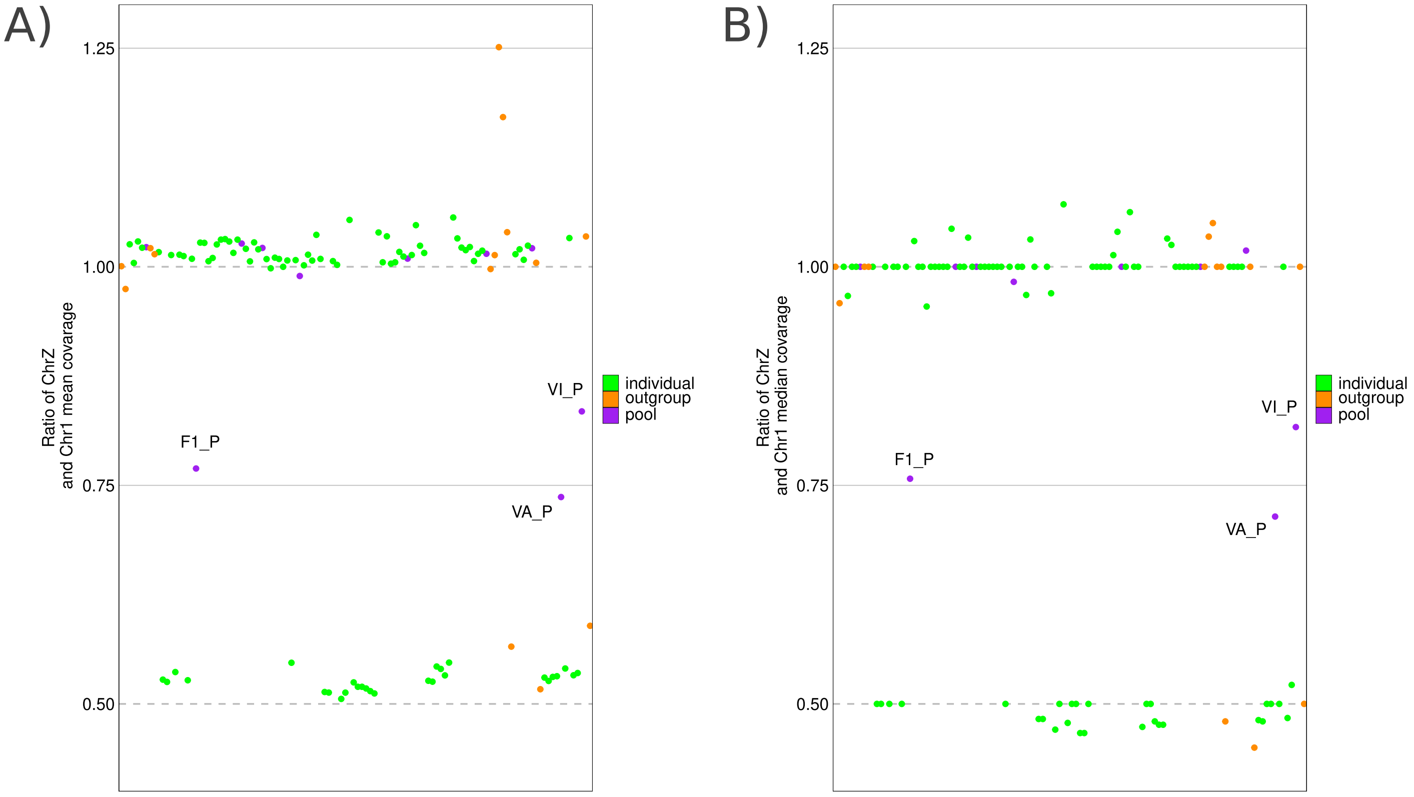


Figure S3: **(A)** Ratio of the mean coverage of chromosome Z to the mean coverage of chromosome 1. The line at 0.5 represents the expected value for identifying females, as heterogametic females exhibit half the coverage on chromosome Z compared to chromosome 1. The line at 1 corresponds to the expected value for males, where coverage is equal between chromosome Z and chromosome 1. **(B)** Ratio of the median coverage of chromosome Z to the mean coverage of chromosome 1.

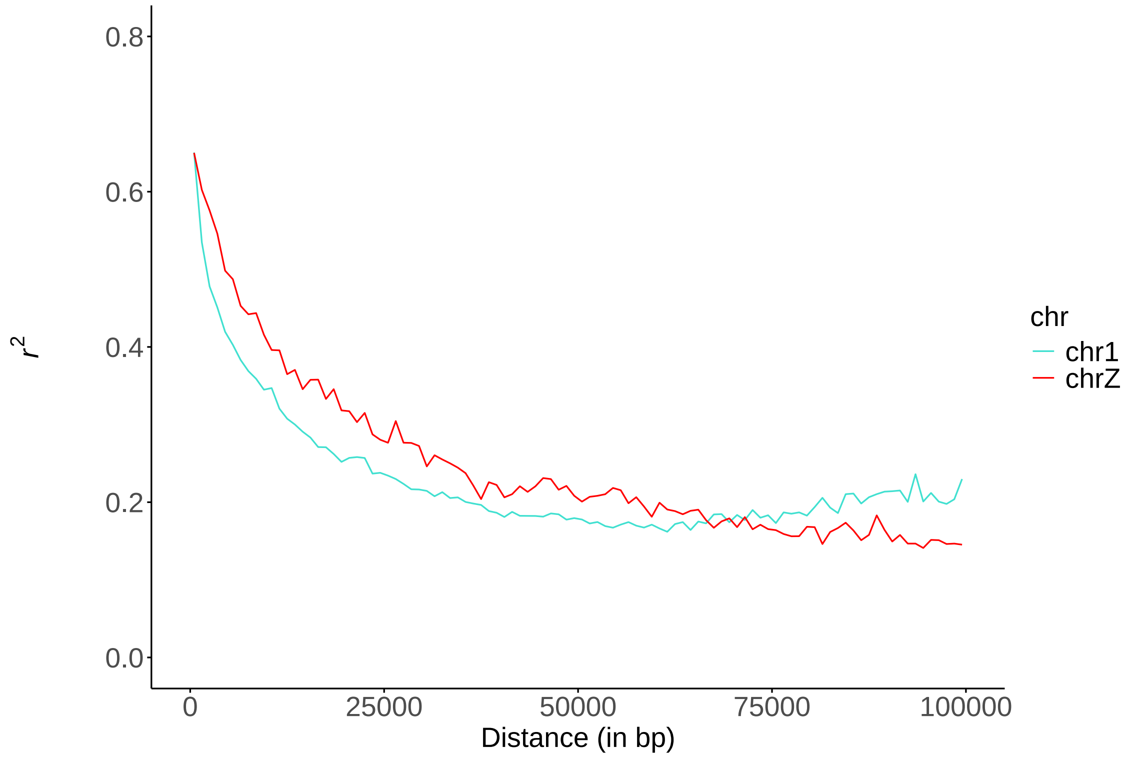


Figure S4: LD decay (in kb) in the LSP individuals. The blue curve represents the LD decay for chromosome 1 and the red curve LD decay for chromosome Z.

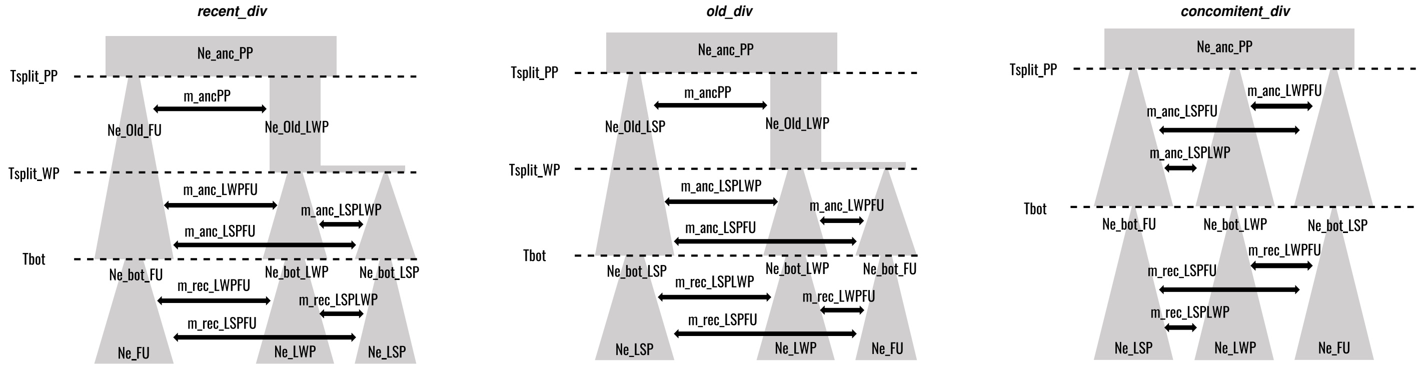


Figure S5: We simulated data under three distinct demographic scenarios. In the recent_div scenario, the divergence between LSP and LWP occurred after the divergence between LWP and FU. Both LWP and FU originated from an ancient population, Ancpp, while the current LWP and LSP populations originated from an ancient LWP population. A bottleaneck event occurred simultaneously in all three populations following the divergence between LSP and LWP. In the old_div scenario, the divergence between LSP and LWP occurred after the divergence between LWP and FU. Both LSP and LWP originated from an ancient population, Ancpp, while the current LWP and FU populations originated from an ancient LWP population. A bottleneck event occurred simultaneously in all three populations following the divergence between LWP and FU. In the concomitant_div scenario, the three populations (LSP, LWP, and FU) diverged simultaneously from an ancient population, Ancpp. A bottleneck event occurred concurrently in all three populations. The recombination rate was fixed at 4cM/Mb, based on recombination rates observed in several closely related species of Lepidoptera (Shipilina *et al.*, 2022; i Torres *et al.*, 2022; Palahí I Torres *et al.*, 2023), and the mutation rate was fixed at $2.9 \times 10^{-9}$ mutations per generation per SNP, as recently estimated for the Lepidoptera *Heliconius melpomene* (Keightley *et al.*, 2015) and used by Leblois *et al.* (2018). All parameters estimated in these scenarios and their priors are provided in Table S4.

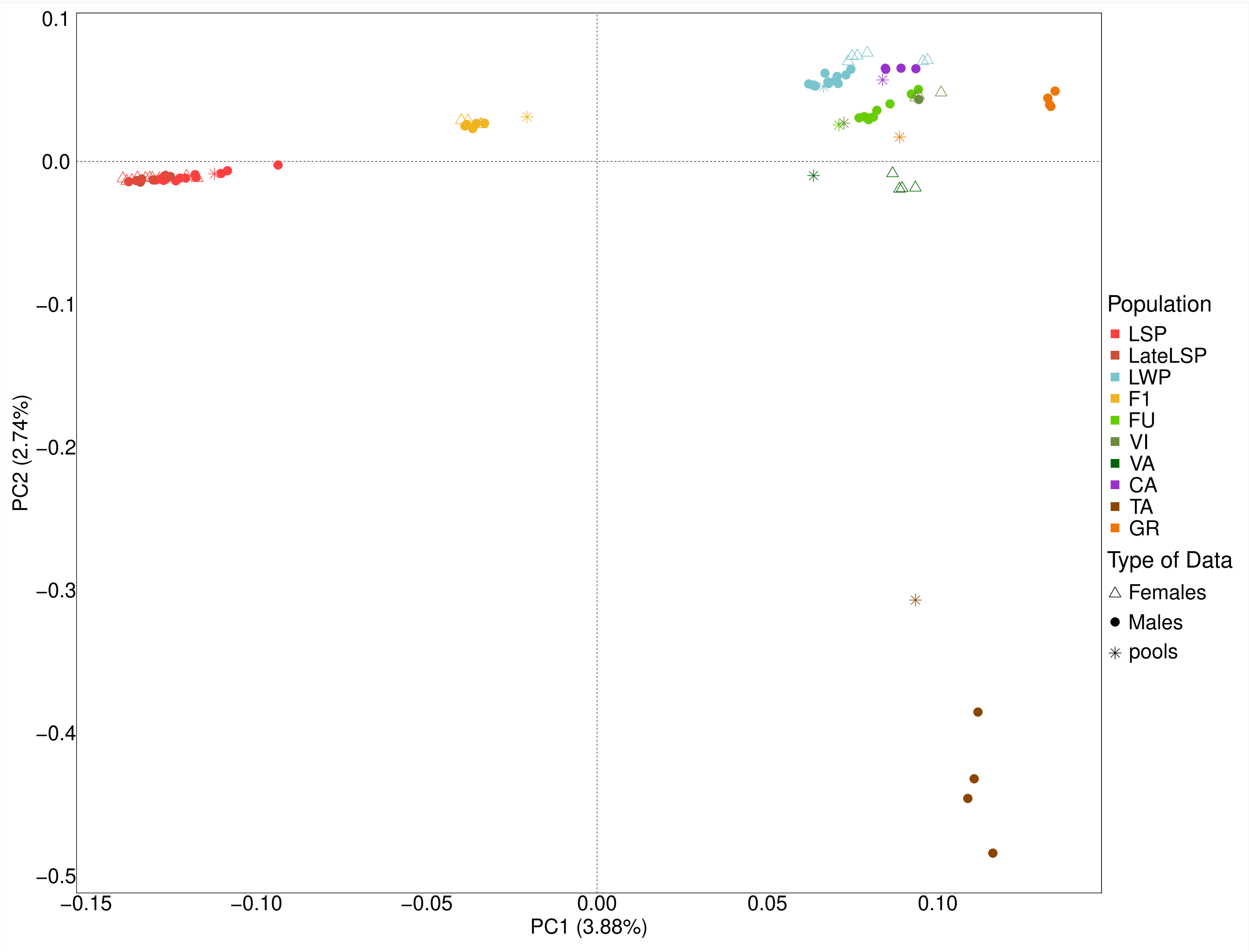


Figure S6: PCA on autosomes for 90 individuals and 10 pools from 8 Portuguese populations.

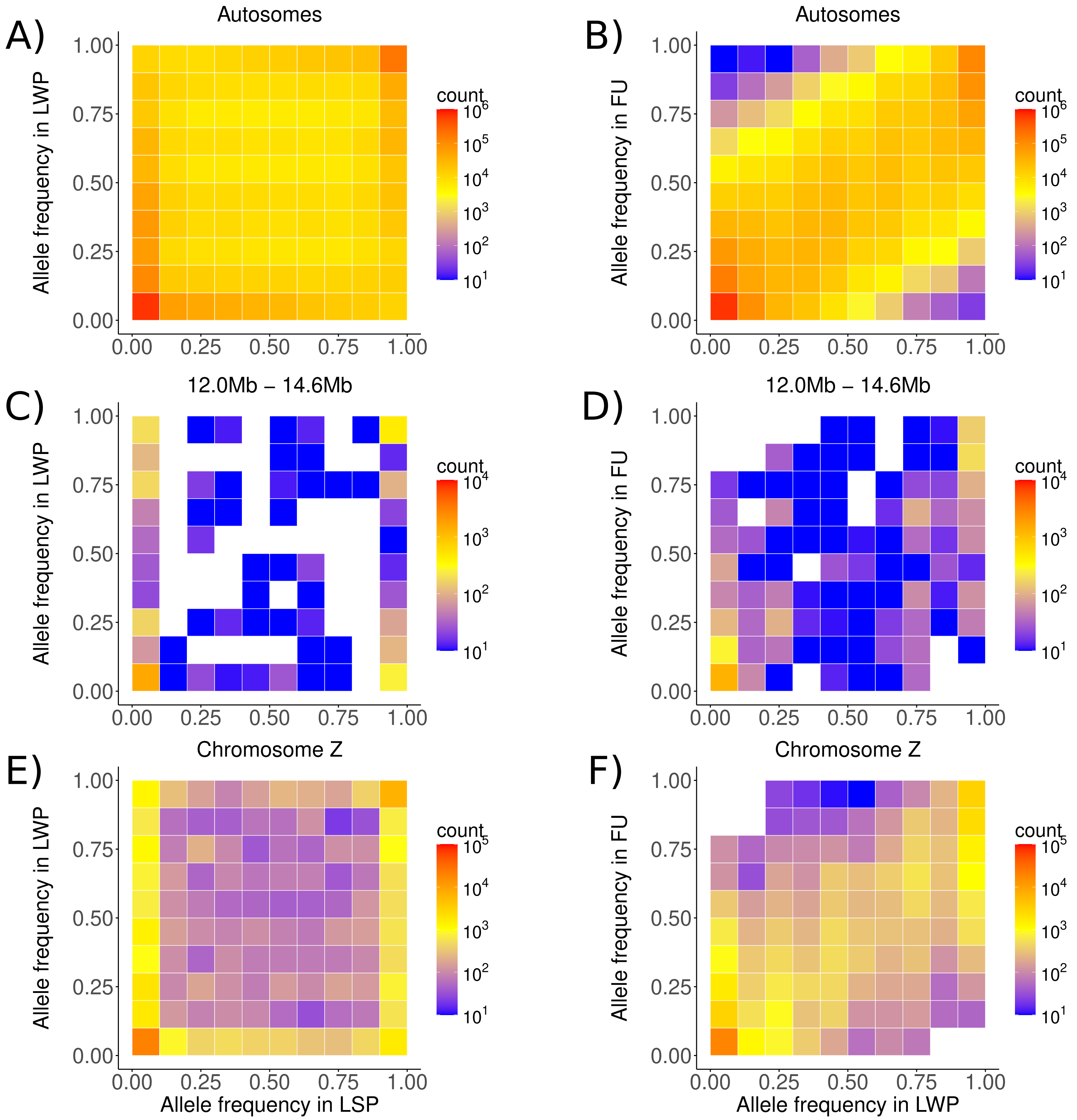


Figure S7: Two-dimensional Site Frequency Spectrum (SFS) for well-polarized derived alleles. **(A)** Joint SFS for autosomes between LSP (25 individuals) and LWP (18 individuals). **(B)** Joint SFS for autosomes between LWP (18 individuals) and FU (10 individuals). **(C)** Joint SFS for the zoomed region from Figure 6 (12 Mb – 14.6 Mb) between LSP (15 male individuals) and LWP (12 male individuals). **(D)** Joint SFS for the same zoomed region between LWP (12 male individuals) and FU (10 male individuals). **(E)** Joint SFS for the entire Z chromosome between LSP (15 male individuals) and LWP (12 male individuals). **(F)** Joint SFS for the entire Z chromosome between LWP (12 male individuals) and FU (10 male individuals).

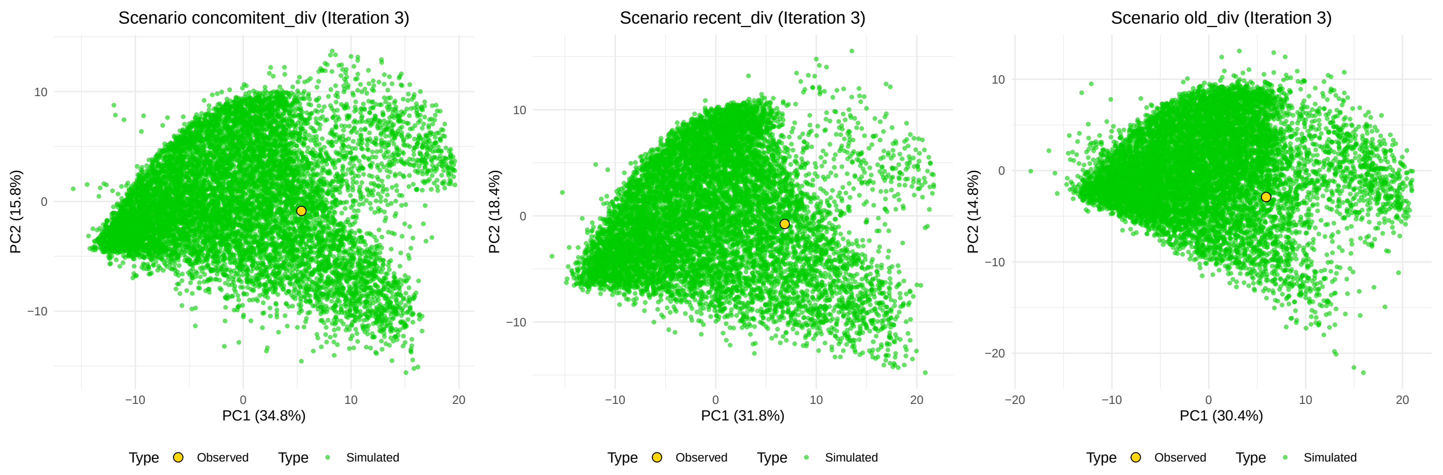


Figure S8: PCA of the simulated data (green dots) and observed data (yellow dot) for the iteration with the lowest local error rate (Table S7) for the concomitent_div, recent_div, and old_div scenarios.

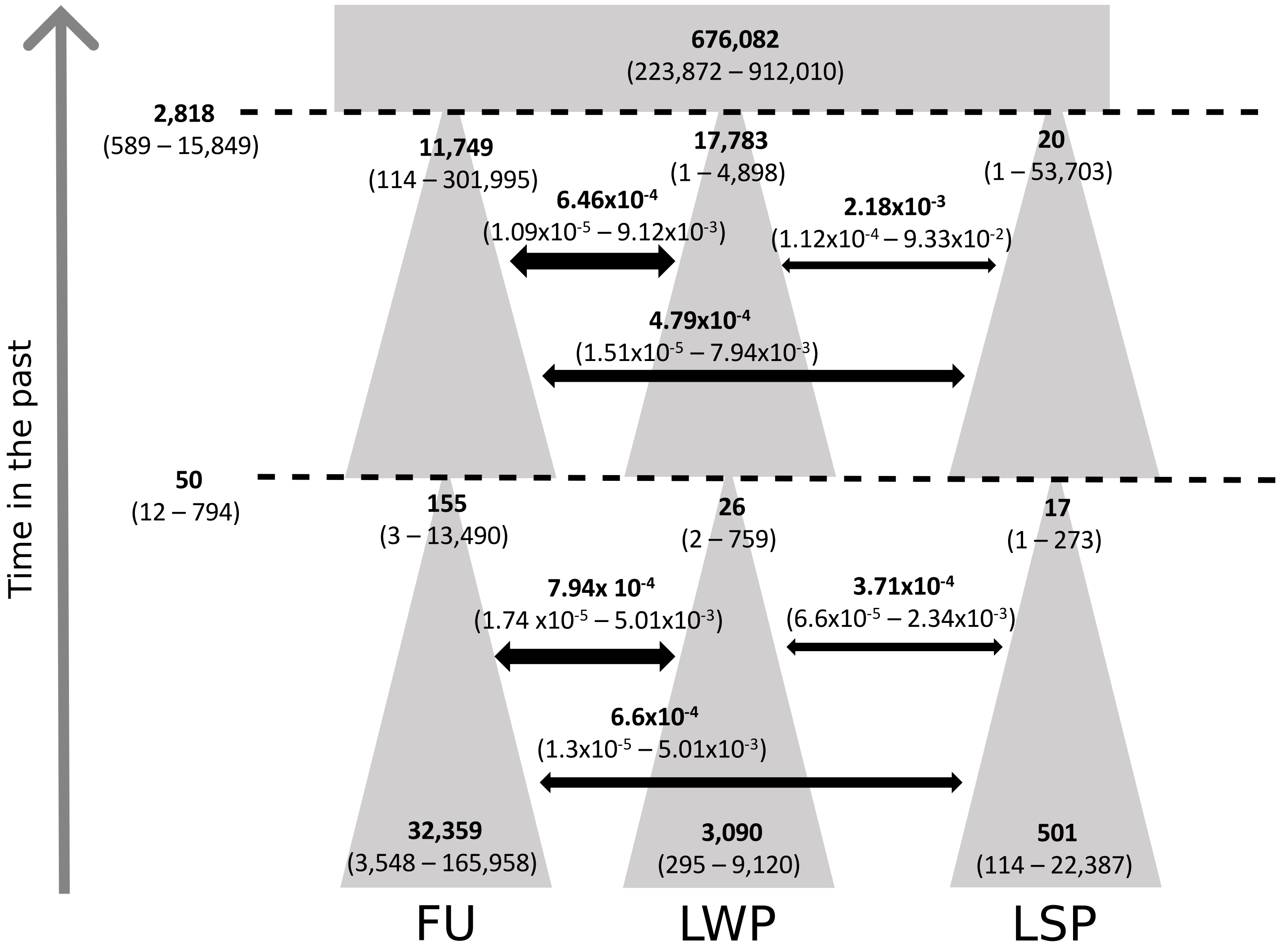


Figure S9: Graphical representation of the divergence among LSP, LWP, and FU populations under the concomitent_div scenario. Parameters were estimated from 100,000 simulations, highlighting a recent bottleneck in the 3 populations. Bold values correspond to the median estimates of the parameters, while the numbers in parentheses represent the 95% confidence interval (lower and upper bounds). The arrow correspond to time in the past.

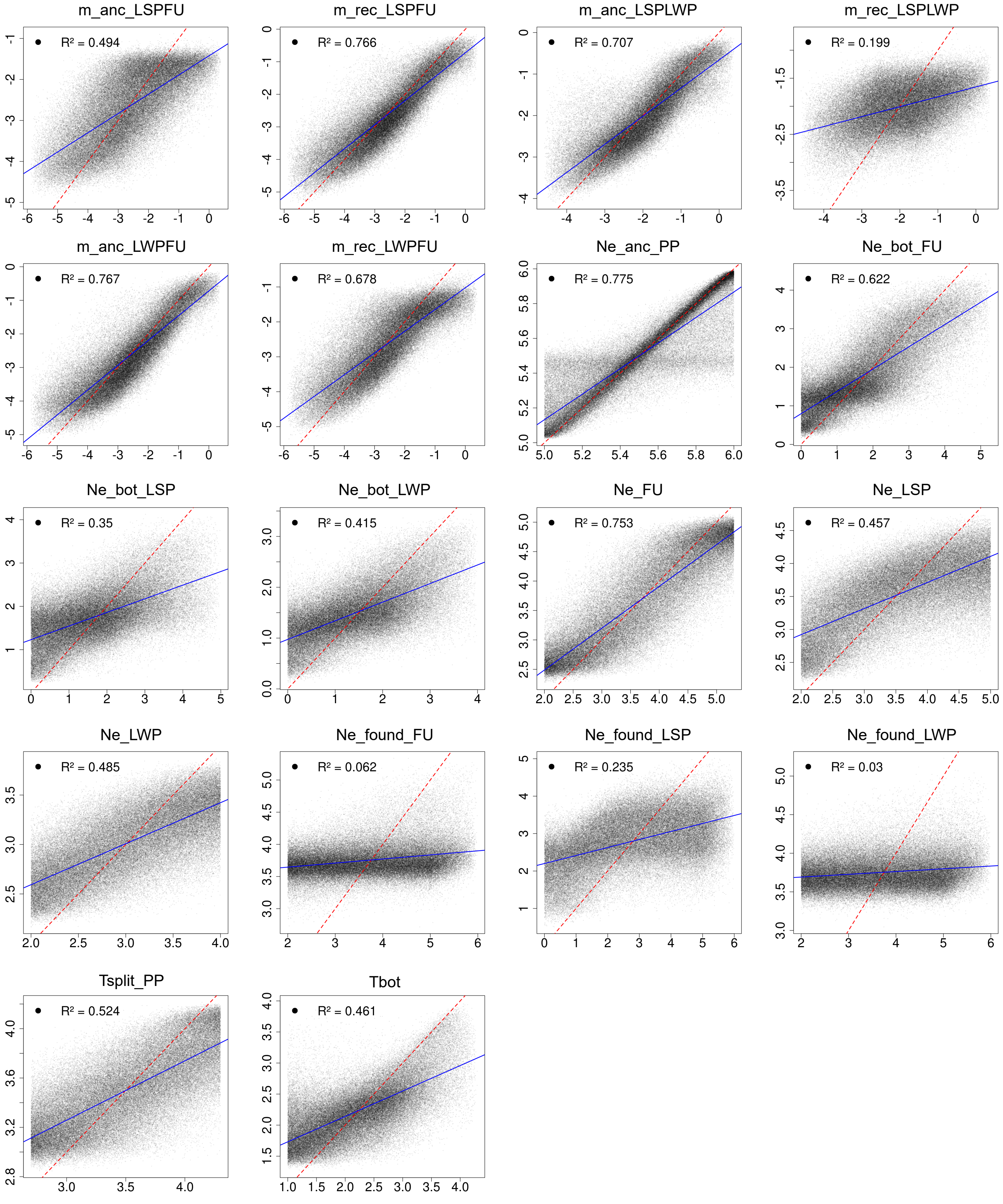


Figure S10: Out-of-bag prediction performance for ABC Random Forest for each parameter inference for the concomitent_div scenario. Points show the relationship between parameters values used to generate out-of-bag simulations (x-axis) and the corresponding predicted parameter values(y-axis), with both on a log10 scale. The red dashed line indicates perfect prediction (identity line), and the blue line represents the linear regression fit. $R^2$ indicates the model's goodness-of-fit.

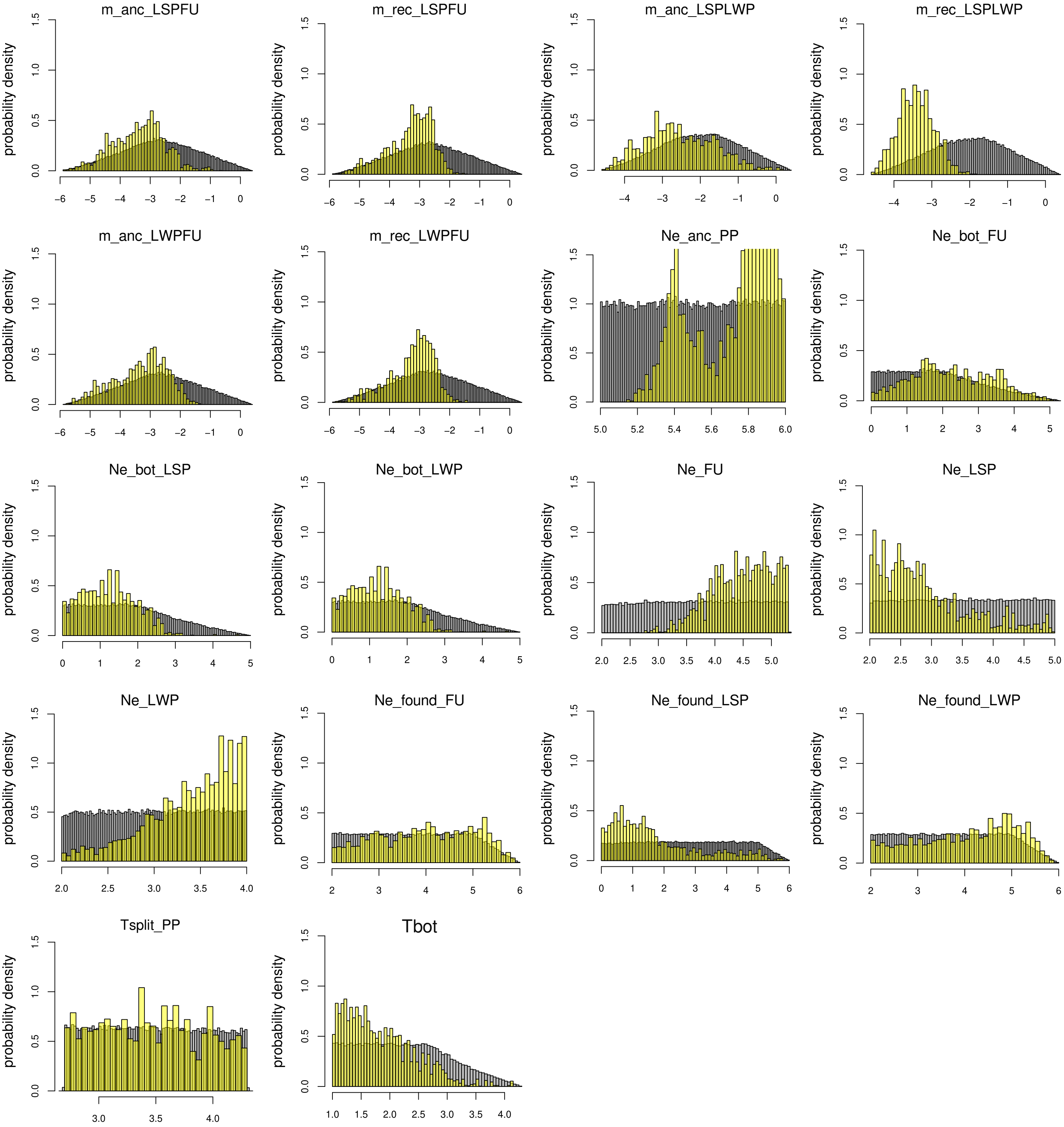


Figure S11: Posterior probability weight after the ABCRF analysis for the concomitant_div scenario. The grey histogram corresponds to the prior distribution of each estimated parameter. The yellow histogram corresponds to the posterior distribution weight of each estimated parameter after the ABCRF analysis.

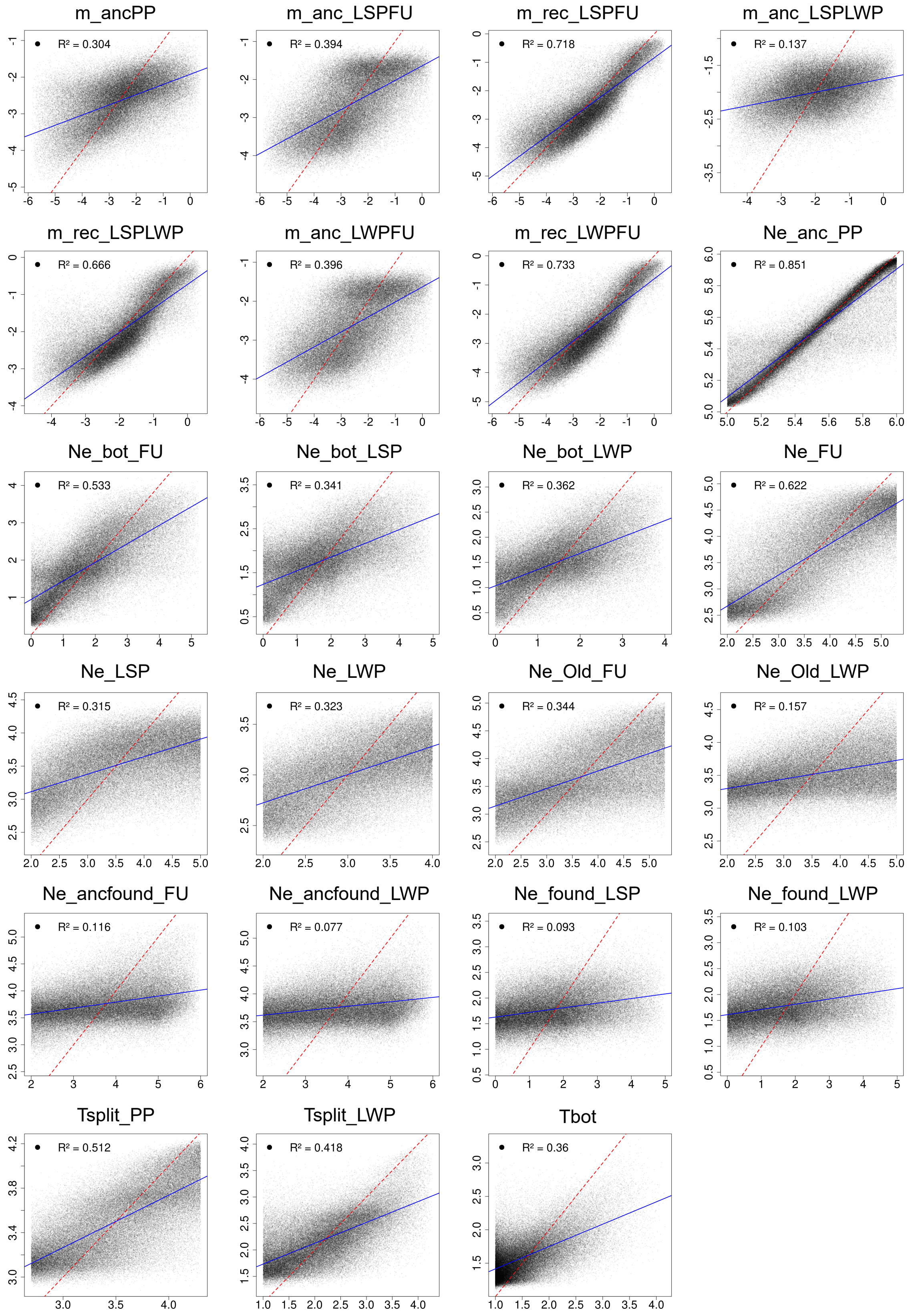


Figure S12: Out-of-bag prediction performance for ABC Random Forest for each parameter inference for the recent_div scenario. Points show the relationship between parameters values used to generate out-of-bag simulations (x-axis) and the corresponding predicted parameter values(y-axis), with both on a log10 scale. The red dashed line indicates perfect prediction (identity line), and the blue line represents the linear regression fit. $R^2$ indicates the model's goodness-of-fit.

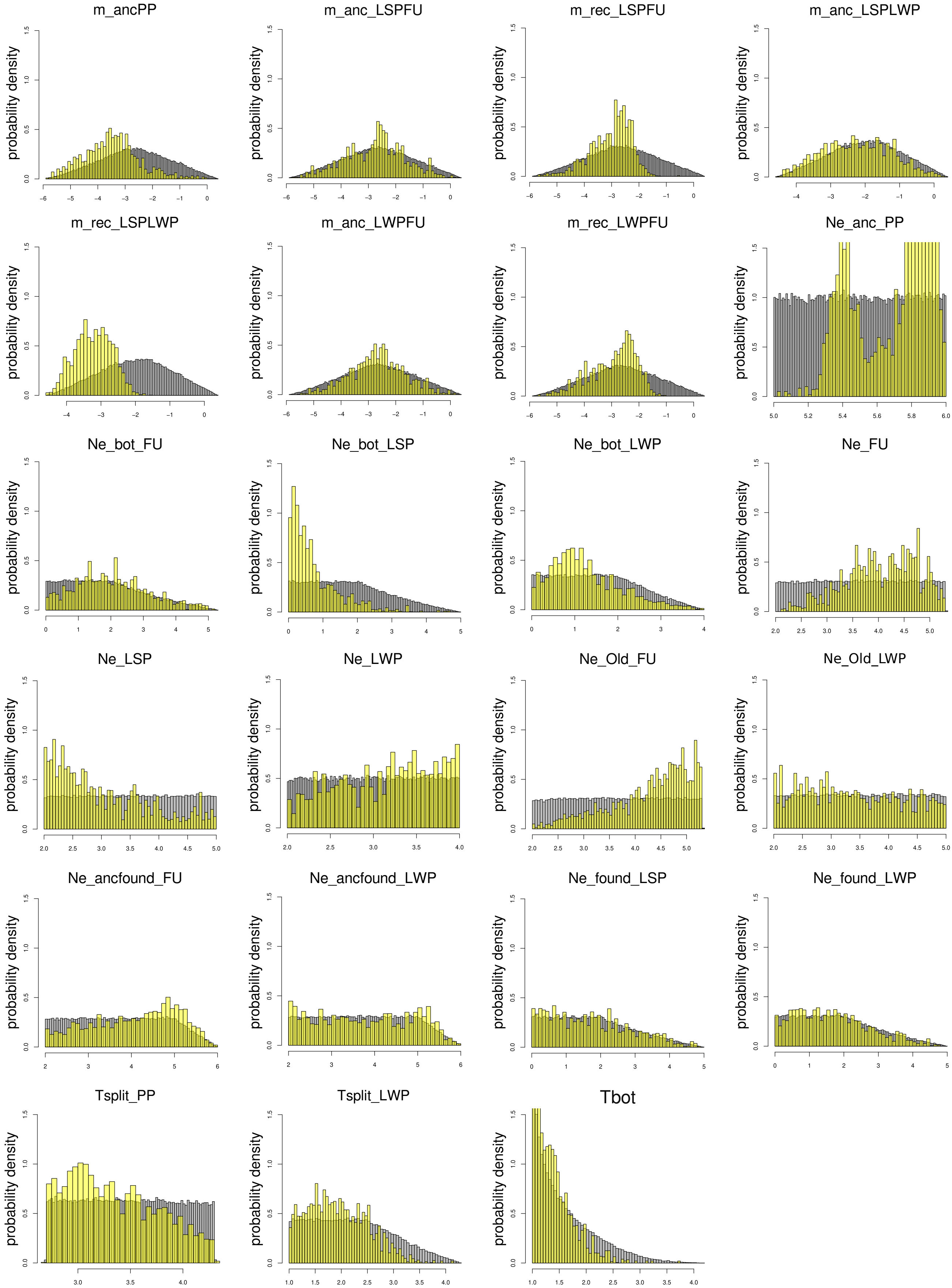


Figure S13: Posterior probability weight after the ABCRF analysis for the recent_div scenario. The grey histogram corresponds to the prior distribution of each estimated parameter. The yellow histogram corresponds to the posterior distribution weight of each estimated parameter after the ABCRF analysis.

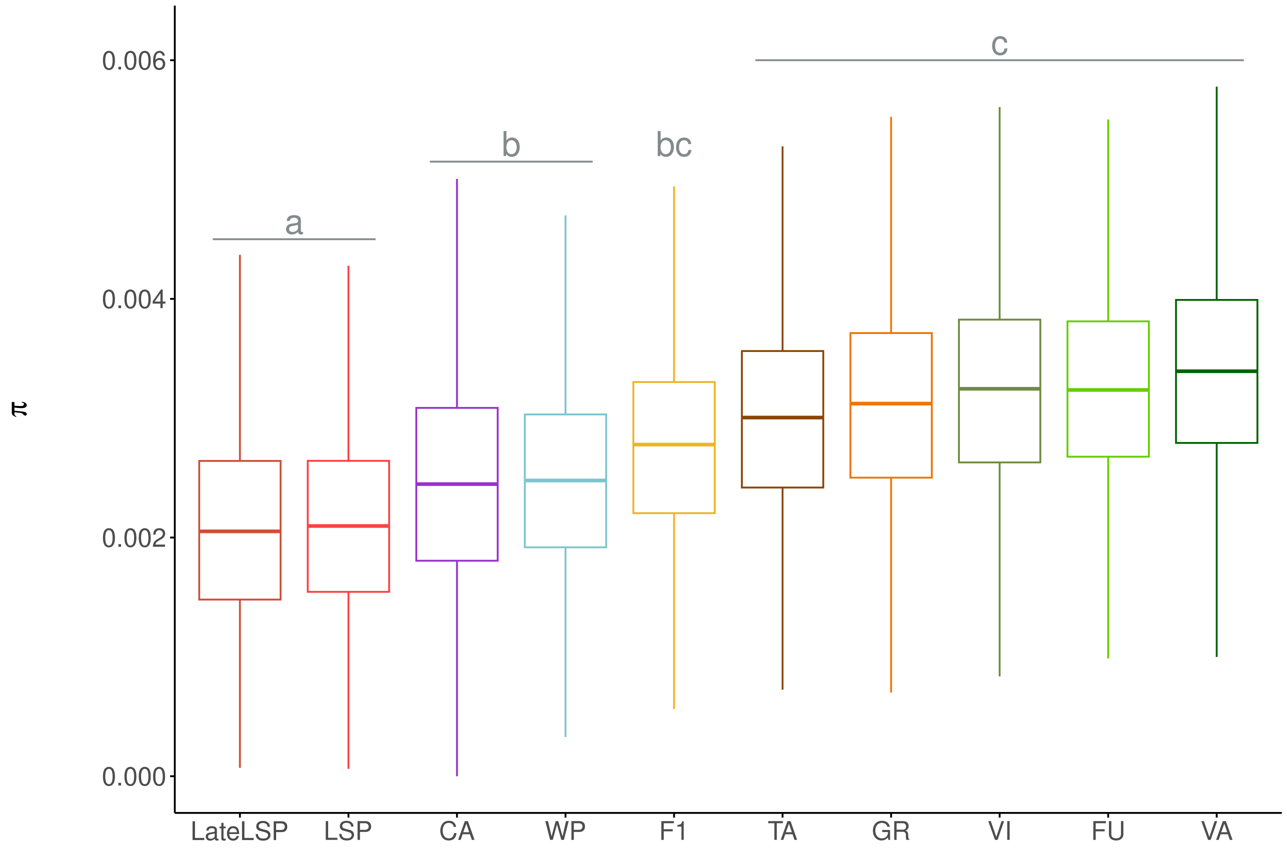


Figure S14: Boxplots of mean individual nucleotide diversity on autosomes for each population (with populations ordered by the median $\pi$ values). Post-hoc tests were conducted using estimated marginal means (i.e., least-squares means or adjusted means) with Bonferroni adjustment, in order to identify pairs of populations with significantly different $\pi$ values. Statistically significant group differences were determined by the letter from *a* to *c*.

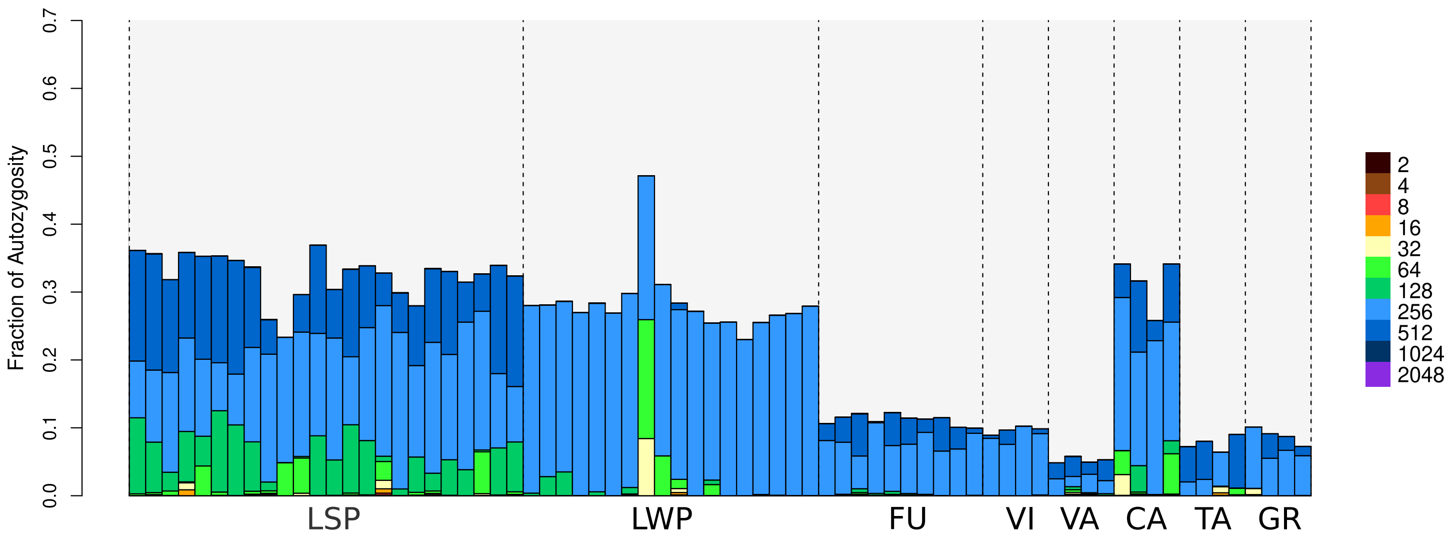


Figure S15: Genomic inbreeding inferred by RZooRoH, with each line represents the mean value of the individual genomic inbreeding coefficient for each population. The X-axis represents the generations of ancestors contributing to each HBD class. Only HBD classes ranging from 2 to 2048 generations are shown.

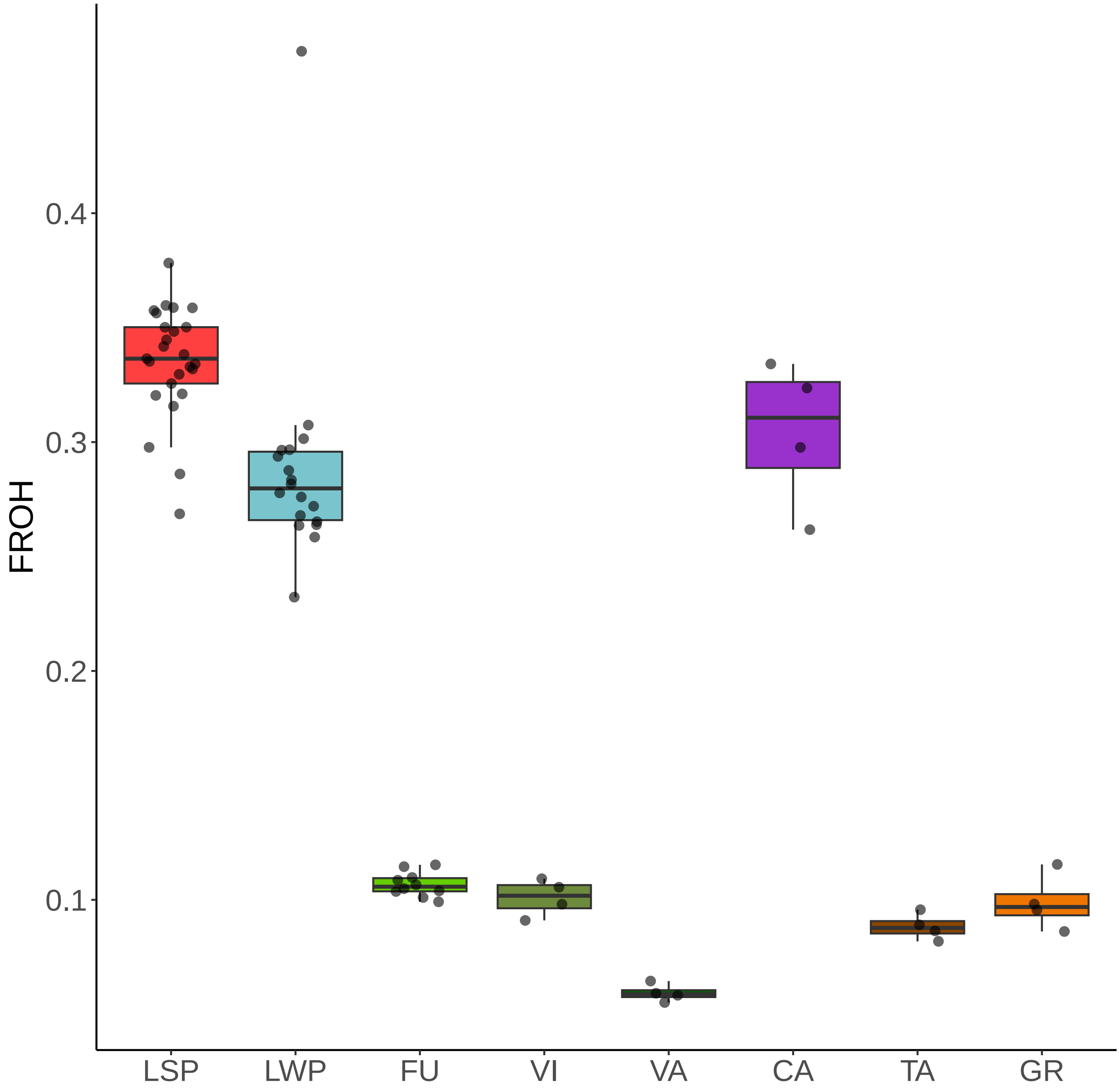


Figure S16: Fraction of the genome covered by ROH>100Kb (inferred by Plink) for each population. Color coding is the same as in Figure 1A. Each point corresponds to an individual.

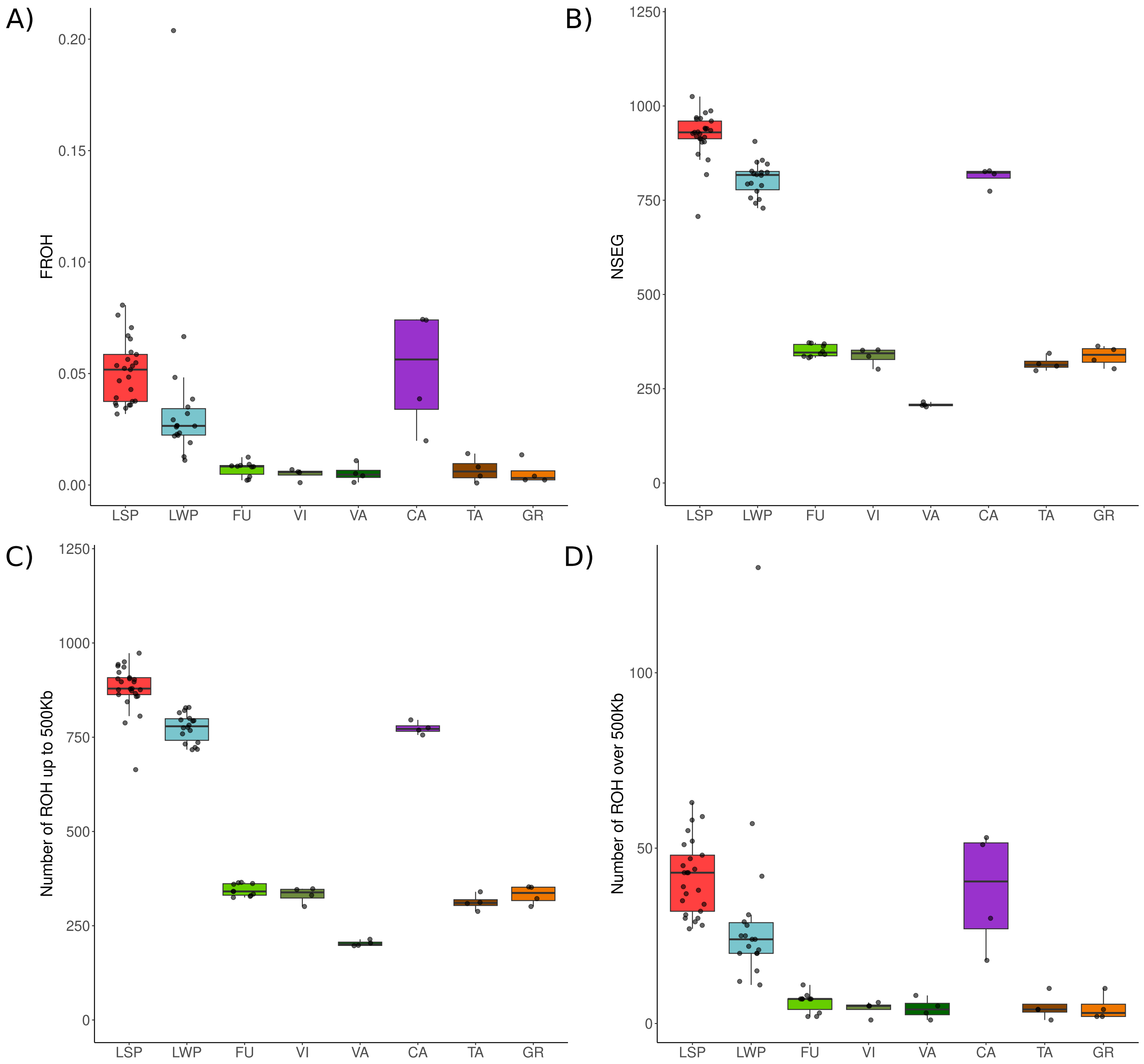


Figure S17: **(A)** Fraction of the genome covered by ROH > 500Kb (inferred by Plink) for each population. Color coding is the same as in Figure 1A. Each point corresponds to an individual. **(B)** The number of runs of homozygosity in total, **(C)** the number of runs of homozygosity up to 500kb, and **(D)** the number of runs of homozygosity over 500kb.

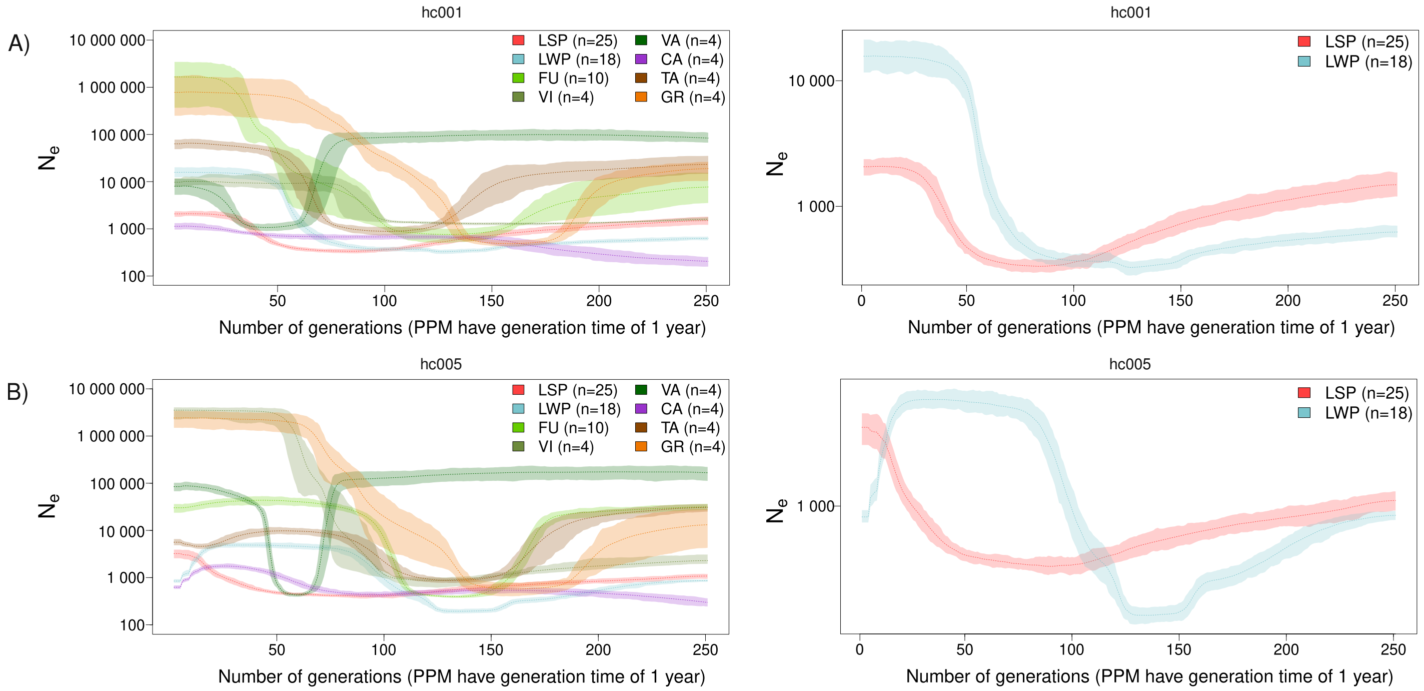


Figure S18: Recent population size history estimated using the *GONe* program (Santiago et al. 2020) for the Portuguese populations, with a zoom on LSP and LWP, and the hc parameter set to **(A)** 0.01 and **(B)** 0.05. For each population, 100 *GONe* analyses were conducted, as *GONe* randomly samples 50,000 SNPs per chromosome. The average $N_e$ trajectories and the 95% confidence intervals, estimated from these 100 analyses, were plotted. Please note that $N_e$ estimates for generations older than 100 are less reliable (Santiago *et al.*, 2020).

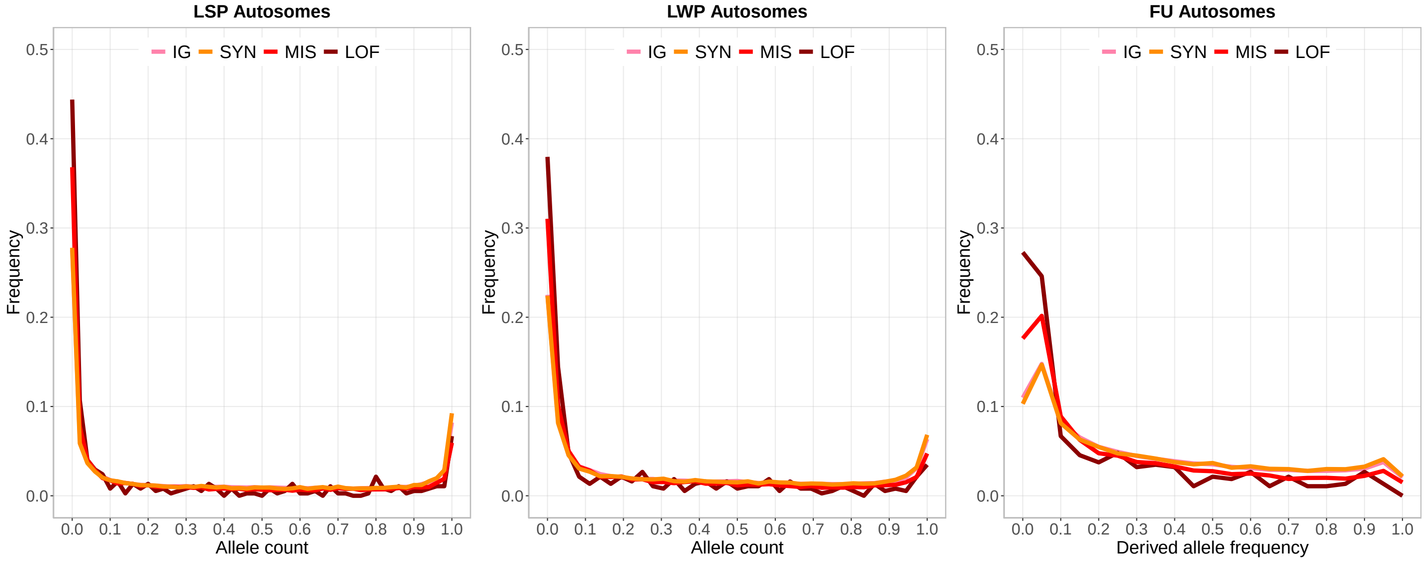


Figure S19: Comparison of the Site Frequency Spectrum (SFS) for intergenic (IG), Synonymous (SYN), Non-synonymous (MIS) and High impact (LOF) variants on autosomes in LSP, LWP and FU.

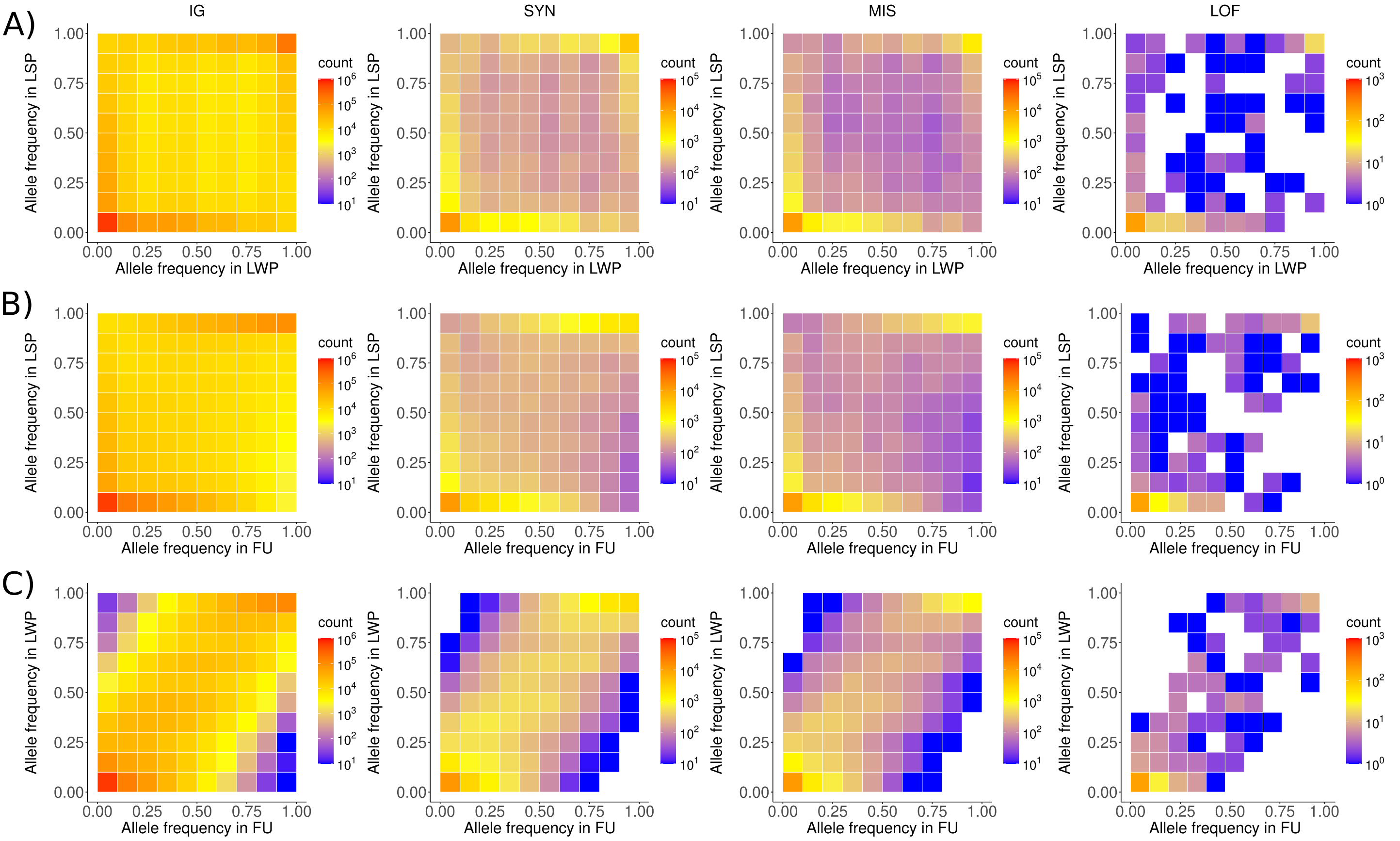


Figure S20: Two-dimensional Site Frequency Spectrum (SFS) for intergenic (IG), Synonymous (SYN), Non-synonymous (MIS) and High impact (LOF) variants. **(A)** Joint SFS for each category between LSP (25 individuals) and LWP (18 individuals). **(B)** Joint SFS for each category between LSP (25 individuals) and FU (10 individuals). **(C)** Joint SFS for each category between LWP (25 individuals) and FU (10 individuals).

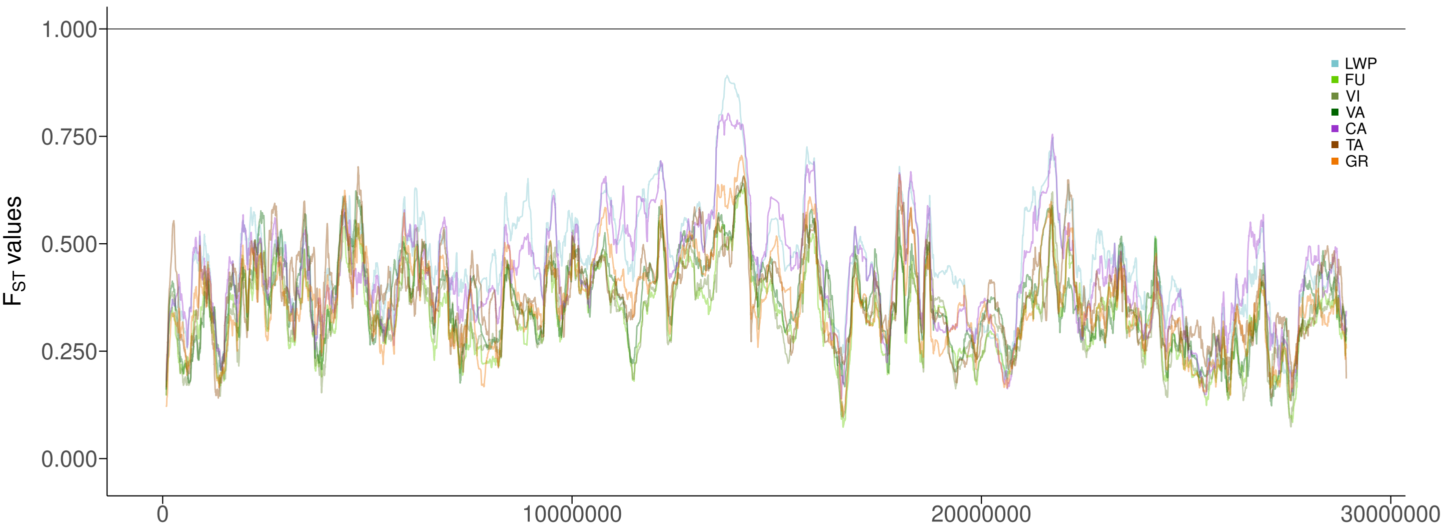


Figure S21: Pairwise $F_{\mathrm{ST}}$ (10-SNP windows) along the Z chromosome for all population pairs between LSP and a Portuguese population.

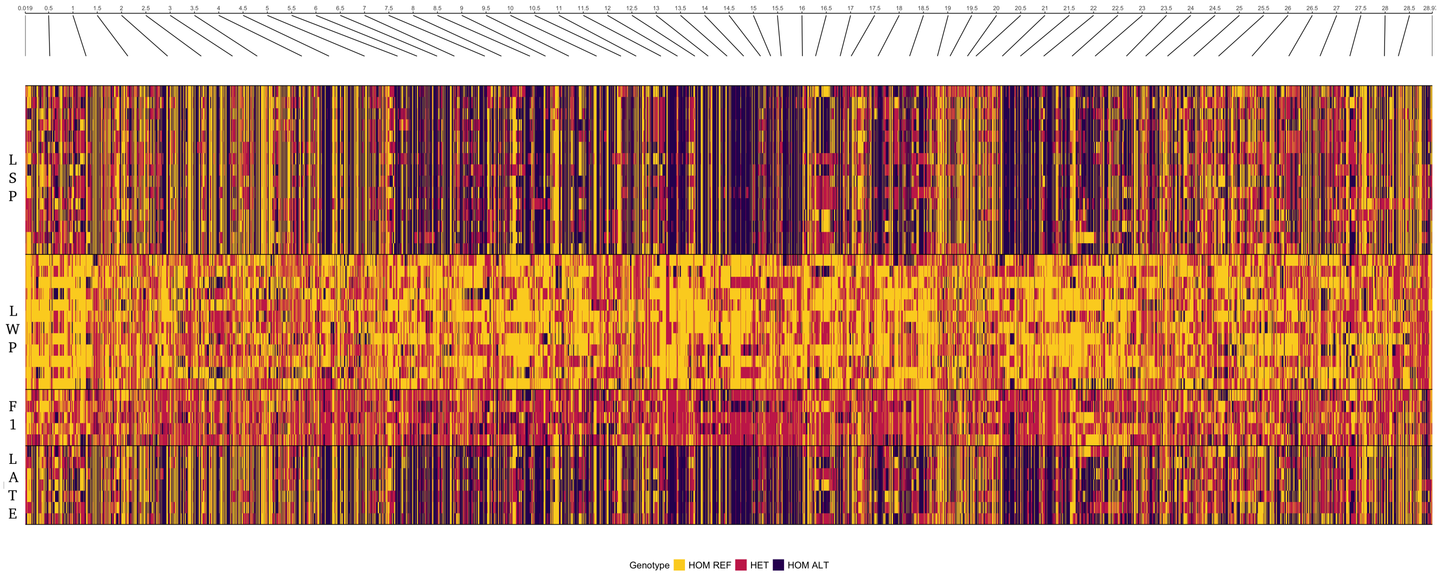


Figure S22: Genotype plot for LSP, LWP, F1 and LateLSP male genotypes, on the Z chromosome.

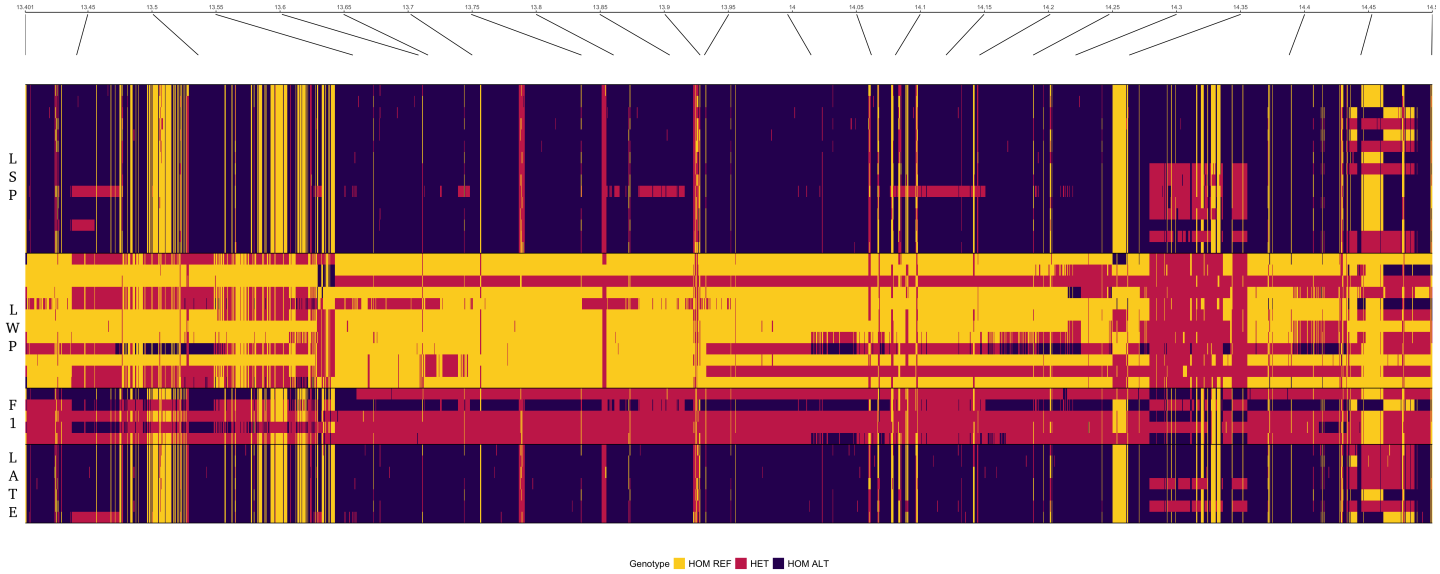


Figure S23: Genotype plot for LSP, LWP, F1 and LateLSP male genotypes, from 13.400 to 14.500 Mb on the Z chromosome.

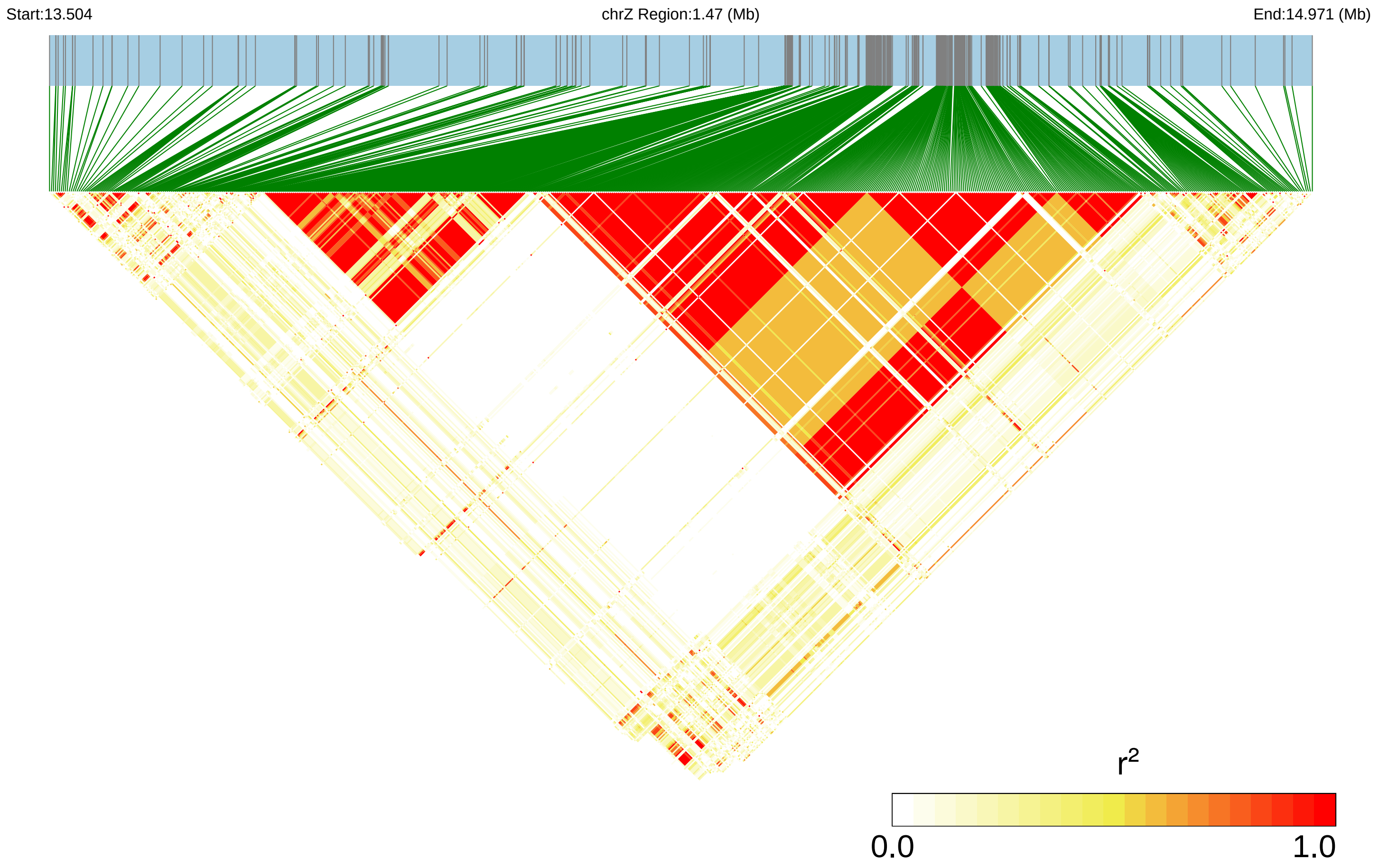


Figure S24: Linkage disequilibrium for SNPs found in the highly differentiated region spanning 13.5 Mb to 15.00 Mb on the Z chromosome.

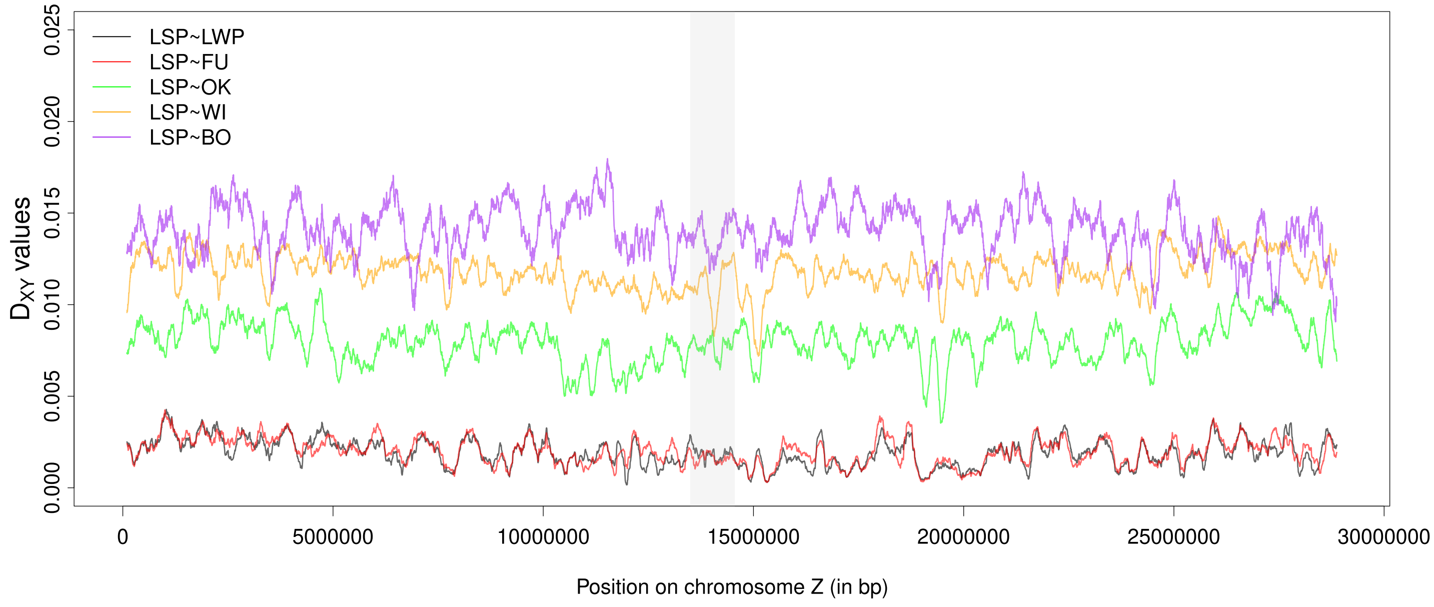


Figure S25: $D_{XY}$ values along the Z chromosome between LSP and Portuguese population and LSP and outgroups. The shaded zone corresponds to the highly differentiated region identified in Figure 4D between 13.55 Mb to 14.30 Mb.

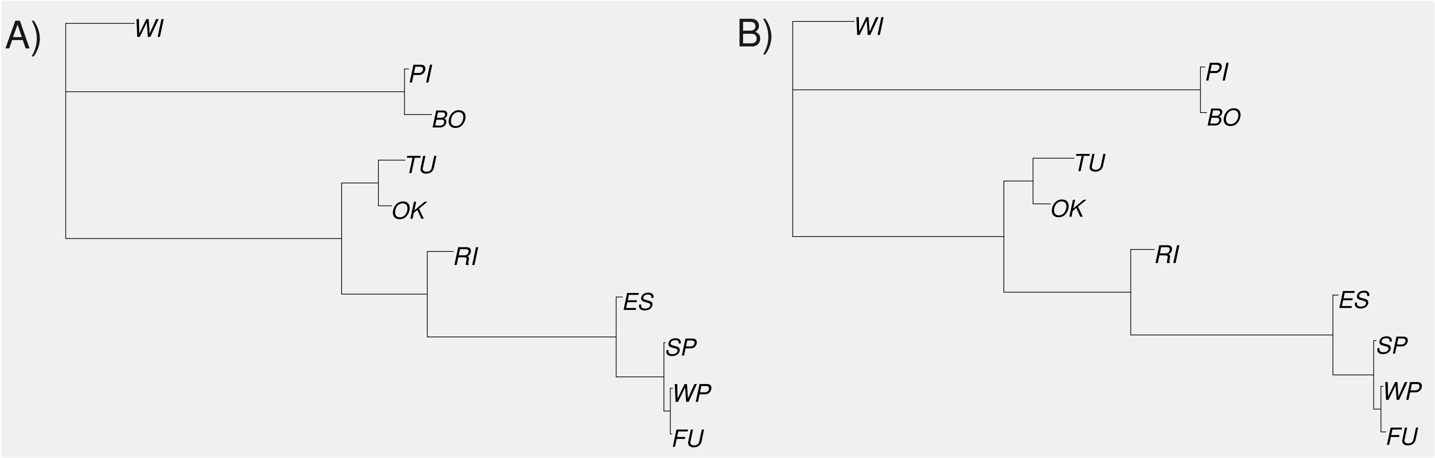


Figure S26: Graphical representation of the distance tree based on pairwise $D_{XY}$ values across the entire Z chromosome and the distance tree based on pairwise $D_{XY}$ values within the highly differentiated region spanning 13.55 Mb to 14.30 Mb on the Z chromosome.

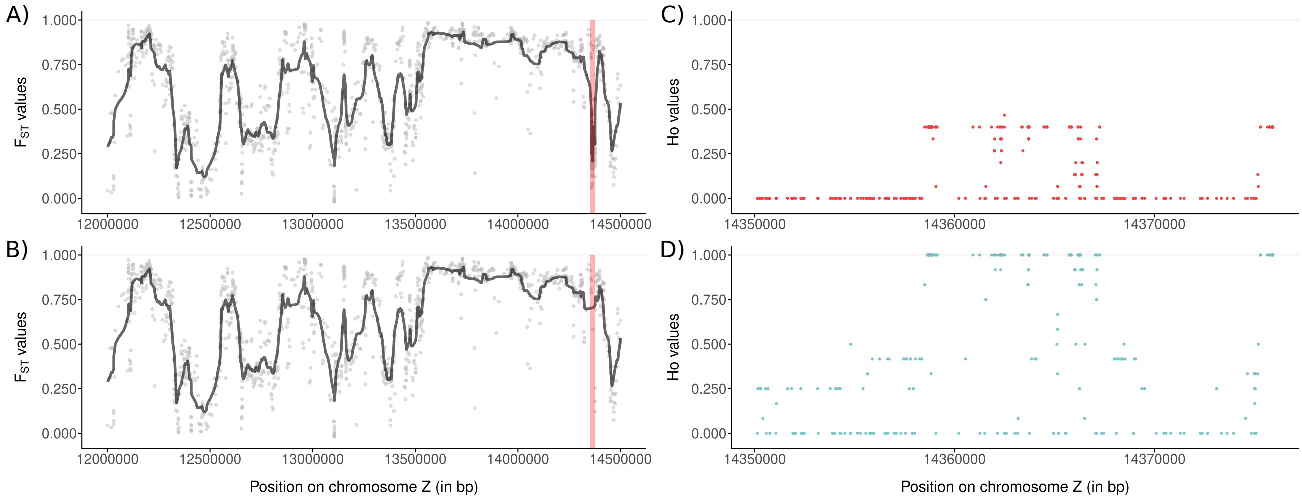


Figure S27: Pairwise $F_{ST}$ (10-SNP windows) values between LSP and LWP (Pool-Seq data) from 12.00 to 14.6 Mb, with (A) and without (B) a region (red rectangle) with excessive read depth (see figure 6K) and observed heterozygosity in LWP (D) compared to LSP (C).

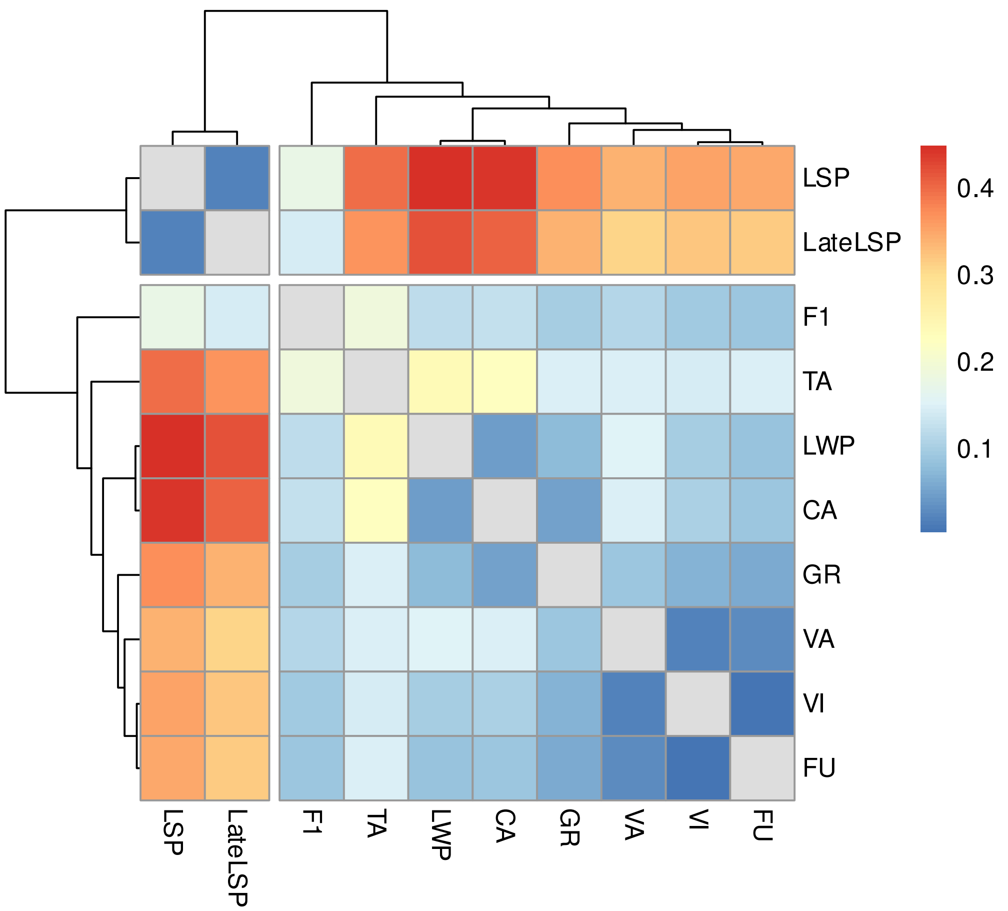


Figure S28: Pairwise $F_{ST}$ matrix between Portuguese populations, with values for the Z chromosome shown. Color gradients represent differentiation intensity, with darker red indicating higher differentiation and darker blue indicating lower differentiation.

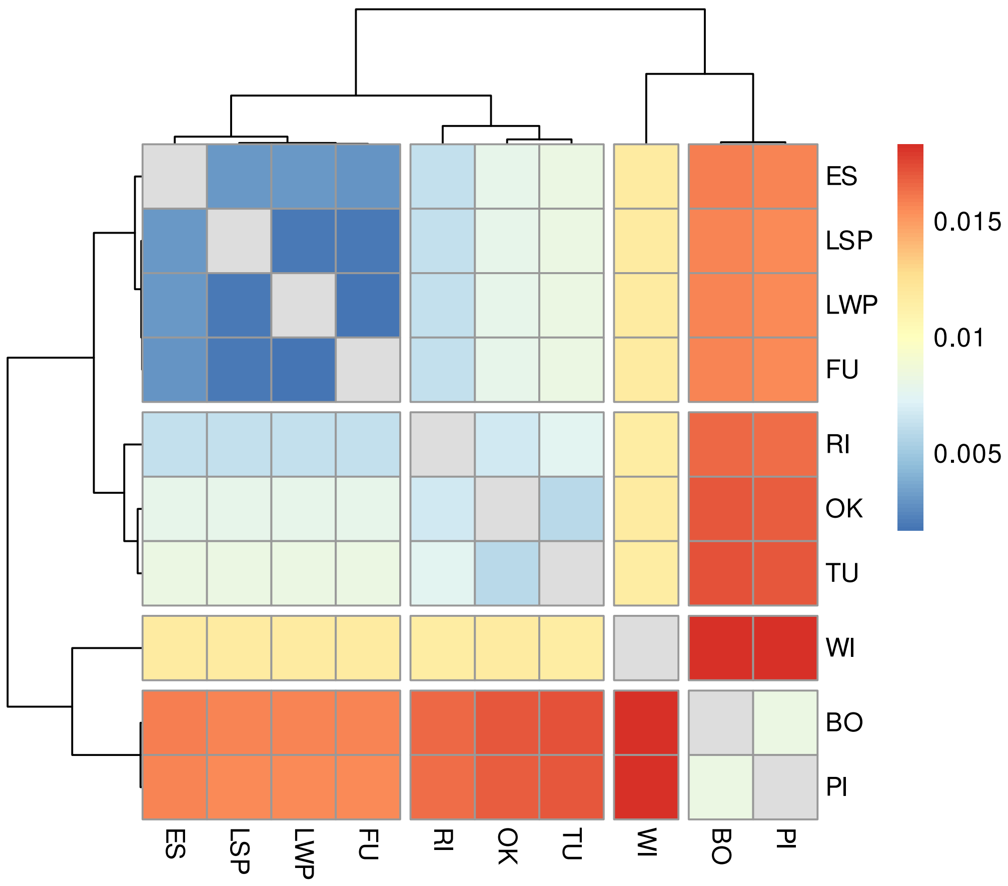


Figure S29: Pairwise $D_{\mathrm{XY}}$ matrix between Portuguese populations LSP, LWP, and FU and outgroups, with values for the Z chromosome shown. Color gradients represent divergence intensity, with darker red indicating higher divergence and darker blue indicating lower divergence.

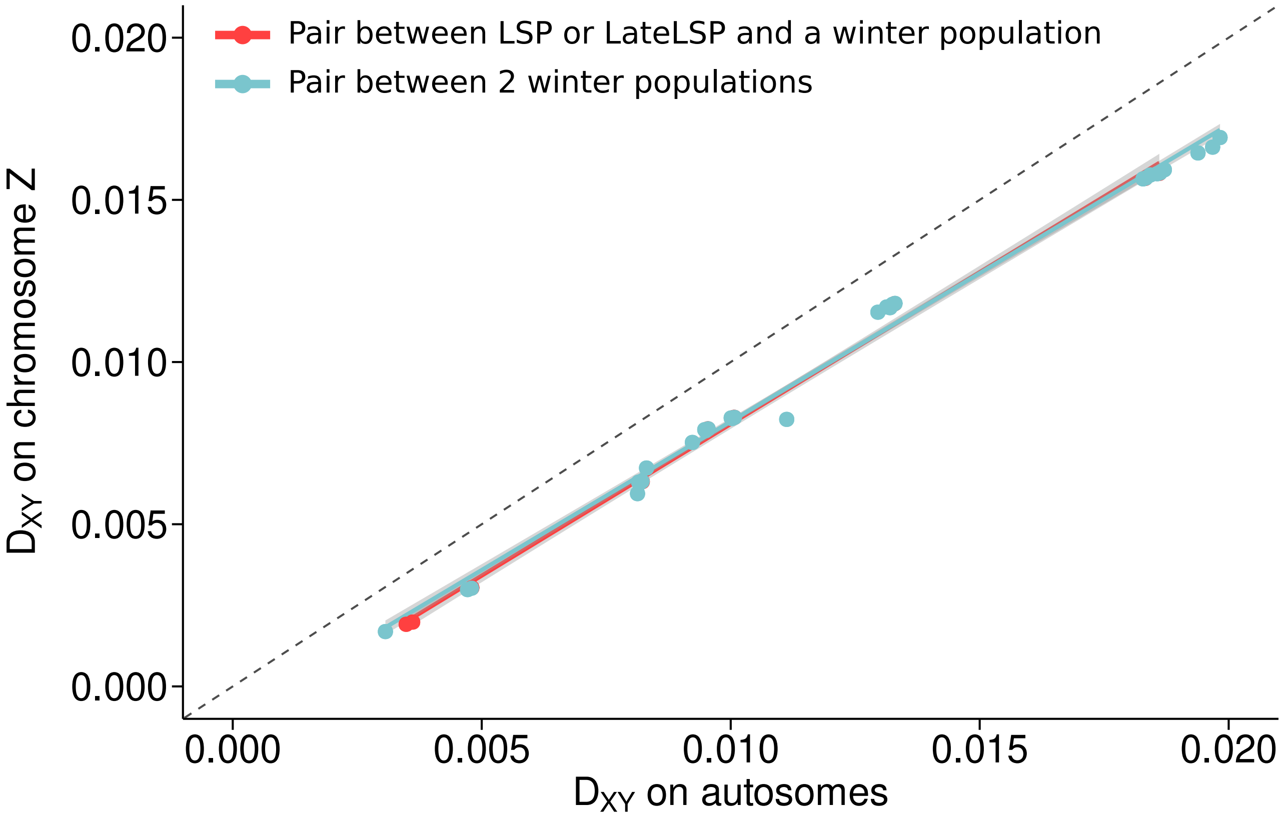


Figure S30: Pairwise $D_{XY}$ values calculated on the Z chromosome compared to pairwise $D_{XY}$ values calculated on autosomes. The red dots represent $D_{XY}$ values involving comparisons with LSP, and the blue dots indicate $D_{XY}$ values that do not involve LSP.

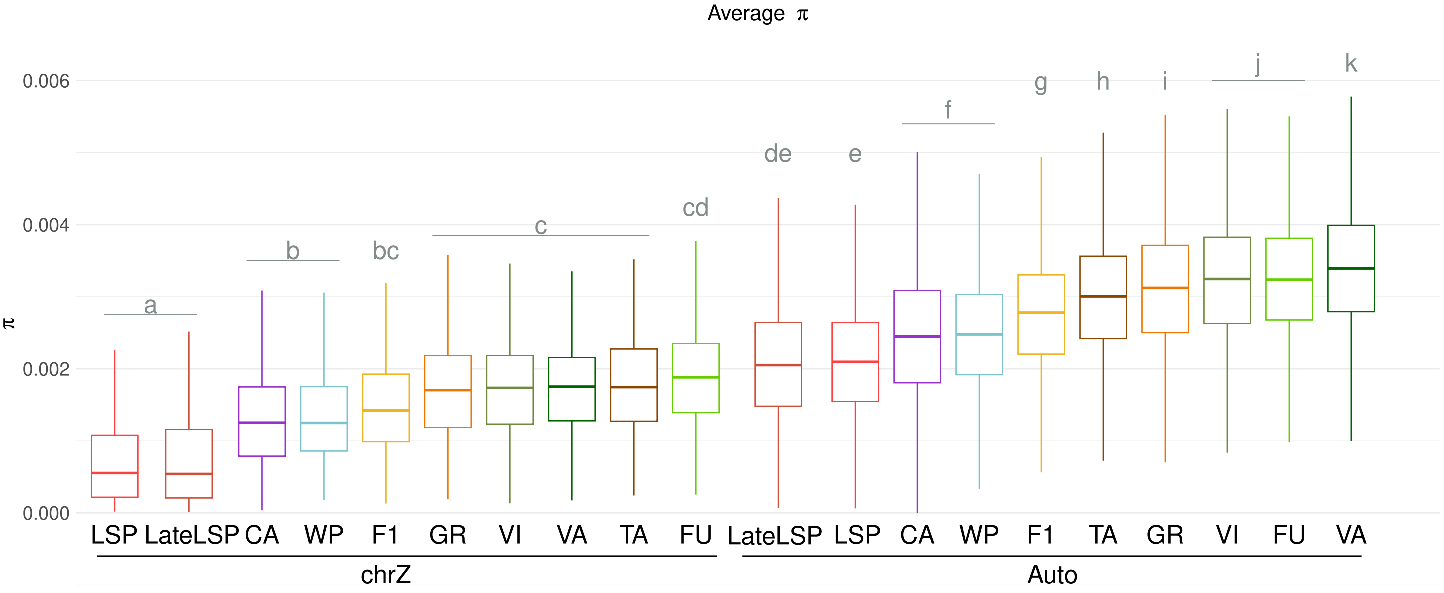


Figure S31: Nucleotide diversity on the Z chromosome (left) and on autosomes (right) for each individual in each population, ordered by the median nucleotide diversity values. Post-hoc tests were conducted using estimated marginal means (i.e., least-squares means or adjusted means) with Bonferroni adjustment, in order to identify pairs of populations with significantly different $\pi$ values. Statistically significant group differences were determined by the letter from a to k.

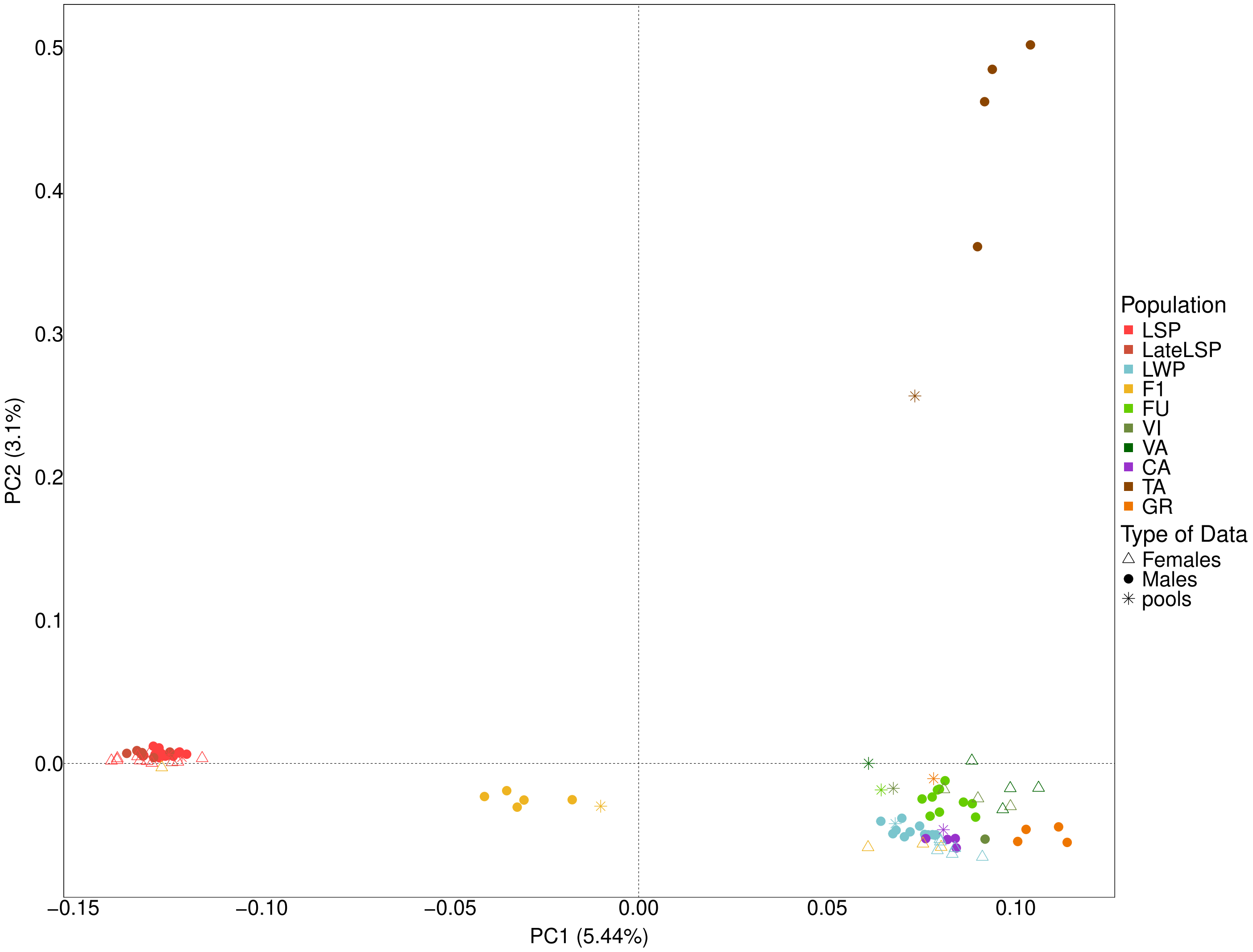


Figure S32: PCA on the Z chromosome for 90 individuals and 10 pools from 8 Portuguese populations.

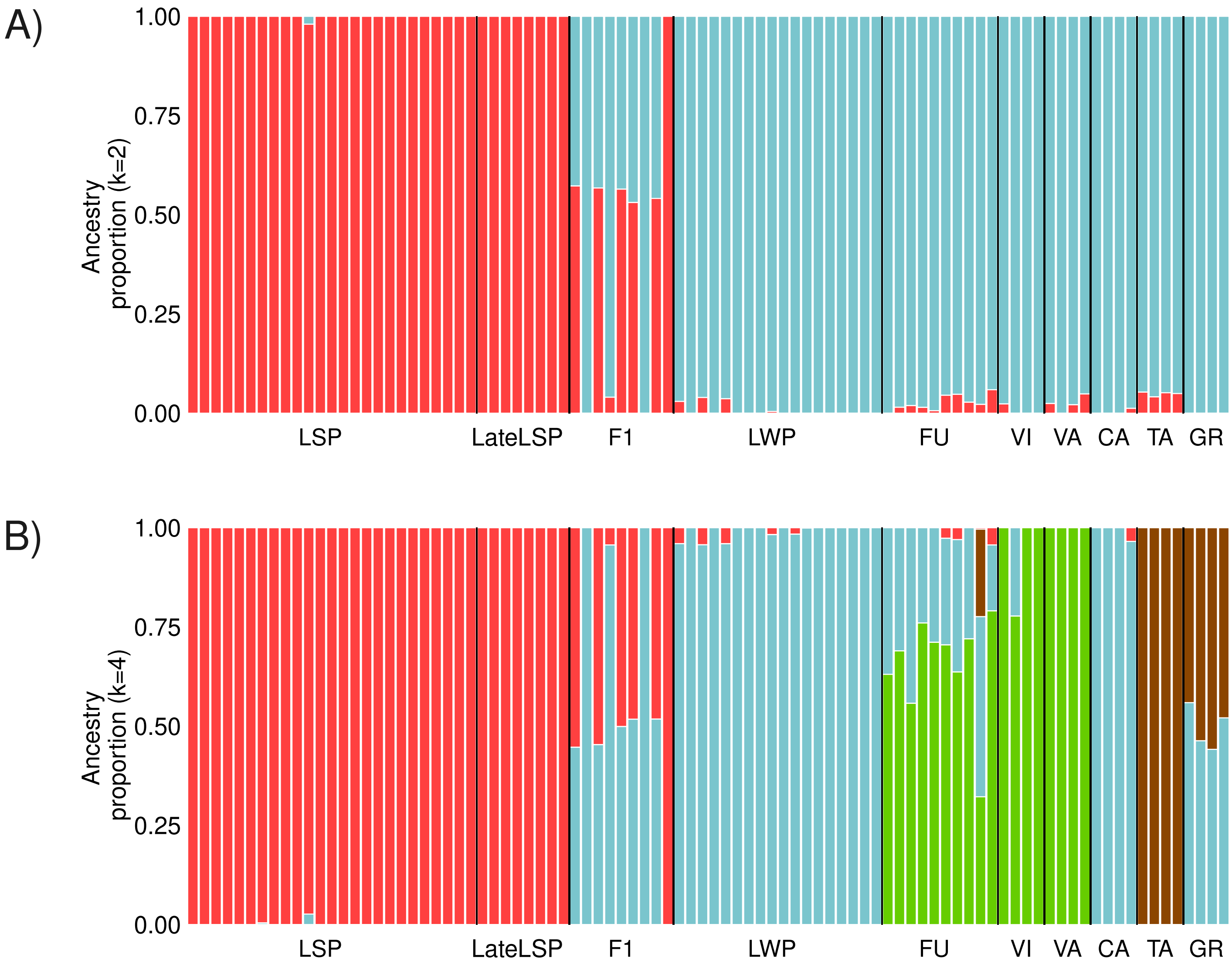


Figure S33: Admixture graph for the Z chromosome at k=2 (A) and at k=4 (B) (vertical lines correspond to individual admixture and colors correspond to distinct genetic groups).

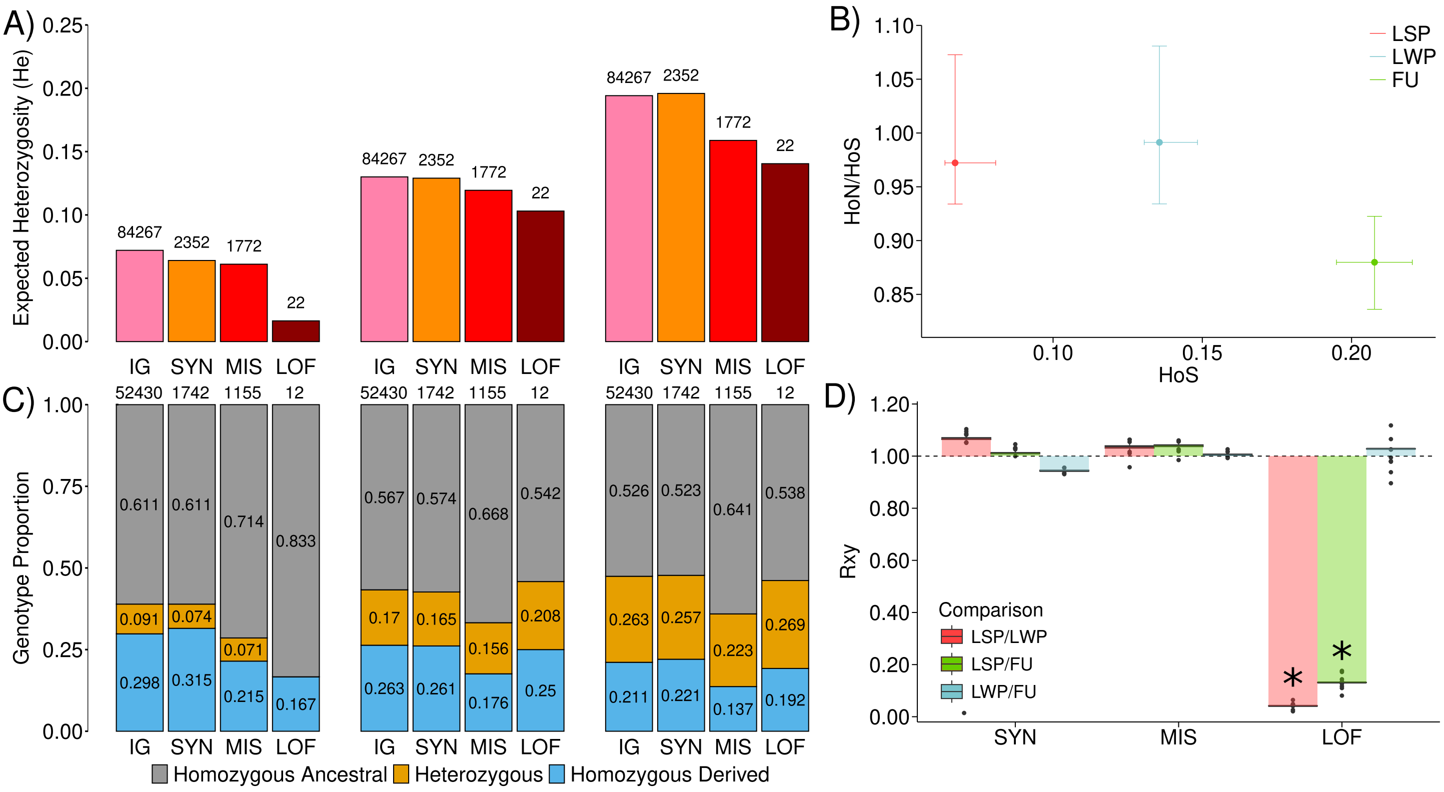


Figure S34: **(A)** Expected heterozygosities at IG, SYN, MIS, and LOF sites on the Z chromosome, with the count of variants indicated above each category. **(B)** Relationship between the ratio of individual heterozygosity at MIS (HoN) to SYN (HoS) sites on the Z chromosome and individual heterozygosity at SYN (HoS) sites on the Z chromosome. Each point represents the population median, with confidence intervals based on individual-level variation.**(C)** Proportions of genotypes that were either homozygous ancestral, homozygous derived or heterozygous, at IG, SYN, MIS, and LOF sites on the Z chromosome, with the count of variants indicated above each category. **(D)** $R_{XY}$ statistics at SYN, MIS, and LOF sites on the Z chromosome. The stars indicate that the $R_{XY}$ value is significantly different of 1 after using a z-score two-tailed test.

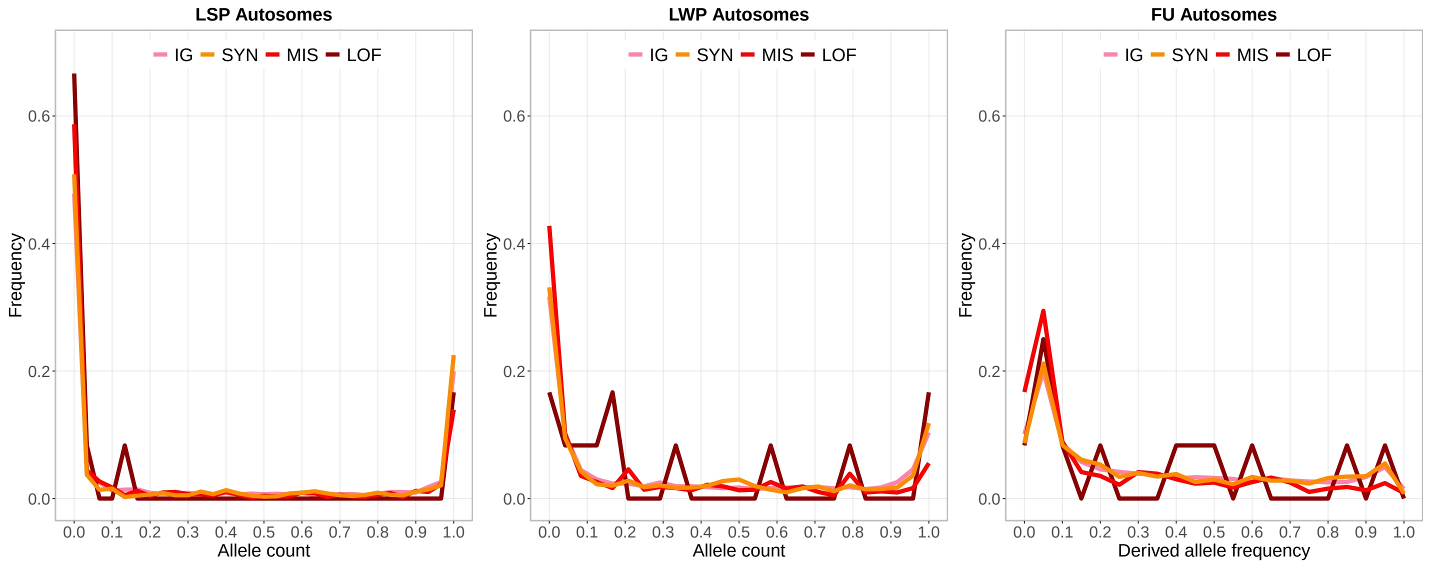


Figure S35: Comparison of the Site Frequency Spectrum (SFS) for intergenic (IG), Synonymous (SYN), Nonsynonymous (MIS) and High impact (LOF) variants on the Z chromosome in LSP, LWP and FU.

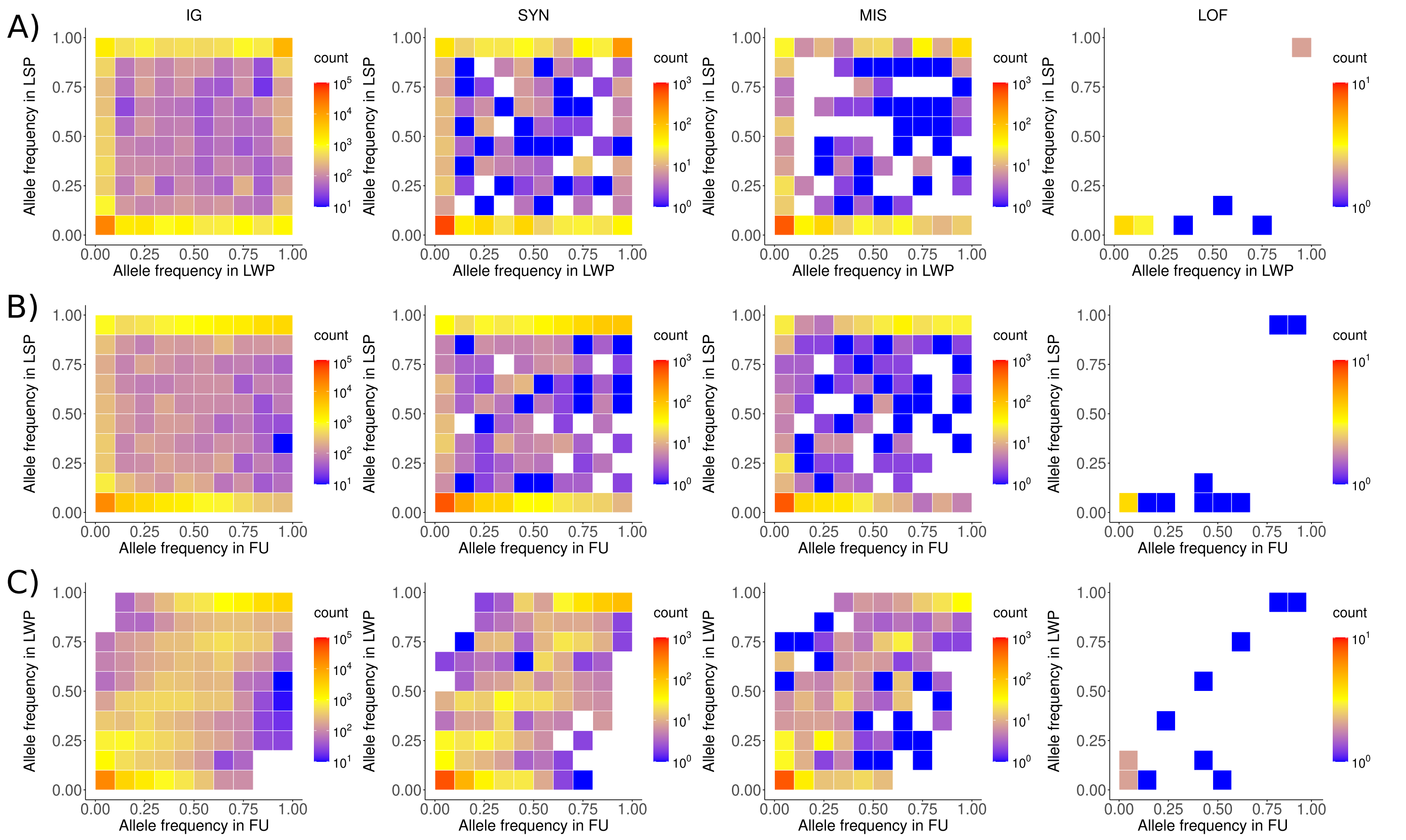


Figure S36: Two-dimensional Site Frequency Spectrum (SFS) for intergenic (IG), Synonymous (SYN), Non-synonymous (MIS) and High impact (LOF) variants on the Z chromosome. **(A)** Joint SFS for each category between LSP (15 males individuals) and LWP (12 males individuals). **(B)** Joint SFS for each category between LSP (15 males individuals) and FU (10 males individuals). **(C)** Joint SFS for each category between LWP (12 males individuals) and FU (10 males individuals).
